# An open-access T-BAS phylogeny for Emerging *Phytophthora* species

**DOI:** 10.1101/2022.08.06.503053

**Authors:** Allison Coomber, Amanda Saville, Ignazio Carbone, Jean Beagle Ristaino

## Abstract

*Phytophthora* species cause severe diseases on food, forest, and ornamental crops. Since the genus was described in 1875, it has expanded to comprise over 190 formally described species. There is a need for an open access bioinformatic tool that centralizes diverse streams of sequence data and metadata to facilitate research and identification of *Phytophthora* species. We used the Tree-Based Alignment Selector Toolkit (T-BAS) to develop a phylogeny of 192 formally described species and 33 informal taxa in the genus *Phytophthora* using sequences of eight nuclear genes. The phylogenetic tree was inferred using the RAxML maximum likelihood method. A search engine was also developed to identify genotypes of *P. infestans* based on genetic distance to known lineages. The T-BAS tool provides a visualization framework allowing users to place unknown isolates on a curated phylogeny of all *Phytophthora* species. Critically, this resource can be updated in real-time to keep pace with new species descriptions. The tool contains metadata such as clade, host species, substrate, sexual characteristics, distribution, and reference literature, which can be visualized on the tree and downloaded for other uses. This phylogenetic resource will allow data sharing among research groups and the database will enable the global *Phytophthora* community to upload sequences and determine the phylogenetic placement of an isolate within the larger phylogeny and to download sequence data and metadata. The database will be curated by a community of *Phytophthora* researchers and housed on the T-BAS web portal in the Center for Integrated Fungal Research at NC State. The T-BAS web tool can be leveraged to create similar metadata enhanced phylogenies for diverse populations of pathogens.

## Introduction

*Phytophthora* is a genus of destructive, oomycete plant pathogens that cause devastating plant diseases on food crops, ornamentals, and in forest and riparian ecosystems (1). *Phytophthora infestans* (Mont.) de Bary was the first species in the genus described, and was the biological agent responsible for the Irish potato famine in the 1840s (2, 3). *Phytophthora infestans* causes late blight on tomato (*Solanum lycopersicum*) and potato (*Solanum tuberosum*) and remains an important threat to crop production globally (4).Other *Phytophthora* species also threaten agricultural production systems and natural ecosystems (1, 5, 6). Important examples include *P. ramorum,* responsible for the disease known as sudden oak death and *P. cinnamomi*, a generalist species with a wide host range (7–9). In western Africa, *P. megakarya* is responsible for black pod disease of cacao and threatens cacao production (10). Globally, other *Phytophthora* species cause destructive plant diseases in many other plant species (11, 12).

Diverse new species of *Phytophthora* are regularly being described from croplands, forests, and water ecosystems around the world (6, 13, 14). To date, more than 192 species have been formally described, most of them in the last 15 years (1, 6, 12). More species are expected to be discovered or described as surveys of water, riparian buffer plants, and forest ecosystems continue (9).

Historically, morphological characteristics have been used to identify *Phytophthora* species (14). Morphological groupings and dichotomous keys have been useful tools in identifying *Phytophthora* species based on observed morphological characteristics (14–18). A Lucid Key for identification of common *Phytophthora* species has also been developed (19). This resource incorporated both morphological and molecular characteristics to aid in species identification (19, 20). However, in the past 10 years, the number of new species discovered has expanded greatly, resulting in many recently described species which are not included in this resource.

Species identification has relied more on molecular identification methods such as single gene or multilocus sequencing and genotyping by sequencing (14, 21).This has also enabled the production of more robust phylogenies based on sequence similarity (14, 22, 23). In 2000, a phylogeny of 50 *Phytophthora* species was developed by Cooke and colleagues based on *ITS* sequence data (22). Since then, expanded phylogenies with additional species and loci have been developed as both the genus and sequencing resources have grown (12, 23–25). Most recently, Yang and colleagues presented a robust phylogeny for the genus, including many newly described species and isolates that have since been or may later be described as individual species (12). They and others have used multilocus genotyping in *Phytophthora* to differentiate species (12, 26). Although multilocus sequencing generates robust phylogenies, the morphological and biological information of the associated species is still of high importance to researchers and is often left out of resulting phylogenies.

The current phylogenetic system and curated knowledge on the *Phytophthora* genus is disjointed, with molecular phylogenies and biological information being presented in disparate resources. Several databases with biological and sequence information on *Phytophthora* are available, but those systems do not integrate new biological information from species descriptions, nor do they allow for expandable phylogenies curated by the research community (27–29). There is a need to connect available *Phytophthora* data into a centralized resource to facilitate species identification and to study the evolution of species within the genus. The software toolkit “T-BAS” has been developed for the integration of phylogenetic placement and visualization of biological metadata (30, 31). This resource has been used effectively for development and presentation of fungal phylogenies, especially for the Ascomycota (32, 33). The effectiveness of T-BAS for other fungal systems, including plant pathogens, indicates its potential utility for making a more centralized resource for *Phytophthora* identification data. We harnessed T-BAS to build a living phylogeny of *Phytophthora*, incorporating sequence data and metadata for most of the recently described species. Most importantly, the live phylogeny format allows for the rapid, curated placement of new species and taxa as they are described.

*Phytophthora infestans* consists of multiple lineages that scientists differentiate to inform research and management practices (34). *Phytophthora infestans* populations are dominated by clonal lineages which have varying agronomic traits, such as fungicide sensitivity and host specificity (35). Tracking the spread and prevalence of these lineages is critical to pathogen management in both the United States and Europe. The temporal and geographic distribution of lineages is monitored by researchers on USABlight and Euroblight in these two regions, respectively (4, 34, 36–39). The current standard practice to identify a lineage of *P. infestans* involves amplification of 12 microsatellite (SSR) markers (34, 39, 40). Once the markers are amplified, they are typed according to the number and size of alleles at the 12 loci using the protocol outlined by Li and colleagues (40). The system in place for identification of *P. infestans* lineages relies on the expertise of a few well-trained researchers and there is no centralized queryable database of genotypes. One online tool, Phytophthora-ID: Genotype-ID has been developed to identify *P. infestans* genotypes based on SSR markers, but not all recent global genotypes are incorporated into that database (28). Like species identification within the genus, *Phytophthora infestans* lineage identification would benefit from a centralized open resource with molecular and biological data integrated and curated by the late blight research community.

Given the disparate data sets and expansion in reports on new species of *Phytophthora,* a more centralized system for curating sequence data and inferring robust phylogenies is needed. The primary objectives of this work were to: 1) Develop an open T-BAS *Phytophthora* phylogeny by synthesizing sequence data, biological trait data, and metadata from various published sources; 2) Make this phylogeny comprehensive and easily updatable by the research community to keep pace with discovery of new *Phytophthora* species; 3) Develop a search engine for *P. infestans* SSR data for the identification of genotypes.

## Methods

### Sequence Data Collection

Publicly available sequence data for described *Phytophthora* species were downloaded for nine loci (*28S*, *60SL10*, *Btub*, *EF1α*, *Enl*, *HSP90*, *TigA, ITS,* and *CoxI*) from GenBank, drawing on previous phylogenetic works and species descriptions for new species (Supplemental Table 1) (12). All of these loci are nuclear with the exception of *CoxI* locus.

Building on this dataset, we sequenced seven of these loci (*28S*, *60SL10*, *Btub*, *EF1α*, *Enl*, *HSP90*, and *TigA*) for two additional species of *Phytophthora* which were recently described, *Phytophthora acaciae* and *Phytophthora betacei* (Supplemental Table 1) (41, 42). *Phytophthora acaciae* was described as a species in Brazil in 2019 and infects black wattle (*Acacia mearnsii*) (41). *Phytophthora betacei* was described in Colombia in 2018 and is a pathogen of tree tomato (*Solanum betaceum*), an important crop species in that area (42). This species was previously classified as *P. andina* lineage EC-3 in earlier published work (43, 44).

Samples of *P. acaciae* were obtained from Dauri Tessmen, Universidade Estadual de Maringá, Parana, Brazil. Samples of *P. betacei* were obtained from Sylvia Restrepo, University of Los Andes, Colombia. DNA extraction was performed using the CTAB method as described previously (45). The primers used to amplify each of the seven loci were from previous published work (Supplemental Table 2) (12, 22, 24–26, 46, 47). PCR reactions were done in 50 μL volumes. Each 50 μL reaction contained 5 μL of 10X PCR buffer (Genesee, SanDiego, CA), 2.5 μL dNTPs (2 mM per nucleotide), 2 μL each 10 μM forward and reverse primer, 1.8 μL MgCl_2_ (50mg/mL), 0.25 μL BSA (20mg/mL), 0.2 μL Taq (5U/μL) (Genesee, SanDiego, CA), with the remainder to 49 μL as dd H_2_O. The final 1 μL consisted of sample DNA. Thermal cycling protocol consisted of 94°C for 5 minutes; then cycles of 94°C for 2 minutes, an annealing step specific to the primers, and 72°C for 2 minutes; followed by a final extension period at 72°C for 2 minutes (Supplemental Table 2). For the primers amplifying the TigA and HS90 loci, the annealing step was for 30 seconds with temperatures ranging from 64°C to 62°C, and 12 cycles completed at each temperature. For the other regions, the annealing step was for 30 seconds with temperatures ranging from 60°C to 53°C, and 3 cycles completed at each temperature plus an additional fifteen cycles with an annealing temperature of 53°C. Temperature reductions in the annealing step over time, known as touchdown PCR, allowed for amplification of multiple loci in one thermocycler protocol. Detailed descriptions of the primers, their optimum annealing temperatures, and the sources they were adapted from can be found in Supplemental Table 2. Amplified fragments from the PCR reactions which were expected to contain the locus of interest were sequenced using Sanger sequencing at the Genomic Sciences Laboratory at North Carolina State University.

In total, sequence data was collected for 192 *Phytophthora* species, 30 informally described *Phytophthora* taxa, and 3 outgroups from related oomycete genera (Supplemental Table 1). Compared to the most recent phylogeny, this represents the addition of 50 new taxa (12).

### Phylogenetic inference and visualization

Phylogenetic trees were inferred with 1000 bootstrap replicates under the GTRGAMMA model using RAxML version 8 via the CIPRES REST API implemented in the DeCIFR toolkit (https://tools.decifr.hpc.ncsu.edu/denovo) (48, 49). In total, three sets of phylogenetic trees were generated. First, a phylogenetic tree based on the eight nuclear loci was constructed after excluding taxa with sequence data for fewer than three loci. An additional phylogenetic tree consisting of only the mitochondrial *CoxI* locus was inferred for species which had sequence data available at this locus. Lastly, independent trees for each locus were also inferred separately. Trees for all nine loci were compared using the Hypha package module of Mesquite v3.51 implemented in the DeCIFR toolkit (https://tools.decifr.hpc.ncsu.edu/trees2hypha) (50, 51). The tree based on the eight nuclear loci was selected for further analysis.

The resulting *Phytophthora* genus tree was uploaded into the Tree-Based Alignment Selector Toolkit (T-BAS version 2.3) developed by Ignazio Carbone and colleagues (30). The T-BAS system was chosen because it has a phylogeny-based placement feature which allows incorporation of new taxa into an existing phylogenetic tree. Metadata for the taxa in the tree were collected using a custom Python script for web scraping with the package BeautifulSoup as well as manual data collection (52). A portion of the metadata for some species was retrieved from IDPhy, a published, open-access *Phytophthora* database developed and curated by Gloria Abad and colleagues (27). Metadata for some traits was summarized into a smaller number of categories to facilitate visualization. For example, host range was characterized as ‘specific’ or ‘broad’ following the categories presented in (9). Each species was manually encoded to fit into one of these categories based on the number of host plants and their relatedness. *Phytophthora* species with few, closely related hosts were placed in the specific category, while those with a larger number of hosts or taxonomically varied hosts were considered broad. For species where host information was lacking, no host range category was determined. A full list of specific hosts was also retained in the metadata. Similarly, pathogen substrate was categorized into soil, water, and/or foliar (27). Above ground disease symptoms were categorized as foliar, while below ground infectivity was categorized as soil. Specific substrate information was also retained in the metadata. *Phytophthora* species with unknown substrate were left blank. Sexual information was encoded as homothallic or heterothallic, where available. All distribution data were both retained in full and summarized into a list of continents for each species. The visualization capabilities of T-BAS were harnessed to overlay this metadata set with the phylogenetic tree inferred from the sequence data. Users have the option to download single and multilocus DNA sequence alignments, metadata, and Newick-formatted trees for focal clades of interest.

### Phylogeny validation

To test real-time phylogenetic placement onto the tree, sequence data for the two newly described *Phytophthora* species sequenced in this study (*P. betacei* and *P. acaciae*) were added to the phylogeny. The new taxa were inserted in the tree using the phylogeny-based evolutionary placement algorithm (EPA) implemented in T-BAS (30, 53, 54). Additionally, independently identified species of *Phytophthora* collected by collaborator Inga Meadows from nursery crop plants were placed in the multilocus tree using the *Phytophthora* reference tree as a backbone constraint tree and 1,000 bootstrap searches in RAxML. A tutorial and practice FASTA files for placement of two “unknown” *Phytophthora* species can be found on the project GitHub repository (https://github.com/allisoncoomber/phytophthora_tbas).

### *P. infestans* lineage classifier development

A dataset of approximately 2200 *Phytophthora infestans* isolates of known lineages and their corresponding microsatellite genotypes were obtained from over thirty years of collecting by Jean Beagle Ristaino (2, 4, 55). Using R, an applet was developed to compare newly typed isolates of *P. infestans* to this dataset of lineages (56). For genotype identification, unknown isolates were compared to a reference genotype using Bruvo’s genetic distance algorithm, implemented by the R package Poppr (57, 58). Bruvo’s distance was selected for comparing unknown genotypes to the reference because it compares the relative genetic closeness of samples regardless of the ploidy (57). *Phytophthora infestans* has been documented to have variations in ploidy from diploid to tetraploid, with triploid clonal lineages being common (59). Variations in ploidy can impact results of other common genetic distance methods, so Bruvo’s distance is a logical choice for this species (57). A newly typed isolate was considered to belong to a genotype if it was within a threshold genetic distance of other isolates in that genotype (described below).

A user interface was developed for the applet using the R Shiny package (60). The SSR genotype classifier is linked to the USABlight.org website under the “Identify an SSR genotype” page. A tutorial describing how to use the classifier and example files can be found in Supplemental File 1.

### *P. infestans* lineage classifier testing

A threshold genetic distance to differentiate lineages was estimated using the cutoff-predictor function from Poppr (58). A histogram of Bruvo’s values for all pairwise comparisons also provided visual confirmation of the threshold value (Supplemental Figure 1). Biologically, this threshold distance approximates the distinction between clonally and sexually related pairs, allowing the classifier to distinguish clonal genotypes.

The applet was also tested using a minimum representative set of distinct reference genotypes representing global populations of *P. infestans* to investigate which genotypes were similar and likely to be confused for each other. In this case, the reference set was developed by selecting the most common microsatellite profile for each genotype in the dataset. The remainder of isolates were then used as the tester dataset. These parameters produced a similar cutoff threshold to that estimated by Poppr.

To test the sensitivity and specificity of the classifier, the reference dataset was split into five randomly selected, non-overlapping samples. Each sample was classified using the remainder of the reference dataset (i.e., those not included in the sample). For some lineages which had only one or a few representative isolates, sensitivity and specificity could not be estimated.

## Results

### Live Phylogeny of *Phytophthora*

A total of 192 formally described *Phytophthora* species and 33 affiliated taxa were included in the genus phylogeny (Figure 1, Supplementary Table 1). The maximum likelihood phylogeny was inferred with RAxML and includes 8 concatenated nuclear loci (*28S, 60SL10, Btub, EF1α, ENL, HS90, ITS,* and *TigA*). The tree was rooted with three outgroups from related oomycete genera (*Halophytophthora fluviatilis, Phytopythium vexans,* and *Elongisporangium undulatum*) (Figure 2). A full list of included species and accession numbers for each locus can be found in Supplemental Table 1. There was strong support for the outgroups being separate and distinct from the *Phytophthora* genus, with the *Phytophthora* genus appearing as a monophyletic clade (Figure 2).

**Figure 1.**
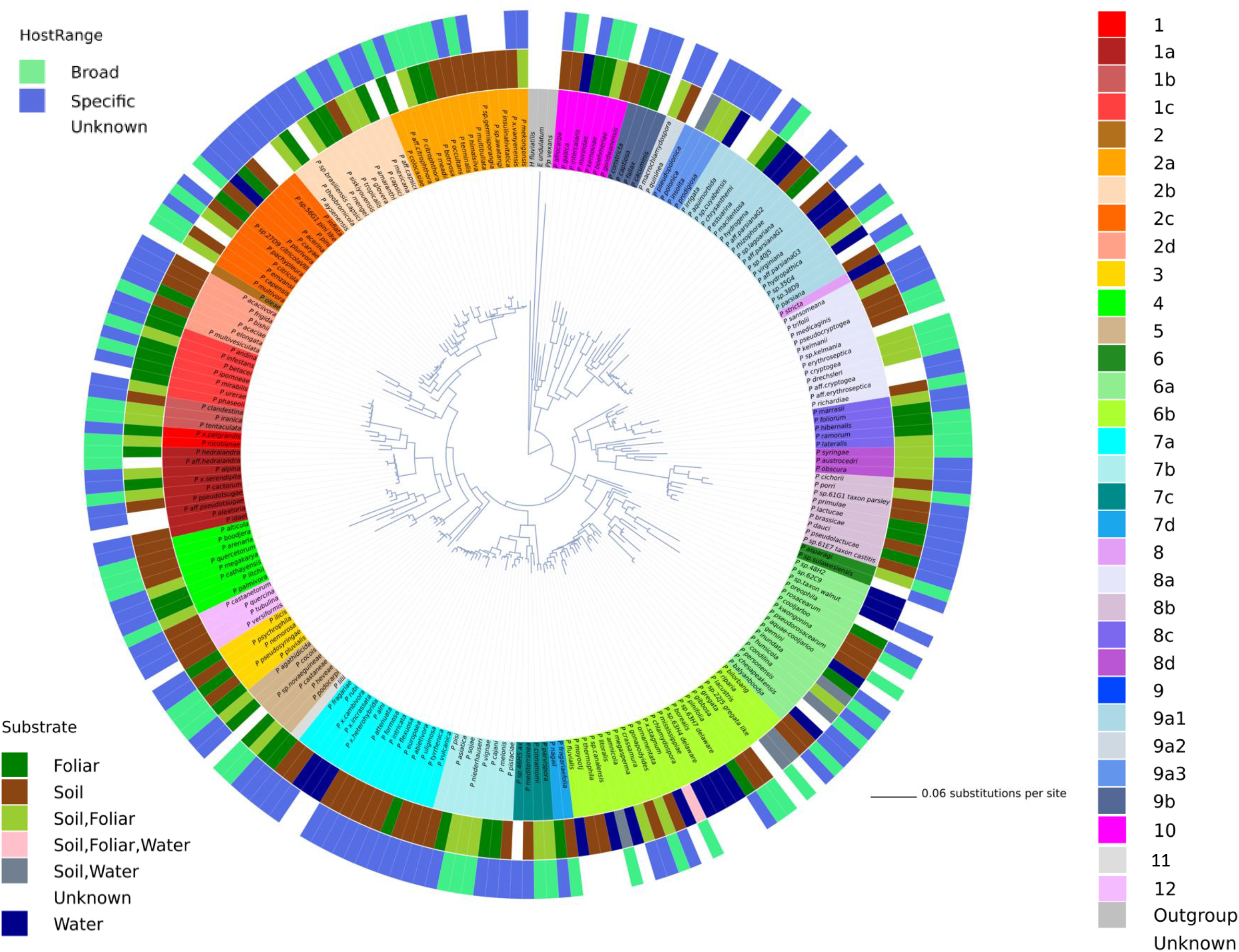
Radial phylogeny of the genus *Phytophthora* inferred with maximum likelihood and 1,000 bootstrap replicates for an alignment of 8 concatenated nuclear genes. Coloring on the inner ring indicates clade. Colors on the middle ring indicate substrate. Colors on the outer ring indicate host range (broad or specific). Branch lengths are drawn proportional to number of substitutions.

**Figure 2.**
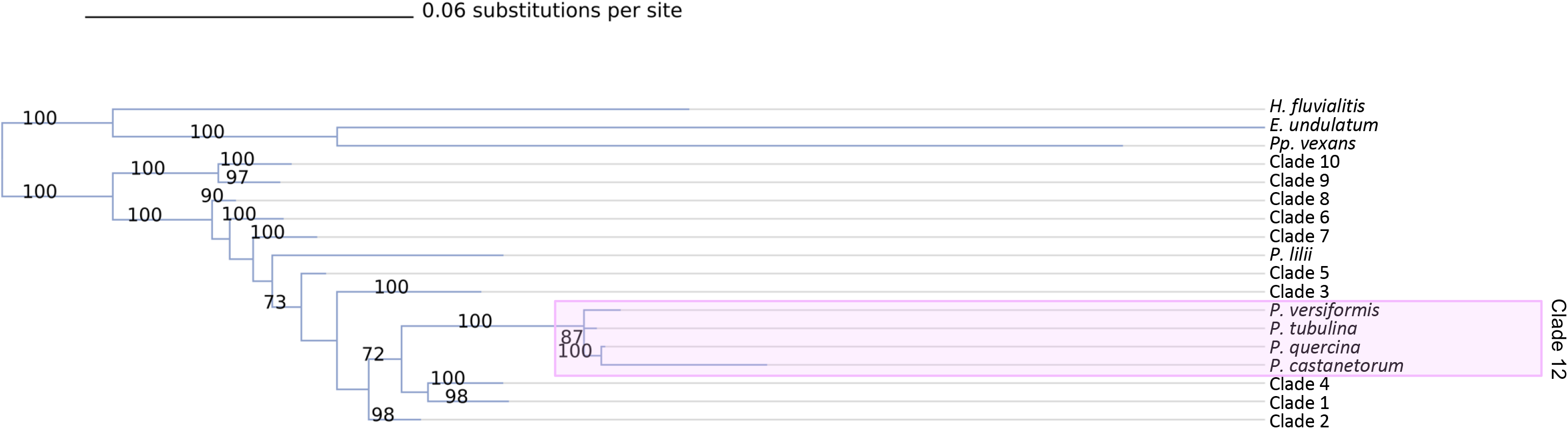
Collapsed phylogenetic tree of the genus *Phytophthora* showing the relationships between the clades, outgroups, and species that do not fit into the conventional ten clade system. New clade 12 is highlighted in pink. Phylogeny is inferred using maximum likelihood and 1,000 bootstrap replicates for an alignment of 8 concatenated nuclear loci. Bootstrap values (in percent) are shown above each branch. Branch lengths are drawn to scale.

The results of this phylogeny were largely in agreement with what has been previously presented. There was strong evidence for the ten canonical clades, and the relationships between clades were mostly in agreement with previous studies (Figure 2). Fifty species or closely affiliated taxa were placed in our tree that were not reported in previously published phylogenies (Table 1, Supplemental Figure 2). Placement of these taxa was overall in agreement with their previous descriptions (references for each in Table 1). Several taxa which were previously given only a general clade were resolved to the subclade level by this study (Table 1). For example, *P. prodigiosa, P. pseudopolonica, P. rhizophorae, P. estuarina,* and *P. cacuminis* were all previously described as belonging to clade 9. Here, we resolve them to the subclade or subclade cluster level as 9a3, 9a3, 9a1, 9a1, and 9b respectively. Additional subclade distinctions and their previous clade classifications are listed in Table 1. Notably, the addition of several new species closely related to *P. quercina - P. versiformis, P. tubulina,* and *P. castanetorum –* allowed for a more detailed resolution of clade 12. This clade was previously referred to as 3b.

**Table 1.**
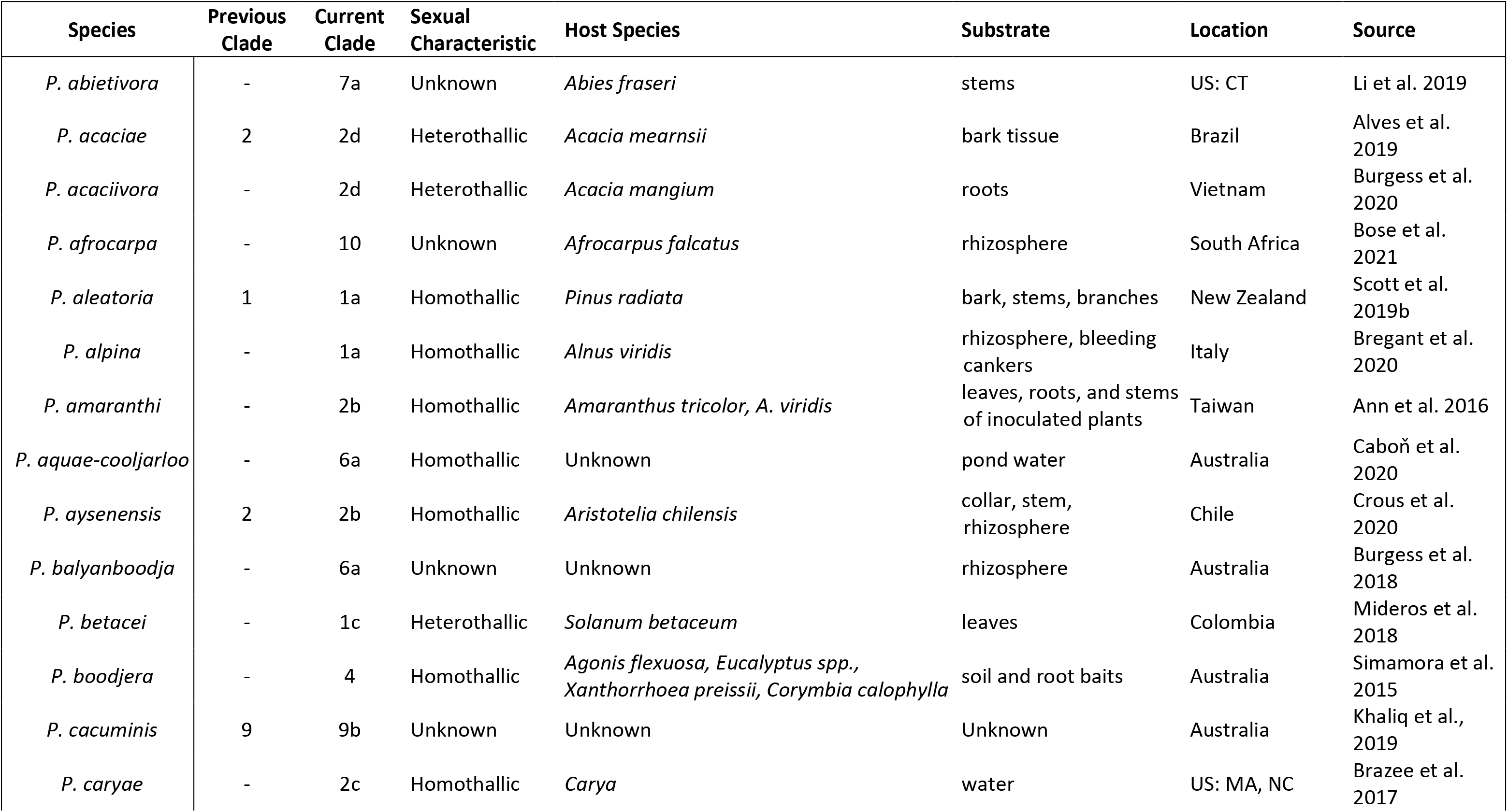
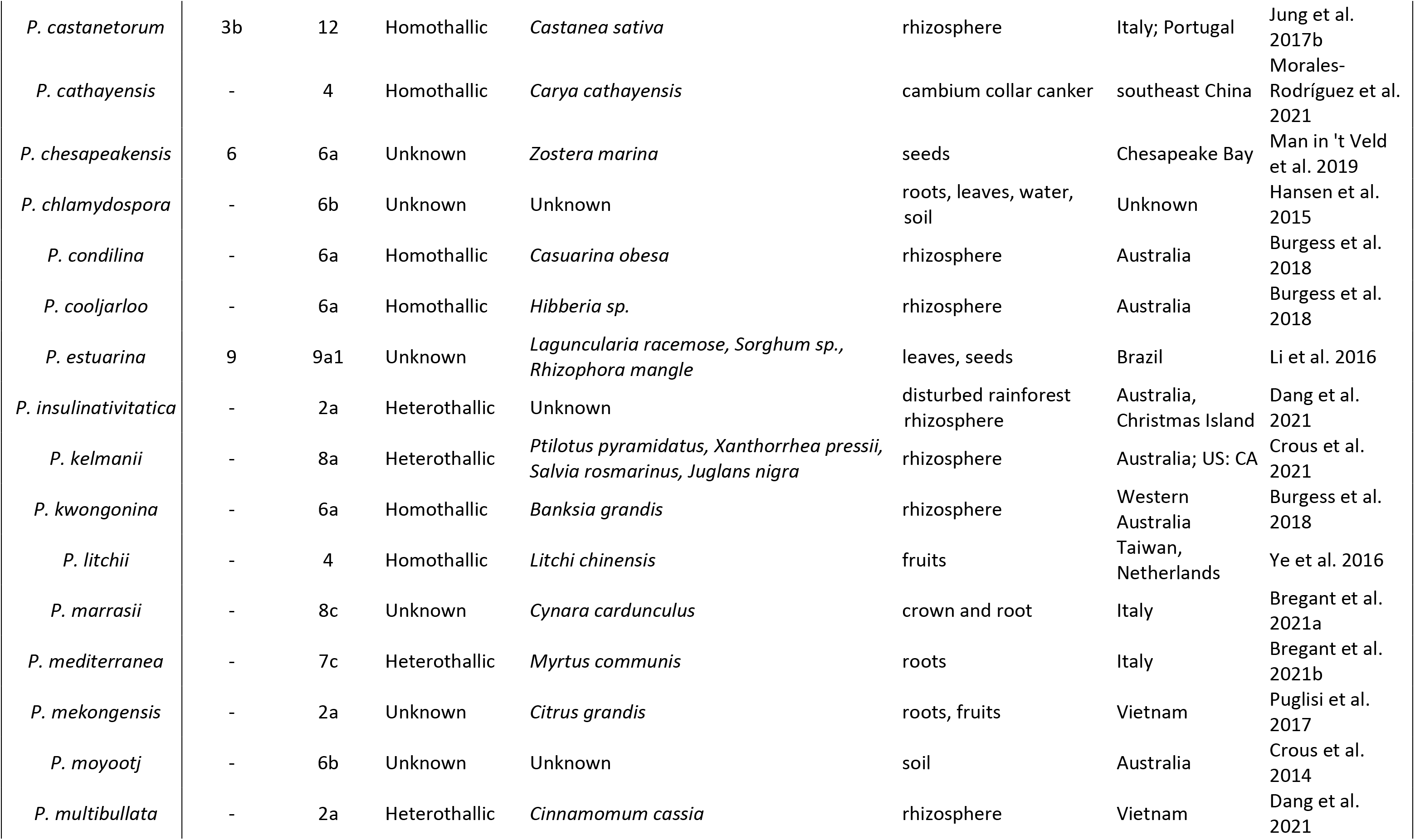
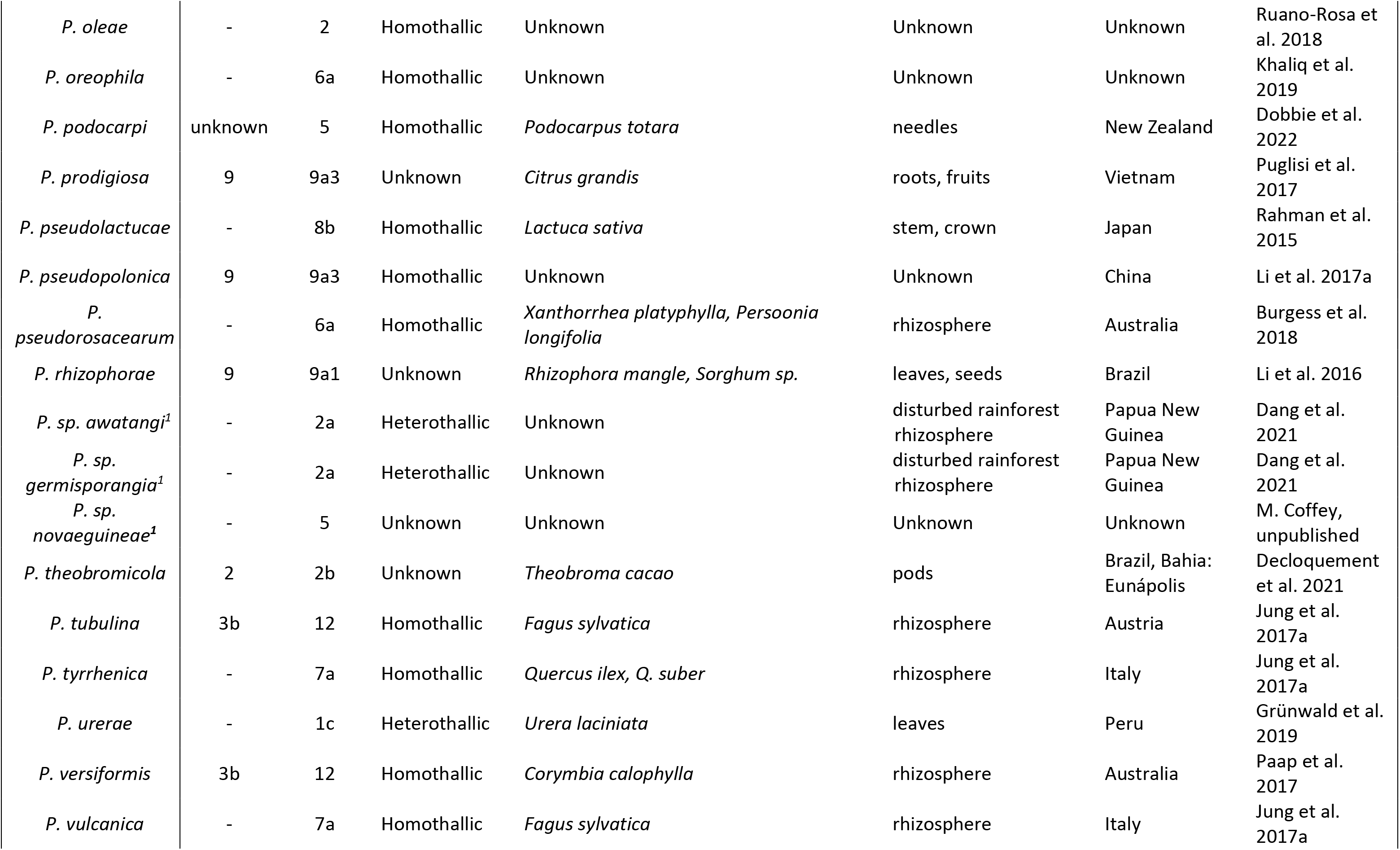
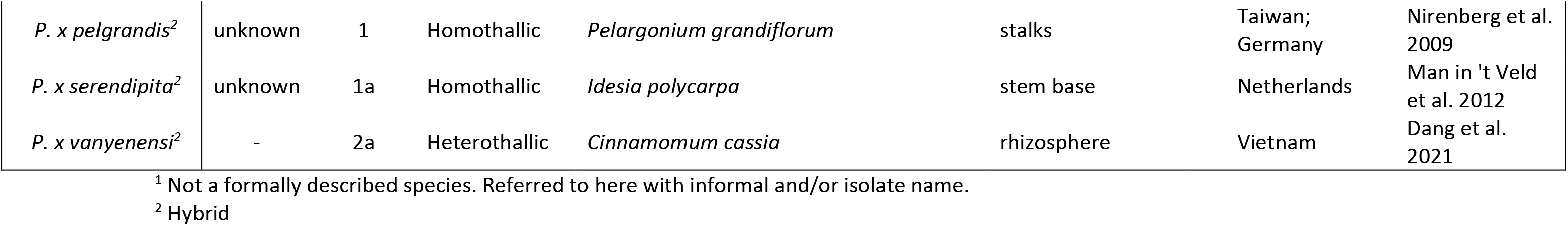
New taxa added to the genus *Phytophthora* in this phylogeny along with a summary of their associated metadata and species description paper.

The topology of the *CoxI* mitochondrial tree differed from the nuclear derived trees, as is expected because mitochondria are uniparentally inherited in *Phytophthora* (Supplemental Figure 3, Supplemental Figure 4). Because of this distinction, the *CoxI* locus was included as a single locus tree, but not with the other nuclear loci in the multilocus tree.

Phylogeny-based placement of *P. acaciae* grouped it in clade 2d with sister group *P. bisheria* (Supplemental Figure 5). *Phytophthora betacei* was placed in clade 1c near both *P. infestans* and *P. andina* (Figure 3).

**Figure 3.**
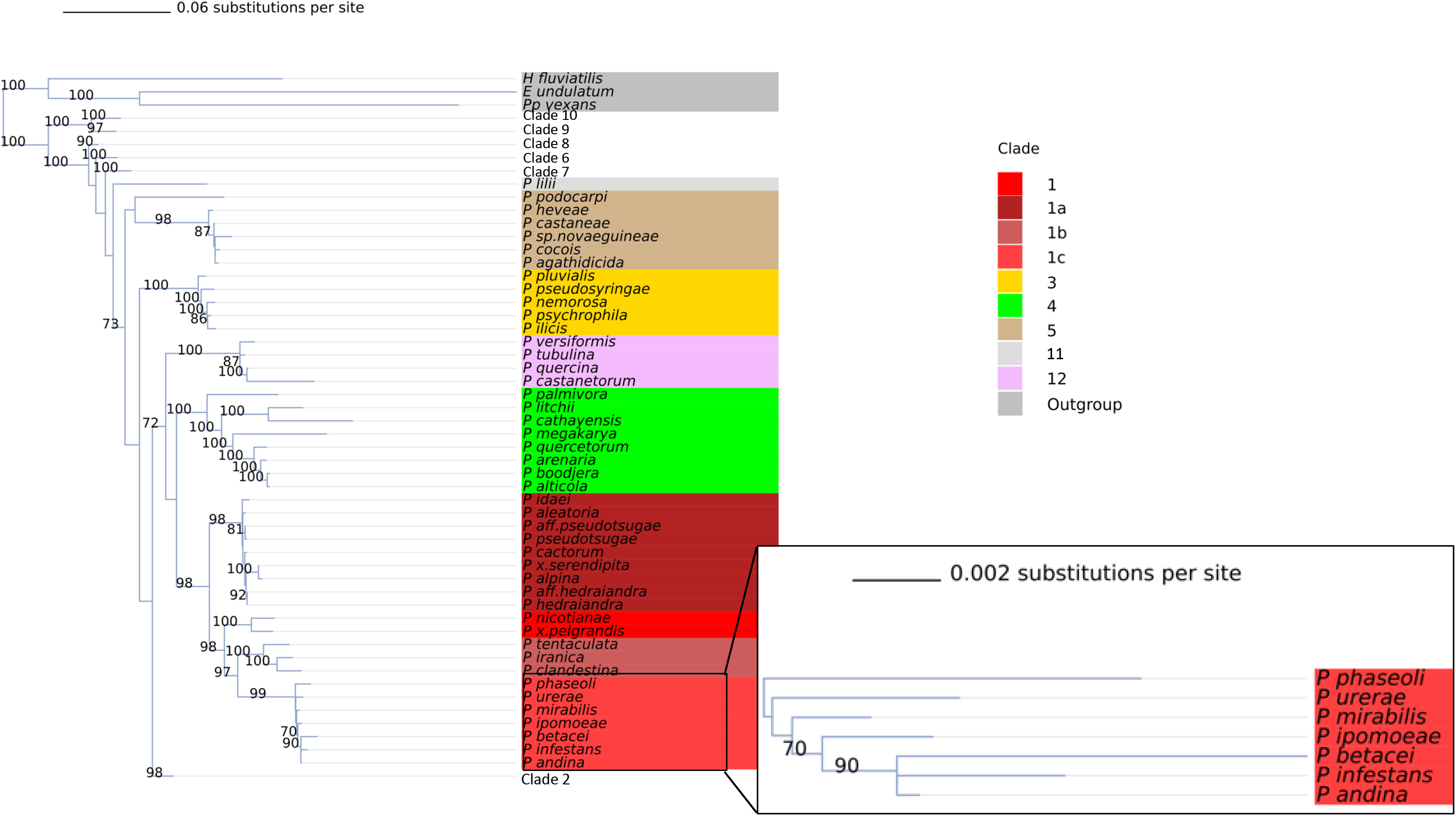
Collapsed phylogeny of the genus *Phytophthora* showing clades 1 (red), 3 (yellow), 4 (green), 5 (beige), and 12 (pink) in detail. Subclade values are shown as variations in color hue. Phylogeny is inferred using maximum likelihood and 1,000 bootstrap replicates for an alignment of 8 concatenated nuclear loci. Bootstrap values (in percent) are shown above each branch. Branch lengths are drawn proportional to number of substitutions.

Collaborator Inga Meadows tested the real-time phylogenetic placement tool using 80 *Phytophthora* isolates spanning seven species from previous sequencing work (*P. nicotianae, P. tropicalis, P. palmivora, P. drescherli, P. cryptogea, P. pseudocryptogea, and P. kelmania*). Phylogenetic placements matched previous identification, with the exception of eight isolates of *P. cryptogea* that were reclassified as *P. pseudocryptogea.* The eight isolates which were identified incorrectly were collected and identified before the description of *P. pseudocryptogea*, explaining the discrepancy (61).

#### Metadata

Metadata, such as sexual reproductive mode (heterothallic, homothallic, or unknown) were collated across the genus (Supplemental Figure 6). Since 2005 the number of homothallic species reported has exceeded the number of heterothallic species (Supplemental Figure 7). In general, the ecological niche where *Phytophthora* are isolated (soil, water, or foliar) is diverse across the genus with additional species reported in each substrate over time (Figure 4a). Over the past 10 years, new *Phytophthora* species have been described from soilborne habitats more often than water or foliar substrates (Figure 4a). However, clusters of multiple water-isolated species are present in Clades 6, 7, and 9 (Figure 5). In recent years, there has been a significant expansion in the number of surveys for water-borne *Phytophthora* species.

**Figure 4.**
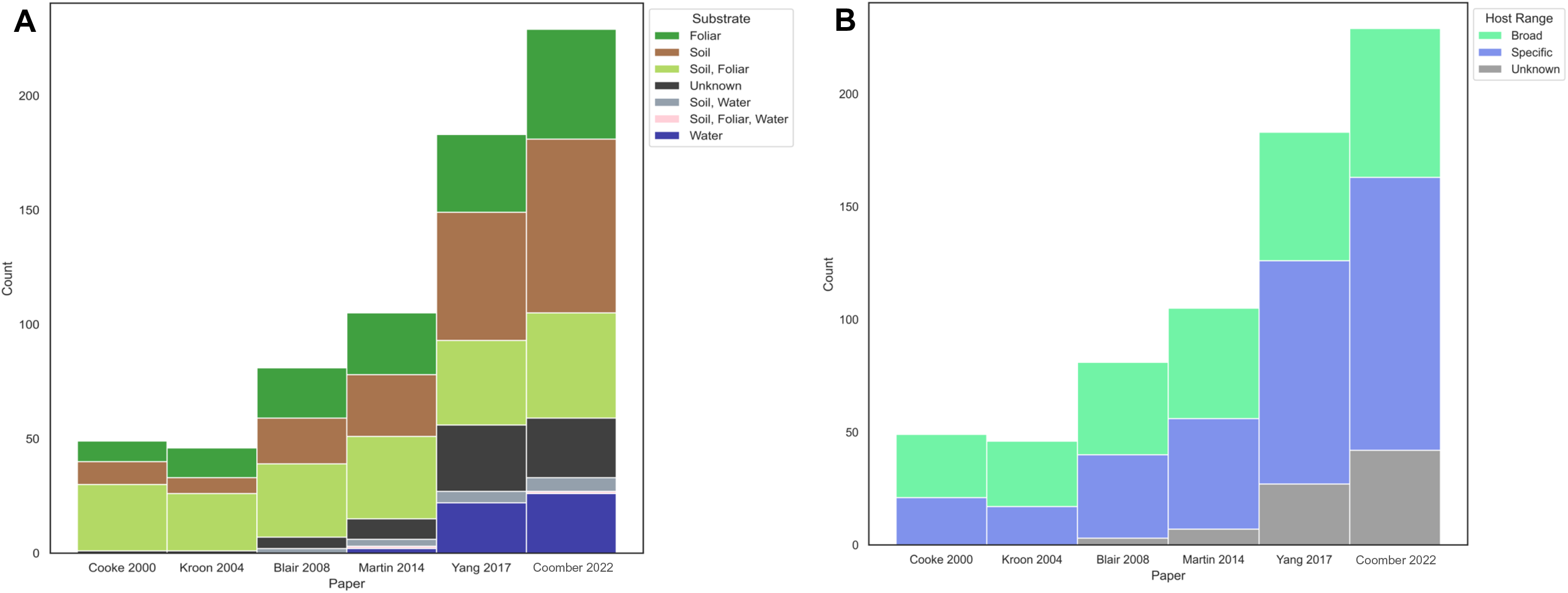
A. Histogram showing the number of *Phytophthora* species in each substrate category in all major phylogenies published since 2000. B. Histogram showing the number of *Phytophthora* species in each host range category in all major phylogenies published since 2000. Host specificity was characterized as “broad” or “specific” based on the number of hosts the pathogen has as well as the relatedness of the hosts.

**Figure 5.**
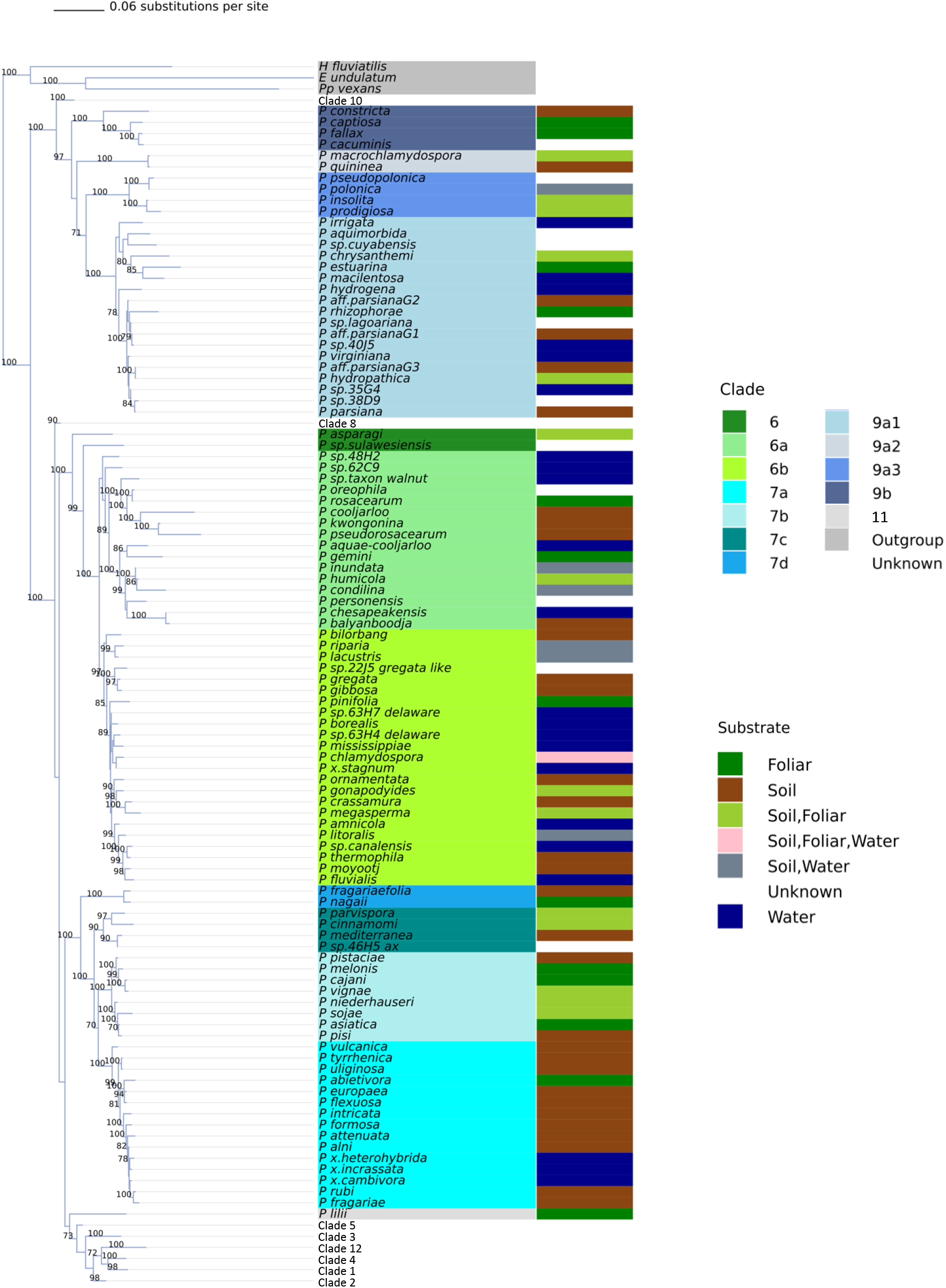
Collapsed phylogeny of the genus *Phytophthora* showing 6 (green), 7 (teal), and 9 (blue) in detail. Subclade values are shown as variations in color hue. Phylogeny is inferred using maximum likelihood and 1,000 bootstrap replicates for an alignment of 8 concatenated nuclear loci. Bootstrap values (in percent) are shown for each branch. Branch lengths are drawn proportional to number of substitutions.

The level of host specificity also varies widely across the genus. Host specificity was characterized as “broad” or “specific” based on the number of hosts the pathogen has as well as the relatedness of the hosts. Species with a broad host range were clustered in Clades 1, 2, and 7 (Figure 1). However, the ability to cause disease on a wide variety of hosts is also generally widespread, being found at least once in all clades (Figure 1, Figure 4b).

### *Phytophthora infestans* SSR Classifier

In order to differentiate clonal lineages of *Phytophthora infestans* we estimated a threshold genetic distance to separate lineages. This threshold genetic distance was estimated as 0.099 by Poppr’s cutoff prediction function. A histogram of all pairwise comparisons of Bruvo’s genetic distance for the included lineages was bimodal, indicating a population with mixed forms of sexual and asexual reproduction, as is characteristic of *P. infestans* (Supplemental Figure 1). Pairs with a Bruvo’s genetic distance below the threshold occurred in the first peak of the distribution and indicated clonal relationships. In other words, the pair consisted of clones of the same lineage or genotype. Above the threshold was another peak in the distribution of Bruvo’s distances. Pairs falling in this range of the distribution represent lineages which are distinctly different, likely as a result of sexual recombination. In other words, these are lineages that should not be considered the same genotype. This result visually confirmed the threshold genetic distance as estimated by Poppr’s cutoff prediction function. The SSR classifier was parameterized to use a cutoff threshold of a Bruvo’s distance of 0.099 in order to consider an unknown *P. infestans* isolate a match to a described lineage.

The reference dataset for classifying genotypes consisted of 2,176 isolates of *P. infestans* for which all 12 microsatellite loci were genotyped. The overall accuracy of the classifier was approximately 0.98, with an unweighted Kappa statistic of 0.977. Various summary statistics were also calculated for each individual class (lineage, n=36) that isolates could be placed into (Supplemental Table 3). Some of the isolates included in the reference dataset had one or only a few representative genotypes. For these isolates, accuracy, precision, and other summary statistics were not calculated because there were no closely matching isolates. Many of these isolates are no longer in circulation (most of the US lineages except US-23, US-8, and US-11) or were transient. Importantly, the SSR classifier was able to identify the US genotypes from the recent past that are currently circulating (Supplemental Table 3). As collection of isolates continues the classifier will continue to grow more robust with the increase in reference data. A list of all genotypes used to develop the classifier and their SSR profiles can be found in Supplemental Table 4. This includes several genotypes for which there were too few isolates to perform statistical testing.

To evaluate the performance of the *P. infestans* classifier with incomplete SSR genotype data, a testing dataset was developed that included genotyped lineages which had at least 7 but no more than 11 SSR loci genotyped. Many of these isolates were from historic samples of the FAM-1 lineage. For incomplete data, overall accuracy was 0.968 and the Kappa statistic was 0.954.

Out of all the genotypes in the dataset, the SSR classifier was unable to distinguish between some genotypes at the decided threshold. Most of these multi-genotype groups contain one genotype which is still in circulation, as well as some genotypes which are no longer found. In this case, the genotype which is still in circulation is returned. If the multi-genotype group contains multiple genotypes still in circulation, all those genotypes are returned after querying the database. For example, genotypes US-6 and US-7 could not be distinguished from US-11 by our classifier. However, US-6 and US-7 were transient lineages which sexually recombined to produce US-11. Of these three lineages, only US-11 has persisted and so this is the lineage name that is returned by the classifier.

## Discussion

### Genus Tree Placements

The *Phytophthora* tree presented here is designed to serve as a starting point for a curated, living phylogeny of the genus that may include species that are the direct ancestors of other contemporary taxa. Including at least three nuclear loci for well described *Phytophthora* taxa and 50 new taxa has allowed for the resolution of conflicts within the genus phylogeny. For example, *P. quercina* and related species have previously been placed in clade 3b in an ITS phylogeny, but the inclusion of other nuclear loci shows that *P. quercina, P. versiformis, P. tubulina,* and *P. castanetorum* are not monophyletic with the rest of clade 3 (Figure 3, Table 1) (27). This was also observed in the species description paper of *P. tubulina* and *P. castanetorum*, which reclassified these species as a part of a new clade 12 (13). The phylogeny presented here supports this conclusion, with *P. quercina, P. versiformis, P. tubulina,* and *P. castanetorum* forming a unique monophyletic clade referred to as clade 12. This clade is separate and distinct from the rest of clade 3, forming a sister group to clades 1 and 4 (Figure 3). This finding resolves low confidence in the resolution of Clade 3 reported in previous studies by including at least three nuclear loci for multiple new species (12, 27).

In their 2017 *Phytophthora* phylogeny, Yang and colleagues subdivided clade 9 into clades 9a and 9b, where 9b is monophyletic and 9a consists of three monophyletic subclades, 9a1, 9a2, 9a3 (12). The addition of new species in our work was congruous with this subdivision of clade 9 (Figure 5). Several species newly included in this phylogeny were previously characterized as only “clade 9” but here we find support for a specific subclade (Figure 5, Table 1). *P. cacuminis* was strongly supported as part of Clade 9b (62). *P. pseudopolonica*, which was previously classified as clade 9, grouped closely with *P. polonica* in clade 9a3 (63). New species *P. prodigiosa* also clustered in clade 9a3, despite being classified as clade 9b by previous ITS-based phylogenies (27, 64). *Phytophthora estuarina, P. sp. lagoariana, P. sp. cuyabensis,* and *P. rhizophorae* were all supported as members of clade 9a1 (65). In general, the discovery of new species in clade 9 supports the current grouping of clade 9 into the subclades proposed by Yang et al. 2017 (Figure 5).

Clade placement was also confirmed for two species that had additional loci sequenced for this study, *P. acaciae* and *P. betacei. Phytophthora acaciae* was placed in clade 2d with close relatives *P. bisheria* and *P. frigida* which is in agreement with previous work on the species (Supplemental Figure 5) (41). *Phytophthora betacei* was confirmed as a member of clade 1c as previously reported (Figure 3) (42).

Though not included in the final genus tree, collected data for *CoxI* and the inferred tree are available for further analysis (Supplemental Figure 3, Supplemental Table 1). The presence of unexpectedly close relationships between *CoxI* loci for some species warrants further investigation into potential hybrids. For example, in the tree based on eight nuclear loci *P. andina* is considered to be a sister group to *P. infestans* with weak support (18) (Figure 3). However, in the tree inferred from the *CoxI* mitochrondiral locus, *P. infestans* and *P. andina* have stronger support as sister groups (100) (Supplemental Figure 3). This indicates that *P. andina*’s mitochondrial genome is more closely related to that of *P. infestans* than the nuclear genome, a result which could be explained by *P. andina* being a hybrid of *P. infestans*, as has been previously reported (14, 23, 66, 67). More data, especially sequence data from mitochondrial loci, needs to be collected for some species to further investigate potential hybridizations. The discordance of the *CoxI* tree with the other single locus nuclear trees, as well as previously published trees, led us to exclude the *CoxI* locus from our final tree (Supplemental Figure 4). Conflict between multilocus analysis and *CoxI* placement was also observed by Yang and Hong in their 2018 evaluation of *Phytophthora* markers (26).

As mentioned above, there is a lack of resolution in the nuclear tree for the relationships between some species in the 1c clade. Although the separate mitochondrial tree can also be used to inform these relationships, more information is needed to confidently understand the linkages between 1c clade species. This clade is economically relevant as the home of *P. infestans* and has seen recent expansion with the additions of *P. urerae* and *P. betacei.* A more comprehensive, genomic phylogeny of this clade should be conducted to strengthen confidence in the phylogeny.

### Metadata

Incorporating the metadata into the phylogenetic visualization allows the user to see relationships in biological characteristics across the phylogeny. For example, the recent proliferation in the description of species which were isolated from water is shown to have occurred in multiple clades (Figure 4a, Figure 5). The primary niches of the *Phytophthora* species (broadly foliar, soil, or water) are widely distributed, with small clusters of the different substrates appearing in multiple clades (Figure 1). Several recently described species from waterways are clustered together in clade 6a and clade 9 (Figure 5). It appears that the ability to have success on different substrates has evolved multiple times in the genus.

Other than clade 5 and clade 3, which had entirely homothallic species, every clade had representatives of both homothallic and heterothallic species (Supplemental Figure 6). Both homothallism and heterothallism have been hypothesized to be the basal state of the genus (22, 25). Blair et al. 2008 agreed with Kroon and colleague’s assessment that homothallism was the basal state based on the lack of homothallism in both Clades 9 and 10, the basal clades (24). Here we show that heterothallic species are also present in these clades, *P. intercalaris* in clade 10 and *P. irrigata* and others in clade 9 (Supplemental Figure 6). The ancestral state of sexuality in *Phytophthora* is therefore unclear as more than 12 transitions from one state to another are shown in our most supported tree (Supplemental Figure 6). Thus reproductive strategy may be more transient than previously assumed. Others have shown that environmental stimuli including stress, aging and even fungicides can shift species such as *P. infestans* from heterothallism to homothallism (68, 69).

Host range from broad to specific is also a variable trait in *Phytophthora* (Figure 1). Again, this indicates relative ease in the evolutionary switch from wide to narrow host range strategy. Additionally, *Phytophthora* species have been clustered into two invasiveness groups based on host range (9). The categorization of each taxon into broad or specific host range could provide useful information on the potential invasiveness of a given species. More specific investigation on the phylogenetic relationships of specific hosts and species of *Phytophthora* might shed additional light on these relationships.

### Comparison to Other Databases

Several efforts have been made over the years to collect disparate information on *Phytophthora* species across the genus to facilitate identification, phylogenetics and evolutionary research. Notable examples include IDPhy, Phytophthora-ID for species and lineage identification, PhytophthoraDB and a lucid, dichotomous key for species identification (19, 20, 27–29). IDPhy presents a wealth of morphological information and other metadata for species identification and is a valuable resource (27). Phytophthora-ID enables phylogenetic placement of unknown strains based on only 2 genes - *CoxI* and *ITS* sequences, or SSR markers for *P. infestans* and *P. ramorum* lineages (28). However, Phytophthora-ID does not contain the full global set of known SSR lineages of *P. infestans* and many from the EuroBlight database are missing. Although both sites provide valuable resources, the need to synthesize genetic information with morphological characteristics and other descriptions is not addressed. Additionally, both sites rely on *ITS*-based trees, which are often not detailed enough to provide accurate species placements. Recently, the *Phytophthora* genus was included in the fourth edition of the “Genera of phytopathogenic fungi” series (GOPHY4) (70). This included a whole genus tree of 192 species. However, this tree used a limited number of loci (up to four) and combined both nuclear and mitochondrial loci in the same tree. *Phytophthora* species are known to undergo interspecies hybridizations, and so including loci with discordant inheritance patterns can result in misleading phylogenies (71–73). Our phylogeny includes more loci and focuses on nuclear loci to mitigate these dynamics. With a rapidly expanding genus, the need to update phylogenies and metadata descriptions to include newly described species and provisional species is critical. The contribution of the database presented here is to synthesize the wealth of sequence data and information that has been collected on *Phytophthora*, and to provide a more robust phylogenetic framework for the genus that can be updated at the pace of discovery by researchers in the community.

### Future directions

By incorporating the *Phytophthora* genus phylogeny into the T-BAS system, researchers will be able to add metadata, sequence data, and taxa as our understanding of the genus continues to expand. Metadata that is not presented here, such as taxonomic characteristics or phenotype data, could easily be added if of interest to the community. Adding a new species will require submission of the metadata and sequence data through the T-BAS website so the data can be curated before uploading. Maintaining a robust and rapidly updating phylogeny will allow the streamlining of scientific information on the genus to keep pace with the discoveries of new species and characteristics. Incorporating the metadata directly into the phylogeny may also provide new insights for *Phytophthora* researchers. Thus, a community of research experts involved in Phytophthora research have been designated to become validators for the genus level phylogeny and the *P. infestans* classifier.

Although the *Phytophthora* genus tree enables rapid phylogenetic placement of newly sequenced isolates, phylogenetic placement alone cannot be used for species level identification. In addition to sequencing the loci used in this study for phylogenetic placement in T-BAS, other methods such as morphology should be used to verify the species an isolate belongs to. Moving forward, many newly described species need to be more thoroughly studied. Several of the newly described species in this phylogeny have only a few sequenced loci, which weakens phylogenetic placement. For example, clade 9 remains difficult to fully resolve in part because only 3-4 of the loci are sequenced for several new species (*P. psuedopolonica, P. cacuminis, P. estuarina,* and *P. rhizophorae*). Furthermore, key metadata characteristics for many species, such as sexual strategy and hosts are missing. Going forward, we recommend at least four of the nuclear loci utilized here be sequenced for new species descriptions in addition to morphological requirements. Significant differences at these loci and in morphological traits is needed to warrant the description of a new species. These requirements will clarify the existing phylogeny and enable more robust phylogenetic placements of new species. Adding information into the phylogeny is facilitated by the T-BAS system, so new information about *Phytophthora* species can be readily available to the community, without the need to publish an updated description.

A review of the phylogeny also shows that newer species are often being described from natural ecosystems, such as riparian areas, as opposed to agricultural systems (Figure 1, Figure 5). Further exploration of these systems is warranted as it will likely reveal more species of *Phytophthora* that are yet to be discovered (9).

The utility of this tool for the plant-pathogenic *Phytophthora* genus serves as a proof of concept for other pathogen phylogenies. Enhancing pathogen phylogenies with metadata and live taxon placement will facilitate research into a diversity of pathogen species. This tool also emphasizes the sharing and standardization of data – including the phylogeny, multiple sequence alignments and sequence data, biological sample data, specimen vouchers and other associated metadata. The comprehensive nature of this tool is enabled by the wealth of *Phytophthora* research that has been collected and shared by our research group and others. We intend to contribute to and facilitate this trend by synthesizing these data streams. The efficacy of the live phylogeny presented here depends on open data sharing between researchers in the *Phytophthora* community.

## Conclusion

In conclusion, this study provides a curated, community-updated phylogeny for the genus *Phytophthora* incorporating both sequence data and biological metadata. This phylogeny will serve as a resource to the community as research on *Phytophthora* continues. Newly described species have been added to the phylogeny as a proof-of-concept, placing congruously with their species descriptions (Table 1). A microsatellite-based classifier for *P. infestans* genotypes was also developed to identify genotypes of *P. infestans* more rapidly. As the number of genotypes continues to grow and change, this tool too can be updated to keep pace with discovery by the research community. This live phylogeny format can also be extended to a diversity of pathogens to facilitate disease research.

## Acknowledgements

The authors would like to thank Montana Knight for gathering some of the sequence data used to develop the RAxML phylogeny and Kristen Pierce and Helen Nocito for their assistance with laboratory techniques and data collection. We thank Inga Meadows for help testing the tool. We thank David Cooke for sharing SSR profiles of European *P. infestans* lineages with us. We thank James White for implementing new custom color editing features in T-BAS v2.3. The graduate assistantship of A. Coomber was supported by the AgBioFEWS National Science Foundation National Research Training Grant 1828820, the USDA APHIS Plant Protection Act 7721 grant numbers APP12904 and APP16653 and the Functional Genomics program in the College of Biological Sciences at NC State.

