## SupplementalFigsTables for "An open-access T-BAS phylogeny for Emerging *Phytophthora* species"

Supplemental Figure 1. Histogram of Bruvo's genetic distance of all pairwise comparisons of isolates of *Phytophthora infestans* included in the SSR classifier reference dataset. The bimodal distribution is typical of species with a mixed reproduction systems like *Phytophthora infestans*. The first peak represents pairs which are closely related and derived from clonal reproduction. The second peak consists of pairs which are related via sexual recombination.

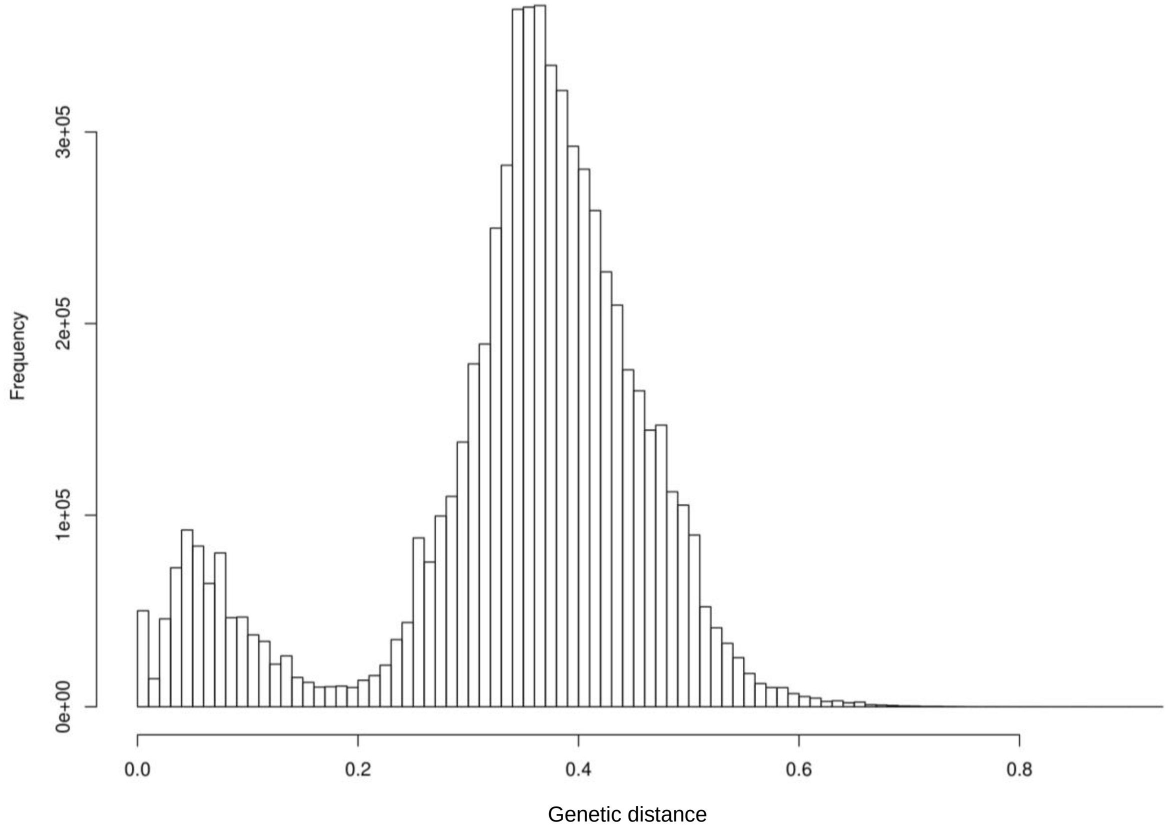

Supplemental Figure 2. Histogram showing the number of species in each clade in major *Phytophthora* phylogenies published since 2000. Subclade distinctions have been removed for simplification.

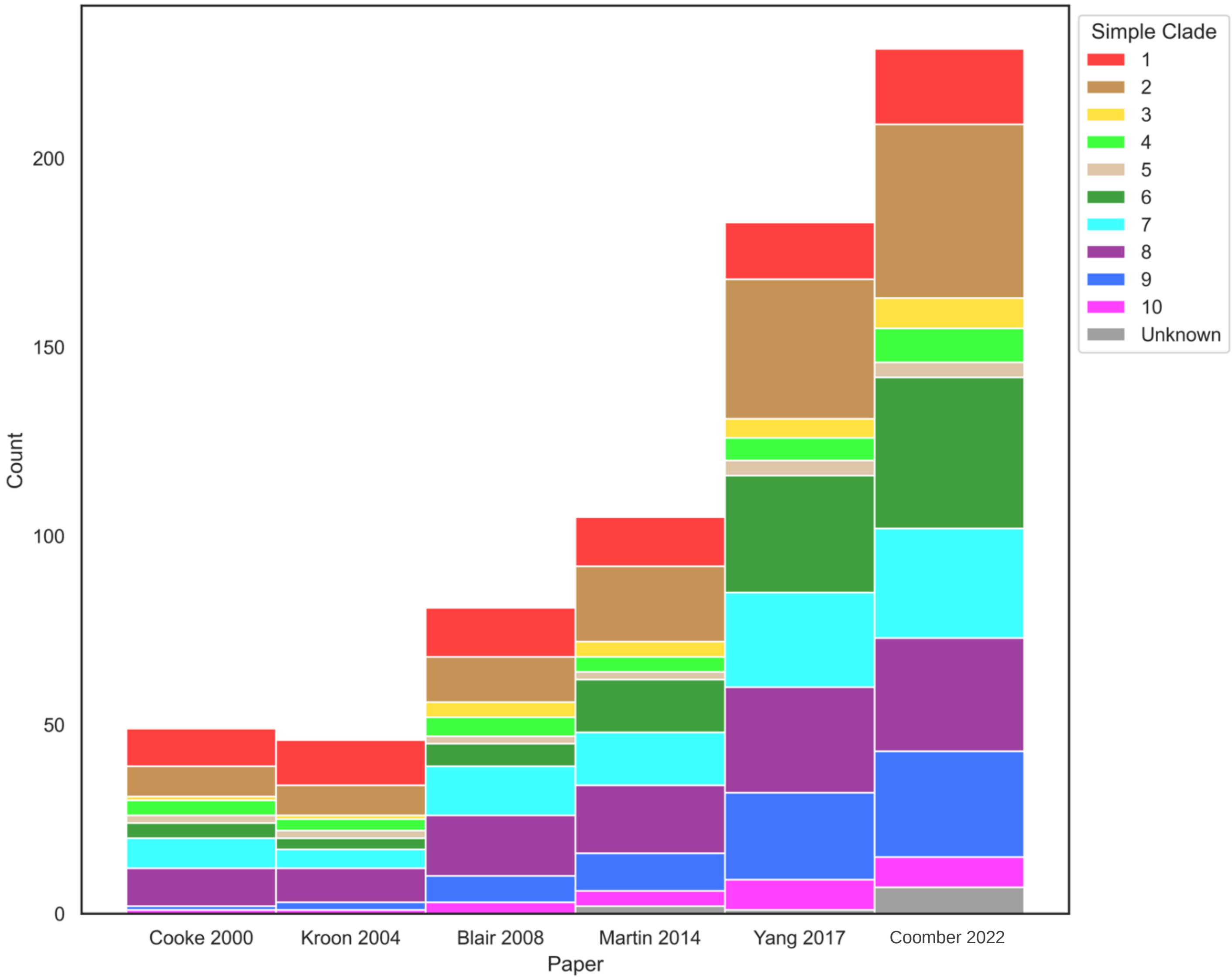

Supplemental Figure 3. Phylogeny of the genus *Phytophthora* inferred using maximum likelihood and 1,000 bootstrap replicates for the mitochondrial *CoxI* locus. Clade and subclade values are shown as variations in color hue. Bootstrap values (in percent) are shown for each branch. Branch lengths are drawn proportional to number of substitutions.

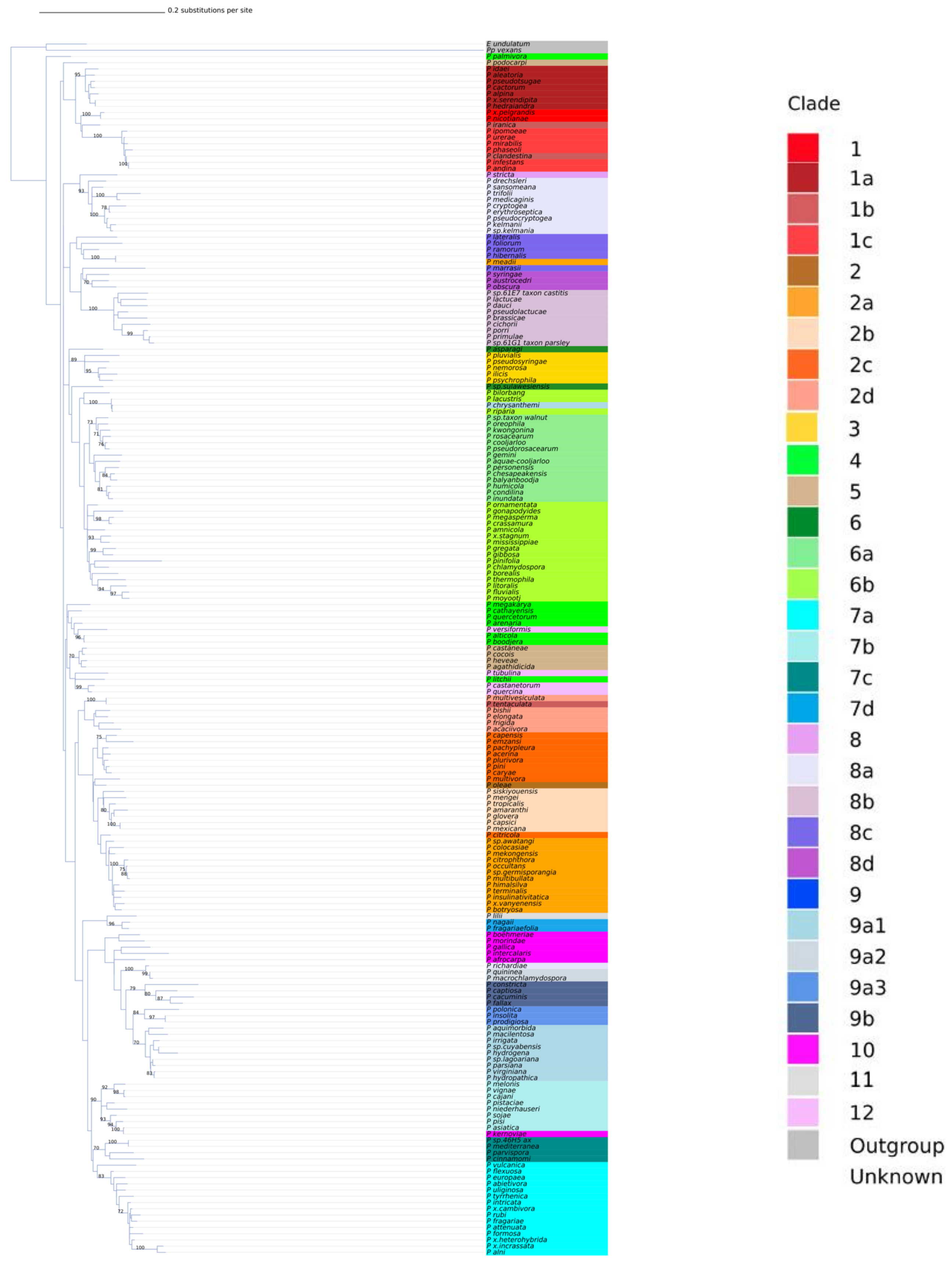

Supplemental Figure 4. Comparison of nine single-locus maximum likelihood phylogenetic trees for the *Phytophthora* genus inferred using Mesquite Hypha. A phylogenetic tree inferred using all eight nuclear loci (i.e., total evidence tree) serves as the backbone tree for comparison. Concordance or discordance between the trees for each locus is shown in grids at each node. Position within the grid corresponds to locus, and box color corresponds to the level of conflict or agreement. Bootstrap values for the total evidence tree are shown for each branch. Branch lengths are drawn proportional to number of substitutions.

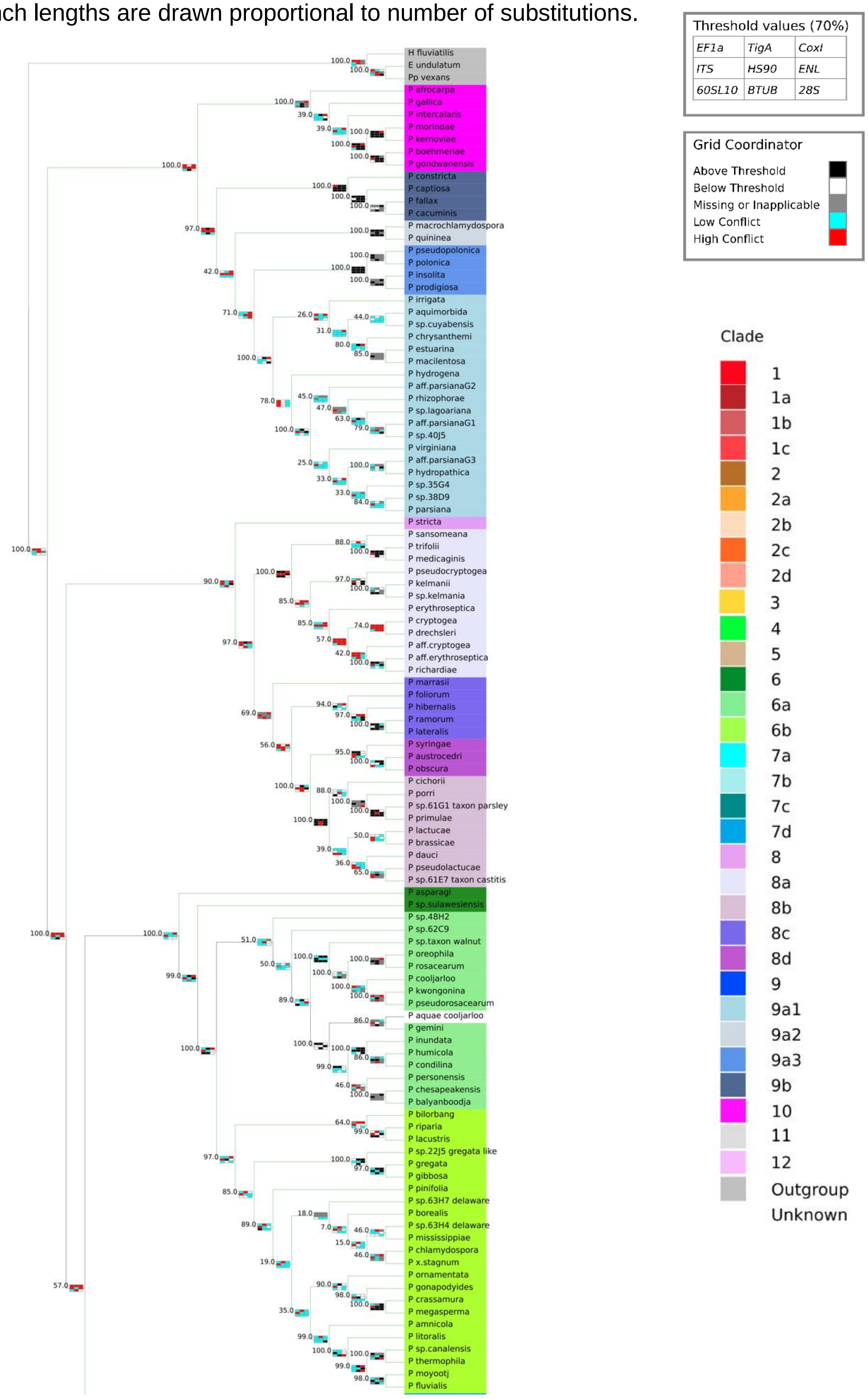

*Supplemental Figure 4 Continued*

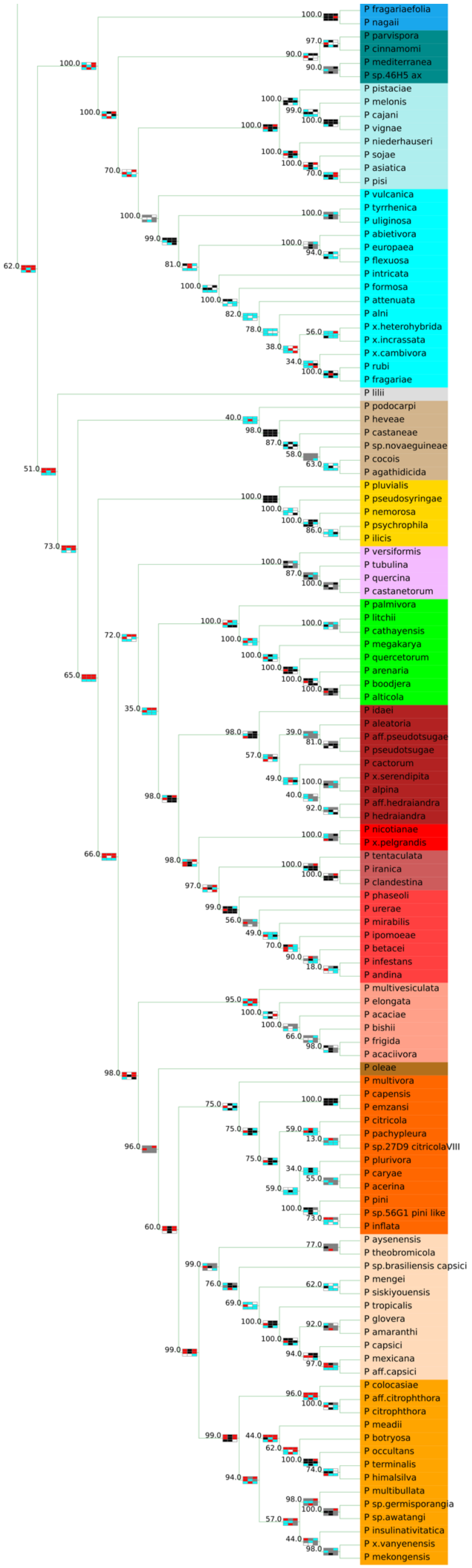

Supplemental Figure 5. Collapsed phylogeny of the genus *Phytophthora* showing clade 2 (orange) in detail. Subclade values are shown as variations in color hue. Bootstrap values are shown for each branch. Branch lengths are drawn proportional to number of substitutions. Phylogeny is inferred using maximum likelihood for 8 concatenated nuclear loci.

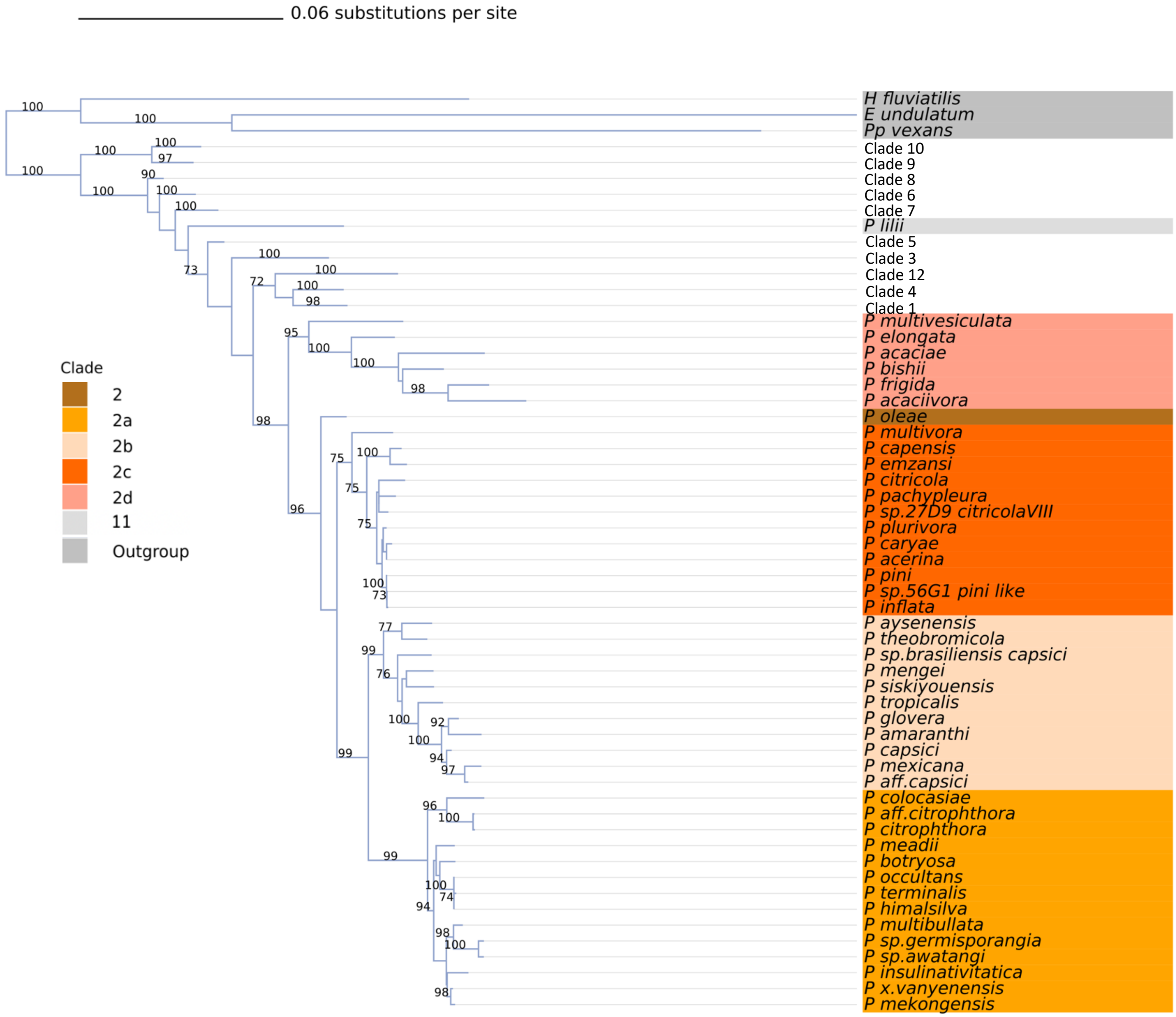

Supplemental Figure 6. Radial phylogeny of the genus *Phytophthora* inferred with maximum likelihood for an alignment of 8 concatenated nuclear genes. Coloring on the inner ring indicates clade. Coloring of the branches indicates sexual characteristic (homothallic, heterothallic, or unknown). Branch lengths are not to scale.

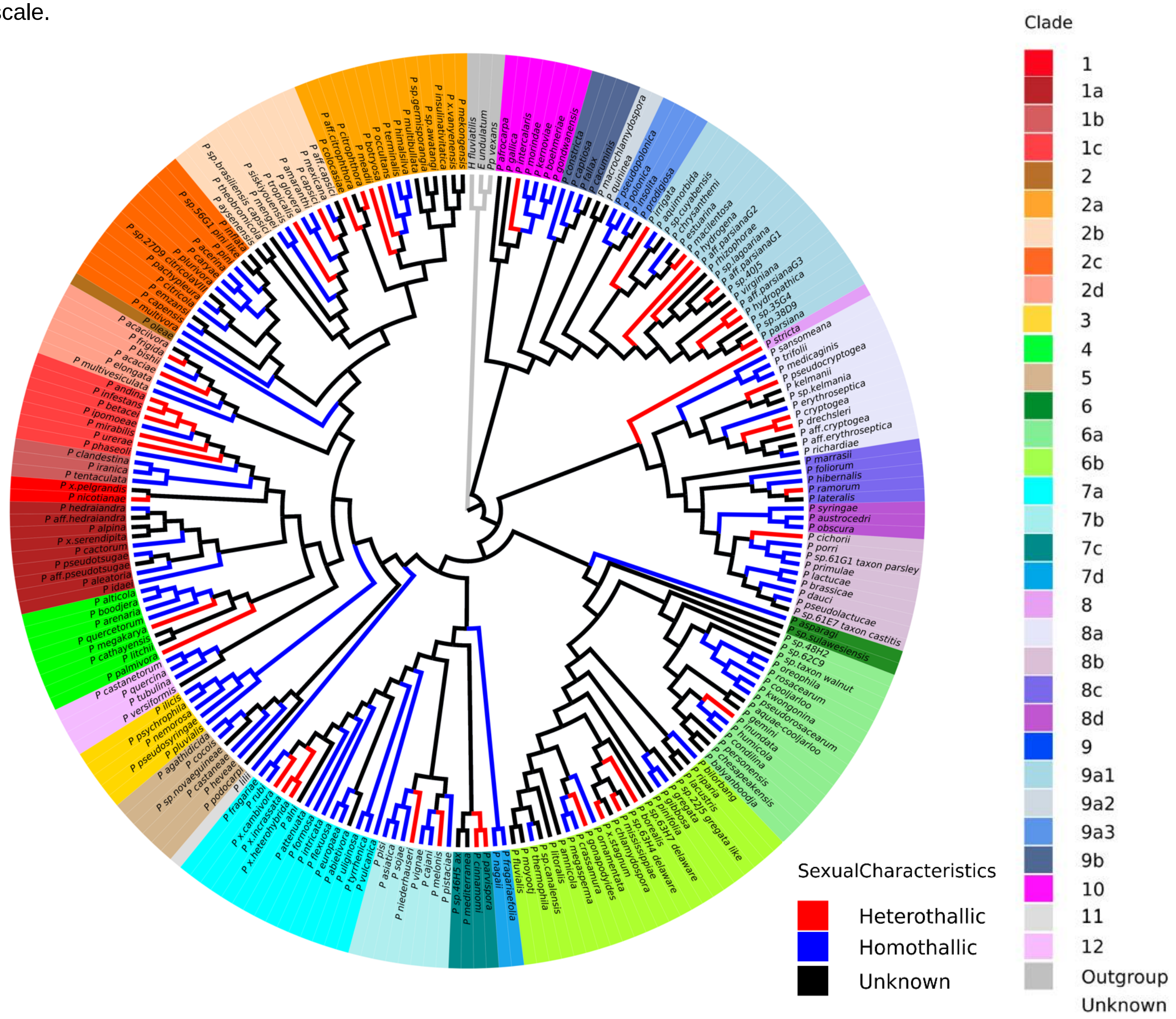

Supplemental Figure 7. Histogram showing the number of *Phytophthora* species in each sexual characteristic category in all major phylogenies published since 2000.

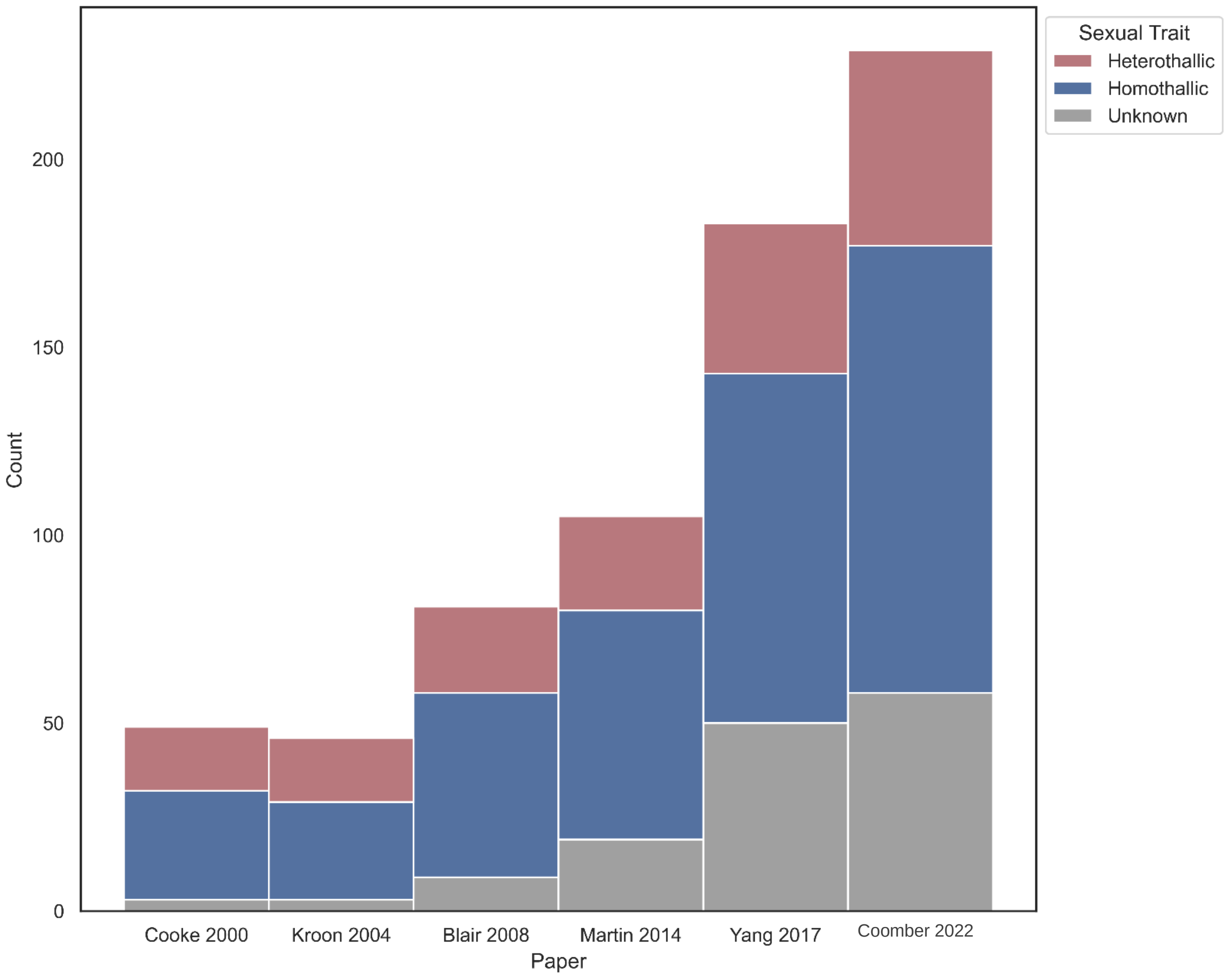

Supplemental Table 1. *Phytophthora* species names, synonyms, description paper, anc

| Species | Synonyms | Source | ITS | CoxI | HS90 | TigA | EF1a | 28S | Enol | 60SL10 | Btub |
|---|---|---|---|---|---|---|---|---|---|---|---|
| E. undulatum | [Outgroup] | Yang, X., Tyler, B. M., & Hong, C. (2017). An expanded phylogeny for the genus Phytophthora. IMA fungus, 8(2), 355-384. | KM061707.1 | HQ708990.1 | EU080443 | EU080445 | EU080442 | EU080444 | | EU080440 | EU080441 |
| H. fluviatilis | [Outgroup] | Yang, X., Tyler, B. M., & Hong, C. (2017). An expanded phylogeny for the genus Phytophthora. IMA fungus, 8(2), 355-384. | | | KX252672 | KX252674 | KX252670 | KX252673 | KX252671 | KX252668 | KX252669 |
| P. abietivora | None | Li, DW; Schultes, NP; LaMondia, JA; Cowles, RS. 2019. Phytophthora abietivora, A New Species Isolated from Diseased Christmas Trees in Connecticut, U.S.A. Plant Disease. 103(12):3057-3064 | MK163944.1 | MK164270.1 | MK164275.1 | | MK507885.1 | | | | MK164274.1 |
| P. acaciae | None | Tatiane C. Albuquerque Alves Dauri J. Tessmann Kelly L. Ivors Jean B. Ristaino & Ivaro F. dos Santos (2019) Phytophthora acaciae sp. nov. a new species causing gummosis of black wattle in Brazil Mycologia 111:3 445-455 | | KX396267 | ON063505, ON063506, ON063507 | | KX396326 | ON063499, ON063500, ON063501 | | ON063502, ON063503, ON063504 | KX396338 |
| P. acaciivora | None | Burgess, TI; Dang, QN; Le, BV; Pham, NQ; White, D; Pham, TQ. 2020. Phytophthora acaciivora sp. nov. associated with dying Acacia mangium in Vietnam. Fungal Systematics and Evolution. 6:243-252 | KX011263.1 | MN991990.1 | KX011239.1 | | OK267376.1 | | | OK533439.1 | MN991983.1 |
| P. acerina | None | Ginetti B; Moricca S; Squires J; Cooke D; Ragazzi A; Jung T. 2014. Phytophthora acerina sp. nov.; a new species causing bleeding cankers and dieback of Acer pseudoplatanus trees in planted forests in Northern Italy. Plant Pathology 63: 858-876. | MG518642.1 | KC156141.2 | KX250716, KX250723 | KX250718, KX250725 | KX250714, KX250721 | KX250717, KX250724 | KX250715, KX250722 | KX250712, KX250719 | KX250713, KX250720 |
| P. aff.capsici[1] | Classified as P. capsici by Mannon Gallegly. | Yang, X., Tyler, B. M., & Hong, C. (2017). An expanded phylogeny for the genus Phytophthora. IMA fungus, 8(2), 355-384. | | | KX250709 | KX250711 | KX250707 | KX250710 | KX250708 | KX250705 | KX250706 |
| P. aff.citrophthora[1] | Classified as P. citrophthora by Mannon Gallegly. | Yang, X., Tyler, B. M., & Hong, C. (2017). An expanded phylogeny for the genus Phytophthora. IMA fungus, 8(2), 355-384. | | | KX250618, EU080388 | KX250620, EU080390 | KX250616, EU080386 | KX250619, EU080389 | KX250617, EU080387 | KX250614, EU080384 | KX250615, EU080385 |
| P. aff.cryptogea[1] | Classified as P. cryptogea by Mannon Gallegly. | Yang, X., Tyler, B. M., & Hong, C. (2017). An expanded phylogeny for the genus Phytophthora. IMA fungus, 8(2), 355-384. | | | KX251969 | KX251971 | KX251967 | KX251970 | KX251968 | KX251965 | KX251966 |
| P. aff.erythroseptica[1] | Classified as P. erythroseptica by Mannon Gallegly. | Yang, X., Tyler, B. M., & Hong, C. (2017). An expanded phylogeny for the genus Phytophthora. IMA fungus, 8(2), 355-384. | | | KX251983, KX251976 | KX251971, KX251978 | KX251974, KX251981 | KX251977, | | | |
| P. aff.hedraiandra[1] | Classified as P. hedraiandra by Mannon Gallegly. | Yang, X., Tyler, B. M., & Hong, C. (2017). An expanded phylogeny for the genus Phytophthora. IMA fungus, 8(2), 355-384. | | | KX250415 | KX250417 | KX250413 | KX250416 | KX250414 | KX250411 | KX250412 |
| P. aff.parsianaG1[1,2] | In World Phytophthora Collection as P. zentmyerii. | Mostowfizadeh-Ghalamfarsa R Cooke DEL Banihashemi Z (2008) Phytophthora parsiana sp. nov. a new high-temperature tolerant species. Mycological Research 112: 783794. | | | KX252395, KX252401, EU080199 | EU080201 | KX252393, KX252399, EU080197 | KX252396, KX252402, EU080200 | KX252394, KX252400, EU080198 | KX252391, KX252397, EU080195 | KX252392, KX252398, EU080196 |
| P. aff.parsianaG2[1,2] | None | Mostowfizadeh-Ghalamfarsa R Cooke DEL Banihashemi Z (2008) Phytophthora parsiana sp. nov. a new high-temperature tolerant species. Mycological Research 112: 783794. | | | KX252437, KX252443 | | KX252435, KX252441 | KX252438, KX252444 | KX252436, KX252442 | KX252433, KX252439 | KX252434, KX252440 |
| P. aff.parsianaG3[1,2] | None | Mostowfizadeh-Ghalamfarsa R Cooke DEL Banihashemi Z (2008) Phytophthora parsiana sp. nov. a new high-temperature tolerant species. Mycological Research 112: 783794. | | | KX252449, KX252467, KX252479 | | KX252447, KX252465, KX252477 | KX252450, KX252468, KX252480 | KX252448, KX252466, KX252478 | KX252445, KX252463, KX252475 | KX252446, KX252464, KX252476 |
| P. aff.pseudotsugae[1] | Classified as P. pseudotsugae by Mannon Gallegly | Yang, X., Tyler, B. M., & Hong, C. (2017). An expanded phylogeny for the genus Phytophthora. IMA fungus, 8(2), 355-384. | | | KX250422 | KX250424 | KX250420 | KX250423 | KX250421 | KX250418 | KX250419 |
| P. afrocarpa | None | Bose, T; Hulbert, JM; Burgess, TI; Paap, T; Roets, F; Wingfield, MJ. 2021. Two novel Phytophthora species from the southern tip of Africa. Mycological Progress. 20(6):755-767 | MT762306.1 | MT762315.1 | MT762333.1 | | | | | | MT762324.1 |
| P. agathidicida | None | WeirBS; Paredes EP; Anand N; Uchida JY; Shaun R; Pennycook SEB; and Beever RE. 2015. A taxonomic revision of Phytophthora Clade 5 including two new species; Phytophthora agathidicida and P. cocois. Phytotaxa 205: 21-38(pg. 29). | MG602692.1 | MH620036.1 | KX251080 | KX251082 | KX251078 | KX251081 | KX251079 | KX251076 | KX251077 |
| P. aleatoria | None | Scott P. Taylor P. Gardner J. Purtolas A. Panda P. Addison S. . . . McDougal R. (2019 January 08). Phytophthora aleatoria sp. nov. associated with root and collar damage on Pinus radiata from nurseries and plantations. Retrieved September 08 2020 fromhttps://link.springer.com/article/10.1007/s13313-019-00631-5 | MK282209.1 | MK294173.1 | | | | | | | |
| P. alni | Classified as P. alni subsp. alni by Mannon Gallegly. | Unknown | MF356297.1 | KU681017.1 | KX251578, KX251585, KX251592, KX251599 | KX251580, KX251587, KX251594, KX251601 | KX251576, KX251583, KX251590, KX251597 | KX251579, KX251586, KX251593, KX251600 | KX251577, KX251584, KX251591, KX251598 | KX251574, KX251581, KX251588, KX251595 | KX251575, KX251582, KX251589, KX251596 |

| | | | | | | | | | | | |
|---|---|---|---|---|---|---|---|---|---|---|---|
| P. alpina | None | Bregant, C; Sanna, GP; Bottos, A; Maddau, L; Montecchio, L; Linaldeddu, BT. 2020. Diversity and Pathogenicity of Phytophthora Species Associated with Declining Alder Trees in Italy and Description of Phytophthora alpina sp. nov. Forests. 11(8, no. 848):1-16 | MT707331.1 | MT729667.1 | | | | | | | MT729672.1 |
| P. alticola | None | MasekoB; BurgessTI; CoutinhoTA; WingfieldMJ. 2007. Two new Phytophthora species from South Africa. Mycol. Res. 111: 1321-1338. | KX247605.1 | KX247591.1 | KX251010 | KX251012 | KX251008 | KX251011 | KX251009 | KX251006 | KX251007 |
| P. amaranthi | None | Ann P Huang J Tsai J Ko W. 2016. Morphological molecular and pathological characterization of Phytophthora amaranthi sp. nov. from amaranth in Taiwan. J. Phytopathol. 164: 94101. | MG783374.1 | MH477739.1 | | | MK864032.1 | | | | KJ179949.1, KJ179948.1 |
| P. amnicola | None | BurgessTI; HberliD; HardyGE StJ; StukelyMJC;and Jung T. 2012. Phytophthora amnicola T. I. Burgess & T. Jung; sp. nov. Persoonia 28: 140-141. | MN545896.1 | JQ029950.1 | KX251171, KX251178 | KX251173, KX251180 | KX251169, KX251176 | KX251172, KX251179 | KX251170, KX251177 | KX251167, KX251174 | KX251168, KX251175 |
| P. andina | P. infestans sensu lato | OlivaRF; KroonLPMN; ChacnG; FlierWG; RistainoJB; andForbesGA. 2010. Phytophthora andina sp. nov.; a newly identified heterothallic pathogen of solanaceous hosts in the Andean highlands. Plant Pathol. 59: 613-625(pg. 622). | MK496515.1 | MH286887.1 | KX250464, KX250471, EU080186 | KX250466, KX250473, EU080188 | KX250462, KX250469, EU080184 | KX250465, KX250472, EU080187 | KX250463, KX250470, EU080185 | KX250460, KX250467, EU080182 | KX250461, KX250468, EU080183 |
| P. aquae-cooljarloo | None | Caboň, M., Adams, G. C., Fechner, N., García, D., Gené, J., Halling, R. E., ... & Dearnaley, J. D. W. (2020). Fungal Planet description sheets: 1112–1181. Persoonia-Molecular Phylogeny and Evolution of Fungi, 45, 251. | JN547639.1 | MT210466.1 | MT210480.1 | | | | | | MT210472.1 |
| P. aquimorbida | None | HongCX; RichardsonPA; HaoW; ShimireSR; KongP; MoormanGW; Lea-CoxJD; and RossDS. 2012. Phytophthora aquimorbida sp. nov. and Phytophthora taxon 'aquatilis' recovered from irrigation reservoirs and a stream in Virginia; USA. Mycologia 104: 1097-1108. | MG783376.1 | MH136847.1 | KX252242, KX252249, KX252252 | KX252244, KX252249, KX252258 | KX252240, KX252247, KX252254 | KX252243, KX252250, KX252257 | KX252241, KX252248, KX252255 | KX252238, KX252245, KX252252 | KX252239, KX252246, KX252253 |
| P. arenaria | None | ReaAJ; BurgessTI; HardyGE StJ; StukelyMJC; and JungT. 2011. Two novel and potentially endemic species of Phytophthora associated with episodic dieback of Kwongan vegetation in the south-west of Western Australia. Plant Pathol. 60: 1055-1068. | MG783377.1 | KJ396696.1 | KX251017, KX251024 | KX251019, KX251026 | KX251015, KX251022 | KX251018, KX251025 | KX251016, KX251023 | KX251013, KX251020 | KX251014, KX251021 |
| P. asiatica | P. cinnamomi var. robiniae | Rahman MZ; Mukobata H; Suga H;and Kageyama K. 2014. Phytophthora asiatica sp. nov.; a new species causing leaf and stem blight of kudzu in Japan. Mycol. Progress 13: 759-769. | MG783379.1 | MH620062.1 | KX251655, KX251662, KX251669 | KX251657, KX251664, KX251671 | KX251653, KX251660, KX251667 | KX251656, KX251663, KX251670 | KX251654, KX251661, KX251668 | KX251651, KX251658, KX251665 | KX251652, KX251659, KX251666 |
| P. asparagi | None | GrankeLL; SaudeC; WindstamST; WebsterBJ; and HausbeckMK. 2012. Phytophthora asparagi Saude & Hausbeck; sp. nov. Persoonia 28: 146-147. | MN545892.1 | HQ012844.1 | KX251470, KX251477 | KX251472, KX251479 | KX251468, KX251475 | KX251471, KX251478 | KX251469, KX251476 | KX251466, KX251473 | KX251467, KX251474 |
| P. attenuata | None | Jung T Jung MH Scanu B Seress D Kovcs GM Maia C Prez-Sierra A Chang T-T Chandelier A Heungens K van Poucke K Abad-Campos P Lon M Cacciola SO Bakonyi J.2017. Six new Phytophthora species from ITS Clade 7a including two sexually functional heterothallic hybrid species detected in natural ecosystems in Taiwan. Persoonia 38: 100135. | MN872723.1 | MN866073.1 | KX251613 | KX251615 | KX251611 | KX251614 | KX251612 | KX251609 | KX251610 |
| P. austrocedri | P. austrocedrae | GreslebinAG HansenEM and SuttonW. 2007. Phytophthora austrocedrae sp. nov. a new species associated with Austrocedrus chilensis mortality in Patagonia (Argentina). Mycol. Res. 111: 308316. | MG783380.1 | HQ917881.1 | KX252165, KX252172 | KX252167, KX252174 | KX252163, KX252170 | KX252166, KX252173 | KX252164, KX252171 | KX252161, KX252168 | KX252162, KX252169 |
| P. aysenensis | None | Crous PW, Wingfield MJ, Chooi YH, et al. Fungal Planet description sheets: 1042-1111. Persoonia. 2020;44:301-459. doi:10.3767/persoonia.2020.44.11 | MN557838.1 | | | | | MN557839.1 | | | MN557840.1 |
| P. balyanboodja | None | Burgess TI Simamora AV White D Wiliams B Schwager M Stukely MJC Hardy GE StJ.2018. New species from Phytophthora Clade 6a: evidence for recent radiation. Persoonia 41: 117. | KJ372259.1 | MF326862.1 | MF326893.1, MF326892.1 | | MK864034.1 | | | | MN207267.1, MF326807.1, MF326806.1 |
| P. betacei | None | M.F. Mideros et al.: Phytophthora betacei a new species within clade 1c | | | ON063525, ON063526, ON063527 | ON063523, ON063524 | ON063511, ON063512, ON063513 | ON063508, ON0635089, ON063510 | ON063514, ON063515, ON063522 | ON063516, ON063518, ON063519 | ON063517, ON063520, ON063521 |
| P. bilorbang | None | AghighiS; Hardy GE StJ; Scott JK; and Burgess TI. 2012. Phytophthora bilorbang sp. nov.; a new species associated with the decline of Rubus anglocandicans (European blackberry) in Western Australia. Eur. J. Plant Pathol. 133: 841-855. | MG696534.1 | JN547644.1 | KX251184 | KX251186 | | KX251185 | KX251183 | KX251181 | KX251182 |
| P. bishii | P. bisheria | Variant spelling Phytophthora bisheria Z.G. Abad J.A. Abad & F.J. Louws 2008 | MN549026.1 | | KX011243.1 | | | | | | MN991985.1 |
| P. boehmeriae | None | SawadaK. 1927. Descriptive catalogue of the Formosan fungi III. Report of the Department of Agriculture; Government Research Institute of Formosa 27: 1-62. | KX396302.1 | KF317111.1 | EU080165, KX252644 | EU080167, KX252646 | EU080163, KX252642 | EU080166, KX252645 | EU080164, KX252643 | EU080161, KX252640 | EU080162, KX252641 |

| | | | | | | | | | | | |
|---|---|---|---|---|---|---|---|---|---|---|---|
| P. boodjera | None | Simamora AV Stukely MJC Hardy GE StJ Burgess TI. 2015. Phytophthora boodjera sp. nov. a damping-off pathogen in production nurseries and from urban and natural landscapes with an update on the status of P. alticola. IMA Fungus 6: 319335 (pg. 326). | KJ372242.1 | KJ396685.1 | KJ396715.1,<br>KJ396714.1,<br>KJ396713.1,<br>KJ396712.1,<br>KJ396711.1,<br>KJ396710.1,<br>KJ396709.1,<br>KJ396708.1,<br>KJ396707.1,<br>KJ396706.1,<br>KJ396705.1,<br>KJ396704.1,<br>MK020277.1 | | MK864036.1 | | KJ396743.1,<br>KJ396742.1,<br>KJ396741.1,<br>KJ396740.1,<br>KJ396739.1,<br>KJ396738.1,<br>KJ396737.1,<br>KJ396736.1,<br>KJ396735.1,<br>KJ396734.1,<br>KJ396733.1,<br>KJ396732.1 | | KJ372288.1,<br>KJ372287.1,<br>KJ372286.1,<br>KJ372285.1,<br>KJ372284.1,<br>KJ372283.1,<br>KJ372281.1,<br>KJ372280.1,<br>KJ372279.1,<br>KJ372278.1,<br>KJ372277.1,<br>KJ372276.1 |
| P. borealis | None | HansenEM; ReeserPW; and SuttonW. 2012. Phytophthora borealis and Phytophthora riparia; new species in Phytophthora ITS Clade 6. Mycologia 104: 1133-1142. | MF034096.1 | JQ626623.1 | KX251191 | KX251193 | KX251189 | KX251192 | KX251190 | KX251187 | KX251188 |
| P. botryosa | None | CheeKH. 1969. Variability of Phytophthora species from Hevea brasiliensis. T. Brit. Mycol. Soc. 52: 425-436. | KC247900.1 | KF317093.1 | KX250527,<br>KX250534,<br>KX250541,<br>EU079938 | KX250529,<br>KX250536,<br>KX250543,<br>EU079940 | KX250525,<br>KX250532,<br>KX250539,<br>EU079936 | KX250528,<br>KX250535,<br>KX250542,<br>EU079939 | KX250526,<br>KX250533,<br>KX250540,<br>EU079937 | KX250523,<br>KX250530,<br>KX250537,<br>EU079934 | KX250524,<br>KX250531,<br>KX250538,<br>EU079935 |
| P. brassicae | None | Man in 't VeldWA; de CockAWAM; IlievaE; and LvesqueCA. 2002. Gene flow analysis of Phytophthora porri reveals a new species: Phytophthora brassicae sp. nov. Eur. J. Plant Pathol. 108: 51-62. | FJ801871.1 | AY564198.1 | KX251997,<br>KX252004 | KX251999,<br>KX252006 | KX251995,<br>KX252002 | KX251998,<br>KX252005 | KX251996,<br>KX252003 | KX251993,<br>KX252000 | KX251994,<br>KX252001 |
| P. cactorum | Phytophthora paeoniae,<br>Pernospora fagi | SchrterJ. 1886. pp. 129-256 (236). In: F. Cohn; Kryptogamen-Flora von Schlesien. Band 3; Heft 3; Pilze J.U. Kern's Verlag; Breslau; 1889; pp. 1-814. | MG696466.1 | AY129174.1 | KX250373,<br>EU080288,<br>KX250380, | KX250375,<br>EU080290,<br>KX250382, | KX250371,<br>EU080286,<br>KX250378 | KX250374,<br>EU080289,<br>KX250381 | KX250372,<br>EU080287,<br>KX250379 | KX250369,<br>EU080284,<br>KX250376 | KX250370,<br>EU080285,<br>KX250377 |
| P. cacuminis | None | Khaliq I. St J Hardy G. E. McDougall K. L. & Burgess T. I. (2019 January). Phytophthora species isolated from alpine and sub-alpine regions of Australia including the description of two new species; Phytophthora cacuminis sp. nov and Phytophthora oreophila sp. nov. Retrieved September 15 2020 from https://www.sciencedirect.com/science/article/pii/S1878614618302332?via%3Dihub | MG542997.1 | MG543011.1 | MG543033.1,<br>MG543032.1 | | | | | | MG543046.1,<br>MG543045.1 |
| P. cajani | P. drechsleri var cajani | AminKS; BaldevB; and WilliamsFJ. 1978. Phytophthora cajani; a new species causing stem blight on Cajanus cajan. Mycologia 70: 171-176. | MT940649.1 | KF358234.1 | KX251676,<br>KX251683,<br>KX251690 | KX251678,<br>KX251685,<br>KX251692, | KX251674,<br>KX251681,<br>KX251688, | KX251677,<br>KX251684,<br>KX251691, | KX251675,<br>KX251682,<br>KX251689 | KX251672,<br>KX251679,<br>KX251686 | KX251673,<br>KX251680,<br>KX251687 |
| P. capensis | None | BezuidenhoutCM; DenmanS; KirkSA; BothaWJ; MostertL; and McLeodA. 2010. Phytophthora taxa associated with cultivated Agathosma; with emphasis on the P. citricola complex and P. capensis sp. nov. Persoonia 25: 32-49(pg. 45). | NR. 147872.1 | MN866070.1 | KX250730,<br>KX250737,<br>KX250744 | KX250732,<br>KX250739,<br>KX250746 | KX250728,<br>KX250735,<br>KX250742 | KX250731,<br>KX250738,<br>KX250745 | KX250729,<br>KX250736,<br>KX250743 | KX250726,<br>KX250733,<br>KX250740 | KX250727,<br>KX250734,<br>KX250741 |
| P. capsici | P. parasitica var capsici, P. hydrophila, P. mexicana | LeonianLH. 1922. Stem and fruit blight of pepper caused by Phytophthora capsici species nov. Phytopathology 12: 401-408. | LT707526.1 | KJ631596.2 | KX250639,<br>EU080855,<br>EU079547 | KX250641,<br>EU080857,<br>EU079549 | KX250637,<br>EU080853,<br>EU079545 | KX250640,<br>EU080856,<br>EU079548 | KX250638,<br>EU080854,<br>EU079546, | KX250635,<br>EU080851,<br>EU079543 | KX250636,<br>EU080852,<br>EU079544 |
| P. captiosa | None | DickMA; DobbieK; CookeDEL; and BrasierCM. 2006. Phytophthora captiosa sp. nov. and P. fallax sp. nov. causing crown dieback of Eucalyptus in New Zealand. Mycol. Res. 110: 393-404. | MG543000.1 | MG543005.1 | KX252551,<br>EU079662,<br>KX252558,<br>EU079669 | KX252553,<br>EU079664,<br>KX252560,<br>EU079671 | KX252549,<br>EU079660,<br>KX252556,<br>EU079667 | KX252552,<br>EU079663,<br>KX252559,<br>EU079670 | KX252550,<br>EU079661,<br>KX252557,<br>EU079668 | KX252547,<br>EU079658,<br>KX252554,<br>EU079665 | KX252548,<br>EU079659,<br>KX252555,<br>EU079666 |
| P. caryae | None | Brazee NJ Yang X and Hong CX. 2016. Phytophthora caryae sp. nov. a new species recovered from streams and rivers in the eastern United States. Plant Pathology 66: 805817. | MT647267.1 | KJ631588.2 | | | | | | | KU695469.1,<br>KP749393.1,<br>KJ631574.1,<br>KJ631573.1,<br>KJ631572.1 |
| P. castaneae | P. katsurae | KatsuraK. 1976. Two new species of Phytophthora causing damping-off of cucumber and trunk rot of chestnut. Trans. Mycol. Soc. Japan 17: 238-242. | KU682560.1 | MN866065.1 | KX251087,<br>KX251094,<br>KX251101 | KX251089,<br>KX251096,<br>KX251103 | KX251085,<br>KX251092,<br>KX251099 | KX251088,<br>KX251095,<br>KX251102 | KX251086,<br>KX251093,<br>KX251100 | KX251083,<br>KX251090,<br>KX251097 | KX251084,<br>KX251091,<br>KX251098 |
| P. castanetorum | None | Jung T Horta Jung M Cacciola SO Cech T Bakonyi J Seress D Mosca S Schena L Seddaiu S Pane A Magnano di San Lio G Maia C Cravador A Franceschini A and Scanu B. 2017. Multiple new cryptic pathogenic Phytophthora species from Fagaceae forests in Austria Italy and Portugal. IMA Fungus 8 (2): 219244. | MF036185.1 | MF036267.1 | MF036246.1,<br>MF036245.1,<br>MF036244.1,<br>MF036243.1,<br>MF036242.1,<br>MF036241.1,<br>MF036240.1 | | | | | | MF036220.1,<br>MF036219.1,<br>MF036218.1,<br>MF036217.1,<br>MF036216.1,<br>MF036215.1,<br>MF036214.1 |

| | | | | | | | | | | | |
|---|---|---|---|---|---|---|---|---|---|---|---|
| P. cathayensis | None | Morales-Rodríguez, C; Wang, Y; Martignoni, D; Vannini, A. 2021. Phytophthora cathayensis sp. nov., a new species pathogenic to Chinese Hickory (Carya cathayensis) in southeast China. Fungal Systematics and Evolution. 7:99-111 | MN385740.1 | MN692210.1 | MT063109.1 | MT063112.1 | MN721978.1 | MN721971.1 | MT063105.1 | MN721974.1 | MT063102.1 |
| P. chesapeakensis | None | Man in t Veld W. A. Rosendahl K. C. Van Rijswick P. C. Meffert J. P. Boer E. Westenberg M. . . . Govers L. L. (2018 August 28). Multiple Halophytophthora spp. and Phytophthora spp. including P. gemini P. inundata and P. chesapeakensis sp. nov. isolated from the seagrass Zostera marina in the Northern hemisphere. Retrieved September 10 2020 from https://link.springer.com/article/10.1007/s10658-018-1561-1/tables/1 | KX172093.1 | KX172095.1 | | | | | | | |
| P. chlamydospora | None | Hansen EM Reeser P Sutton W and Brasier CM. 2015. Redesignation of Phytophthora taxon Pgchlamydo as Phytophthora chlamydospora sp. nov. North American Fungi 10 (2): 114. | MG696468.1 | KU682588.1 | MK020285.1 | | MH358970.1 | | | | MH493919.1 |
| P. chrysanthemi | None | NaherM; MotohashK; WatanabeH; ChikuoY; SendaM; SugaH; BrasierC; and KageyamaK. 2011. Phytophthora chrysanthemi sp. nov.; a new species causing root rot of chrysanthemum in Japan. Mycol. Prog. 10: 21-31. | MG865472.1 | MH620093.1 | KX252263, KX252270 | KX252265, KX252272 | KX252261, KX252268 | KX252264, KX252271 | KX252262, KX252269 | KX252259, KX252266 | KX252260, KX252267 |
| P. cichorii | P. aff. cichorii[1] | BertierL; BrouwerH; de CockAWAM; CookeDEL;OlssonCHB; and HfteM. 2013. The expansion of Phytophthora clade 8b: three new species associated with winter grown vegetable crops. Persoonia 31: 63-76. | KC478774.1 | MH620083.1 | KX252011 | KX252013 | KX252009 | KX252012 | KX252010 | KX252007 | KX252008 |
| P. cinnamomi | None | RandsRD. 1922. Streepkanker van Kaneel; veroorzaakt door Phytophthora cinnamomi n. sp. Meded. Inst. voor Plantenziekt 54: 1-54. | MF541547.1 | MN866069.1 | KX251801, KX251808, KX251815 | KX251803, KX251810, KX251817 | KX251799, KX251806, KX251813 | KX251802, KX251809, KX251816 | KX251800, KX251807, KX251814 | KX251797, KX251804, KX251811 | KX251798, KX251805, KX251812 |
| P. citricola | Phytophthora cactorum var. applanata, P. pini (P. citricola group I was reclassified as P. pini) | SawadaK. 1927. Descriptive catalogue of the Formosan fungi III. Report of the Department of Agriculture; Government Research Institute of Formosa 27: 1-62. | MF115422.1 | MN866097.1 | KX250751, KX250758, KX250842 | KX250753, KX250760, KX250844 | KX250749, KX250756, KX250840 | KX250752, KX250759, KX250843 | KX250750, KX250757, KX250841 | KX250747, KX250754, KX250838 | KX250748, KX250755, KX250839 |
| P. citrophthora | Phytophthora imperfecta var. citrophthora | LeonianLH. 1925. Physiological studies on the genus Phytophthora. American Journal of Botany 12: 444-498. (Phytophthora citrophthora) | JN618718.1 | KJ631594.2 | KX250548, KX250555 | KX250550, KX250557 | KX250546, KX250553 | KX250549, KX250556 | KX250547, KX250554 | KX250544, KX250551 | KX250545, KX250552 |
| P. clandestina | None | TaylorPA; PascoeIE; and GreenhalghFC. 1985. Phytophthora clandestina sp. nov. in roots of subterranean clover. Mycotaxon 22: 77-85. | EU257132.1 | AY564172.1 | EU079870, KX250429, KX250436 | EU079872, KX250431, KX250438 | EU079868, KX250427, KX250434 | EU079871, KX250430, KX250437 | EU079869, KX250428, KX250435 | EU079866, KX250425, KX250432 | EU079867, KX250426, KX250433 |
| P. cocois | None | WeirBS; Paredes EP; Anand N; Uchida JY; Shaun R; Pennycook SEB; and Beever RE. 2015. A taxonomic revision of Phytophthora Clade 5 including two new species; Phytophthora agathidicida and P. cocois. Phytotaxa 205: 21-38(pg. 32). | MG865478.1 | MH136874.1 | KX251108 | KX251110 | KX251106 | KX251109 | KX251107 | KX251104 | KX251105 |
| P. colocasiae | Phytophthora parasitica var. colocasiae | Raciborski; M. 1900. Parasitic algae and fungi; Java. Batavia Bulletin of the New York State Museum 19: 189(pg. 9). | GQ148558.1 | AY129173.1 | KX250562, KX250569 | KX250564, KX250571 | KX250560, KX250567 | KX250563, KX250570 | KX250561, KX250568 | KX250558, KX250565 | KX250559, KX250566 |
| P. condilina | None | Burgess TI Simamora AV White D Wiliams B Schwager M Stukely MJC Hardy GE StJ. 2018. New species from Phytophthora Clade 6a: evidence for recent radiation. Persoonia 41: 117. | MG707826.1 | HQ012883.1 | MF326872.1, MF326871.1, MF326870.1, MF326869.1, MF326868.1, MF326867.1 | | MK864040.1 | | | | MN207271.1, MF326815.1, MF326814.1, MF326813.1, MF326812.1, MF326811.1, MF326810.1, MF326809.1, MF326808.1, HQ012928.1, HQ012927.1, MK020292.1 |
| P. constricta | None | ReaAJ; BurgessTI; HardyGEStJ; StukelyMJC; and JungT. 2011. Two novel and potentially endemic species of Phytophthora associated with episodic dieback of Kwongan vegetation in the south-west of Western Australia. Plant Pathol. 60: 1055-1068. | EU301148.2 | KC733450.1 | KX252565 | KX252567 | KX252563 | KX252566 | KX252564 | KX252561 | KX252562 |
| P. cooljarloo | None | Burgess TI Simamora AV White D Wiliams B Schwager M Stukely MJC Hardy GE StJ. 2018. New species from Phytophthora Clade 6a: evidence for recent radiation. Persoonia 41: 117. | HQ012961.1 | HQ012885.1 | HQ012929.1, HQ012925.1, MK020294.1 | | MK864041.1 | | | | MF326817.1, MF326816.1 |
| P. crassamura | None | Scanu B; Linaldeddu BT; Deidda A; and Jung T. 2015. Diversity of Phytophthoraspecies from declining Mediterranean maquis vegetation; including two new species; Phytophthora crassamura and Phytophthora ornamentata sp. nov. PLoS ONE 10 (12): 1-24(pg. 24). | MG554691.1 | KP863482.1 | KX251198, KX251205 | KX251200, KX251207 | KX251196, KX251203 | KX251199, KX251206 | KX251197, KX251204 | KX251194, KX251201 | KX251195, KX251202 |

| | | | | | | | | | | |
|---|---|---|---|---|---|---|---|---|---|---|
| P. cryptogea | Phytophthora oryzae | PethybridgeGHand LaffertyHA. 1919. A disease of tomato and other plants caused by a new species of Phytophthora. Scient. Proc. R. Dubl. Soc. N.S. 15: 487-503. | HM004230.1 | KJ469047.1 | KX251871 | KX251873 | KX251869 | KX251872 | KX251870 | KX251867 | KX251868 |
| P. dauci | None | BertierL; BrouwerH; de CockAWAM; CookeDEL; OlssonCHB; and HfteM. 2013. The expansion of Phytophthora Clade 8b: three new species associated with winter grown vegetable crops. Persoonia 31: 63-76. | GU258834.1 | MH620084.1 | KX252018, KX252025, KX252032, KX252039 | KX252020, KX252027, KX252034, KX252041 | KX252016, KX252023, KX252030, KX252037 | KX252019, KX252026, KX252033, KX252040 | KX252017, KX252024, KX252031, KX252038 | KX252014, KX252021, KX252028, KX252035 | KX252015, KX252022, KX252029, KX252036 |
| P. drechsleri | Phytophthora erythroseptica var. drechsleri | TuckerCM. 1931. Taxonomy of the genus Phytophthora de Bary. Research Bulletin of the Missouri Agricultural Experiment Station 153: 1-208(pg. 188). | L76549.1 | MN866087.1 | KX251878, KX251885, KX251892, EU079510 | KX251880, KX251887, KX251894, EU079512 | KX251876, KX251883, KX251890, EU079508 | KX251879, KX251886, KX251893, EU079511 | KX251877, KX251884, KX251891, EU079509 | KX251874, KX251881, KX251888, EU079506 | KX251875, KX251882, KX251889, EU079507 |
| P. elongata | None | ReaAJ; JungT; BurgessTI; StukelyMJC; GilesE; and Hardy GEStJ. 2010. Phytophthora elongata sp. nov.; a novel pathogen from the Eucalyptus marginata forest of Western Australia. Australas. Plant Path. 39: 477-491. | MG865485.1 | MH620031.1 | KX250884, KX250891, KX250898 | KX250886, KX250893, KX250900 | KX250882, KX250889, KX250896 | KX250885, KX250892, KX250899 | KX250883, KX250890, KX250897 | KX250880, KX250887, KX250894 | KX250881, KX250888, KX250895 |
| P. emzansi | Now P. emzansi, P. taxon emzansi | Bose, T., Hulbert, J. M., Burgess, T. I., Paap, T., Roets, F., & Wingfield, M. J. (2021). Two novel Phytophthora species from the southern tip of Africa. *Mycological Progress*, *20* (6), 755-767. | GU191228.1 | MH620029.1 | KX250863, KX250870 | KX250865, KX250872 | KX250861, KX250868 | KX250864, KX250871 | KX250862, KX250869 | KX250859, KX250866 | KX250860, KX250867 |
| P. erythroseptica | Phytophthora himalayensis | PethybridgeGH. 1913. On the rotting of potato tubers by a new species of Phytophthora having a method of sexual reproduction hitherto undescribed. Scientific Proceedings of the Royal Dublin Society 13: 529-565(pgs. 547-548). | KJ755119.1 | MN566606.1 | KX251899 | KX251901 | KX251897 | KX251900 | KX251898 | KX251895 | KX251896 |
| P. estuarina | None | MaranoAL Jesus & Pires-Zottar. 2016.Phytophthora estuarina. inFungal Diversity 78: 194.LiGJ HydeKD ZhaoRL et al. Fungal Diversity (2016) 78: 1. https://doi.org/10.1007/s13225-016-0366-9 | KT886034.1 | | | | | | | | |
| P. europaea | None | JungT; Hansen EM; WintonL; OsswaldW; and DelatourC. 2002. Three new species of Phytophthora from European oak forests. Mycol. Res. 106: 397-411. | KJ755092.1 | MH620055.1 | KX251512, KX251519, KX251526, KX251606 | KX251514, KX251521, KX251528, KX251608 | KX251510, KX251517, KX251524, KX251604 | KX251513, KX251520, KX251527, KX251607 | KX251511, KX251518, KX251525, KX251605 | KX251508, KX251515, KX251522, KX251602 | KX251509, KX251516, KX251523, KX251603 |
| P. fallax | None | DickMA; DobbieK; CookeDEL; and BrasierCM. 2006. Phytophthora captiosa sp. nov. and P. fallax sp. nov. causing crown dieback of Eucalyptus in New Zealand. Mycol. Res. 110: 393-404 (pg398). | MG542994.1 | KC733451.1 | KX252572, KX252579, KX252586, EU080038 | KX252574, KX252581, KX252588, EU080040 | KX252570, KX252577, KX252584, EU080036 | KX252573, KX252580, KX252587, EU080039 | KX252571, KX252578, KX252585, EU080037 | KX252568, KX252575, KX252582, EU080034 | KX252569, KX252576, KX252583, EU080035 |
| P. flexuosa | None | Jung T Jung MH Scanu B Seress D Kovcs GM Maia C Prez-Sierra A Chang T-T Chandelier A Heungens K van Poucke K Abad-Campos P Lon M Cacciola SO and Bakonyi J.2017 Six new Phytophthora species from ITS Clade 7a including two sexually functional heterothallic hybrid species detected in natural ecosystems in Taiwan. Persoonia 38: 100135. | KU517152.1 | MH620056.1 | KX251620 | KX251622 | KX251618 | KX251621 | KX251619 | KX251616 | KX251617 |
| P. fluvialis | None | JungT; BurgessTI; HuberliD; HardyGEStJ; and StukelyMJC. 2011. Phytophthora fluvialis. Persoonia 26: 146-147 (pg147). In: Crous PW; Groenewald JZ; ShivasRG;EdwardsJ;SeifertKA. 2011. Fungal Planet Description Sheets: 69-91. Persoonia. 26: 108-156. | EU593261.2 | JF701440.1 | KX251212 | KX251214 | KX251210 | KX251213 | KX251211 | KX251208 | KX251209 |
| P. foliorum | None | DonahooR; BlomquistCL; ThomasSL; MoultonJK; CookeDEL; and LamourKH. 2006. Phytophthora foliorum sp. nov.; a new species causing leaf blight of azalea. Mycol. Res. 110: 1309-1322 (pg. 1318). | MG865493.1 | EU124918.1 | KX252116 | KX252118 | KX252114 | KX252117 | KX252115 | KX252112 | KX252113 |
| P. formosa | None | Jung T Jung MH Scanu B Seress D Kovcs GM Maia C Prez-Sierra A Chang T-T Chandelier A Heungens K van Poucke K Abad-Campos P Lon M Cacciola SO and Bakonyi J.2017. Six newPhytophthoraspecies from ITS Clade 7a including two sexually functional heterothallic hybrid species detected in natural ecosystems in Taiwan. Persoonia 38: 100135. | KU517153.1 | KU899350.1 | KX251627 | KX251629 | KX251625 | KX251628 | KX251626 | KX251623 | KX251624 |
| P. fragariae | None | Hickman CJ. 1940. The red core root disease of strawberry caused by Phytophthora fragariae n. sp. J. Pomol. Hortic. Sci. 18: 89-118 (pg. 103). | MH178333.1 | AY564180.1 | KX251533, KX251540, KX251547, KX251857 | KX251535, KX251542, KX251549, KX251859 | KX251531, KX251538, KX251545, KX251855 | KX251534, KX251541, KX251548, KX251858 | KX251532, KX251539, KX251546, KX251856 | KX251529, KX251536, KX251543, KX251853 | KX251530, KX251537, KX251544, KX251854 |
| P. fragariaefolia | None | Rahman MZ; Uematsu S; Takeuchi T; Shirai K; Ishiguro Y; Suga H; and Kageyama K. 2014. Two new species; Phytophthora nagaii sp. nov. and P. fragariaefolia sp. nov.; causing serious diseases on rose and strawberry plants; respectively; in Japan. J. Gen. Plant Pathol. 80: 348-365. | MG865495.1 | MH620073.1 | KX251857 | KX251859 | KX251855 | KX251858 | KX251856 | KX251853 | KX251854 |
| P. frigida | None | MasekoB; BurgessTI; CoutinhoTA; and WingfieldMJ. 2007. Two new Phytophthora species from South Africa. Mycol. Res. 111: 1321-1338. | MG865496.1 | KF317098.1 | KX250905, KX250912, KX250919 | KX250907, KX250914, KX250921 | KX250903, KX250910, KX250917 | KX250906, KX250913, KX250920 | KX250904, KX250911, KX250918 | KX250901, KX250908, KX250915 | KX250902, KX250909, KX250916 |

| | | | | | | | | | | | |
|---|---|---|---|---|---|---|---|---|---|---|---|
| P. gallica | None | JungTand NechwatalJ. 2008. Phytophthora gallica sp. nov.; a new species from rhizosphere soil of declining oak and reed stands in France and Germany. Mycol. Res. 112: 1195-1205 (pg. 1200). | MG774933.1 | KF317112.1 | KX252593, KX252600 | KX252595, KX252602 | KX252591, KX252598 | KX252594, KX252601 | KX252592, KX252599 | KX252589, KX252596 | KX252590, KX252597 |
| P. gemini | None | Man in 't VeldWA; RosendahlKC; BrouwerH; and de CockAW. 2011. Phytophthora gemini sp. nov.; a new species isolated from the halophilic plant Zostera marina in the Netherlands. Fungal Biol. 115: 724-732. | KY769194.1 | MF326859.1 | KX251129, KX251136 | KX251131, KX251138 | KX251127, KX251134 | KX251130, KX251137 | KX251128, KX251135 | KX251125, KX251132 | KX251126, KX251133 |
| P. gibbosa | None | JungT; StukelyMJC; HardyGEStJ; WhiteMD; PaapT; DunstanWA; and BurgessTI. 2011. Multiple new Phytophthora species from ITS Clade 6 associated with natural ecosystems in Australia: evolutionary and ecological implications. Persoonia 13: 13-39. | MG865499.1 | HQ012846.1 | KX251219, KX251226 | KX251221, KX251228 | KX251217, KX251224 | KX251220, KX251227 | KX251218, KX251225 | KX251215, KX251222 | KX251216, KX251223 |
| P. glovera | None | AbadZG; Ivors KL; Gallup CA; AbadJAand Shew HD. 2011. Morphological and molecular characterization of Phytophthora glovera sp. nov. from tobacco in Brazil. Mycologia 103: 341-350. | MG865500.1 | MH620022.1 | KX250646, KX250653 | KX250648, KX250655 | KX250644, KX250651 | KX250647, KX250654 | KX250645, KX250652 | KX250642, KX250649 | KX250643, KX250650 |
| P. gonapodyides | None | Buisman; C. J. 1927. Root rots caused by Phycomycetes. Mededelingen uit het Phytopathologisch laboratorium 'Willie Commelin Scholten' 11: 1-65. | MF034094.1 | MG721475.1 | KX251233, KX251240 | KX251235, KX251242 | KX251231, KX251238 | KX251234, KX251241 | KX251232, KX251239 | KX251229, KX251236 | KX251230, KX251237 |
| P. gondwanensis | None | Shuttleworth LA; Scarlett K; Daniel R; and GuestDI. 2016.Phytophthora gondwanensis. Persoonia 35: 298-299.In CrousP; et al. 2015. Fungal Planet description sheets: 371-399. Persoonia 35: 264-327. | | | KX252607 | KX252609 | KX252605 | KX252608 | KX252606 | KX252603 | KX252604 |
| P. gregata | None | JungT; StukelyMJC; HardyGEStJ; WhiteMD; PaapT; DunstanWA; and BurgessTI. 2011. Multiple new Phytophthora species from ITS Clade 6 associated with natural ecosystems in Australia: evolutionary and ecological implications. Persoonia 13: 13-39. | MG865503.1 | MN866084.1 | KX251247, KX251254, KX251414 | KX251249, KX251256, KX251416 | KX251245, KX251252, KX251412 | KX251248, KX251255, KX251415 | KX251246, KX251253, KX251413 | KX251243, KX251250, KX251410 | KX251244, KX251251, KX251411 |
| P. hedraiandra | None | De Cock AWAM; Lvesque A; 2004. New species of Pythium and Phytophthora. Studies in Mycology 50: 481-488. | MG707790.1 | KJ567084.1 | KX250387, KX250394, KX250401 | KX250389, KX250396, KX250403 | KX250385, KX250392, KX250399 | KX250388, KX250395, KX250402 | KX250386, KX250393, KX250400 | KX250383, KX250390, KX250397 | KX250384, KX250391, KX250398 |
| P. heveae | Phytophthora palmivora var. heveae | ThompsonA. 1929. Phytophthora species in Malaya. Malayan Agric. J. 17: 53-100. | JX996048.1 | MN866088.1 | KX251115, KX251122 | KX251117, KX251124 | KX251113, KX251120 | KX251116, KX251123 | KX251114, KX251121 | KX251111, KX251118 | KX251112, KX251119 |
| P. hibernalis | None | Carne WM. 1926. A brown rot of citrus in Australia (Phytophthora hibernalis n. sp.). Journal of the Royal Society of Western Australia 12: 13-42. | MG865506.1 | AY129170.1 | KX252123, KX252130 | KX252125, KX252132 | KX252121, KX252128 | KX252124, KX252131 | KX252122, KX252129 | KX252119, KX252126 | KX252120, KX252127 |
| P. himalsilva | None | VettrainoAM; BrasierCM; BrownAV; and VanniniA. 2011. Phytophthora himalsilva sp. nov. an unusually phenotypically variable species from a remote forest in Nepal. Fungal Biol. 115: 275-287. | MG865507.1 | HM752800.1 | KX250576, KX250583 | KX250578, KX250585 | KX250574, KX250581 | KX250577, KX250584 | KX250575, KX250582 | KX250572, KX250579 | KX250573, KX250580 |
| P. humicola | None | Ko WHand Ann JP. 1985. Phytophthora humicola; a new species from soil of a citrus orchard in Taiwan. Mycologia 77: 631-636. | GU111608.1 | KM883118.1 | KX251150, KX251143 | KX251145, KX251152 | KX251141, KX251148 | KX251144, KX251151 | KX251142, KX251149 | KX251139, KX251146 | KX251140, KX251147 |
| P. hydrogena | None | Yang X; Gallegly ME; and Hong C. 2014. A high-temperature tolerant species in Clade 9 of the genus Phytophthora: P. hydrogena sp. nov. Mycologia 106: 57-65. | MG865508.1 | KC249961.1 | KX252277, KX252284, KX252291 | KX252279, KX252286, KX252293 | KX252275, KX252282, KX252289 | KX252278, KX252285, KX252292 | KX252276, KX252283, KX252290 | KX252273, KX252280, KX252287 | KX252274, KX252281, KX252288 |
| P. hydropathica | P. drechsleri II | Hong CX; Gallegly ME; Richardson PA; Kong P; Moorman GW; Lea-Cox JD; and Ross DS. 2010. Phytophthora hydropathica; a new pathogen identified from irrigation water; Rhododendron catawbiense and Kalmia latifolia. Plant Pathology 59: 913-921. | MT707339.1 | KC855138.1 | KX252298, KX252305 | KX252300, KX252307 | KX252296, KX252303 | KX252299, KX252306 | KX252297, KX252304 | KX252294, KX252301 | KX252295, KX252302 |
| P. idaei | None | KennedyDMand DuncanJM. 1995. A papillate Phytophthora species with specificity to Rubus. Mycol. Res. 99: 57-68. | MG865509.1 | AY564185.1 | EU080133, KX250408 | EU080135, KX250410 | EU080131, KX250406 | EU080134, KX250409 | EU080132, KX250407 | EU080129, KX250404 | EU080130, KX250405 |
| P. ilicis | None | BuddenhagenIWand YoungRA. 1957. A leaf twig disease of English holly caused by Phytophthora ilicis n.sp. Phytopathology 47: 95-101. | MG865510.1 | AY129172.1 | KX250940, KX250947, KX250954 | KX250942, KX250949, KX250956 | KX250938, KX250945, KX250952 | KX250941, KX250948, KX250955 | KX250939, KX250946, KX250953 | KX250936, KX250943, KX250950 | KX250937, KX250944, KX250951 |
| P. infestans | None | De BaryA. 1876. Researches into the nature of the potato fungus - Phytophthora infestans. Journal of the Royal Agricultural Society XII (IX): 240-242 (pg. 240). | MH178350.1 | MH286886.1 | KX250478, EU079629 | KX250480, EU079631 | KX250476, EU079627 | KX250479, EU079630 | KX250477, EU079628 | KX250474, EU079625 | KX250475, EU079626 |
| P. inflata | Should just be P. inflata, P. sp. 28D1 in Yang et al. | Caroselli, N.E.; Tucker, C.M. 1949. Pit canker of elm. Phytopathology. 39:481-488 | | | KX250765 | KX250767 | KX250763 | KX250766 | KX250764 | KX250761 | KX250762 |
| P. insolita | None | AnnPJand KoWH. 1980. Phytophthora insolita; a new species from Taiwan. Mycologia 72: 1180-1185. | MG865515.1 | AY564188.1 | KX252529, EU080179, EU080213 | KX252531, EU080181, EU080215 | KX252527, EU080177, EU080211 | KX252530, EU080180, EU080214 | KX252528, EU080178, EU080212 | KX252525, EU080175, EU080209 | KX252526, EU080176, EU080210 |
| P. insulinativitatica | None | Dang, QN; Pham, TQ; Arentz, F; Hardy, GEStJ; Burgess, TI. 2021. New Phytophthora species in clade 2a from the Asia-Pacific region including a re-examination of P. colocasiae and P. meadii. Mycological Progress. 20(2):111-129 | | MT583646.1 | MT583676.1 | | OK267379.1 | | | OL342764.1 | MT583631.1 |

| | | | | | | | | | | | |
|---|---|---|---|---|---|---|---|---|---|---|---|
| P. intercalaris | None | Yang X; Balci Y; Brazee NJ; Loyd AL; and Hong CX. 2015. A unique species in PhytophthoraClade 10: Phytophthora intercalaris sp. nov. recovered from stream and irrigation water in eastern United States. International Journal of Systematic and Evolutionary Microbiology 66: 845-855. | MF543341.1 | KT163332.1 | KX252614, KX252621, KX252628 | KX252616, KX252623, KX252630 | KX252612, KX252619, KX252626 | KX252615, KX252622, KX252629 | KX252613, KX252620, KX252627 | KX252610, KX252617, KX252624 | KX252611, KX252618, KX252625 |
| P. intricata | None | Jung T Jung MH Scanu B Seress D Kovcs GM Maia C Prez-Sierra A Chang T-T Chandelier A Heungens K van Poucke K Abad-Campos P Lon M Cacciola SO and Bakonyi J.2017. Six newPhytophthoraspecies from ITS Clade 7a including two sexually functional heterothallic hybrid species detected in natural ecosystems in Taiwan. Persoonia 38: 100135. | KU517155.1 | MH620059.1 | KX251634 | KX251636 | KX251632 | KX251635 | KX251633 | KX251630 | KX251631 |
| P. inundata | None | BrasierCM; Sanchez-HernandezE; and KirkSA. 2003. Phytophthora inundata sp. nov.; a part heterothallic pathogen of trees and shrubs in wet or flooded soils. Mycological Research 107: 477-484. | KJ755115.1 | HQ012860.1 | KX251157, KX251164, EU080206 | KX251159, KX251166, EU080208 | KX251155, KX251162, EU080204 | KX251158, KX251165, EU080207 | KX251156, KX251163, EU080205 | KX251153, KX251160, EU080202 | KX251154, KX251161, EU080203 |
| P. ipomoeae | None | GrnwaldNJ. 2013. Phytophthora ipomoea Flier & Grnwald; sp. nov. Fungal Planet 197. Persoonia 31: 264-265. In: Crous PW; et al. 2013. Fungal Planet description sheets: 154-213. Persoonia 31: 188-296. | KF777191.1 | HM590420.1 | EU080841, EU080834, EU080848 | EU080843, EU080836, EU080850 | EU080839, EU080832, EU080846 | EU080842, EU080835, EU080849 | EU080840, EU080833, EU080847 | EU080837, EU080830, EU080844 | EU080838, EU080831, EU080845 |
| P. iranica | None | ErshadD. 1971. Beitrag zur Kenntnis der Phytophthora Arten in Iran und ihrer Phytopathlogischen Bedeutung (Contribution to the knowledge of Phytophthora species in Iran and their phytopathogenic importance). Mitt. Biol. Bundesanst. Land Forstwirtsch. Berl. Dahlem 140: 1-90 (pgs 60-64). | FJ196753.1 | MH136913.1 | KX250443 | KX250445 | KX250441 | KX250444 | KX250442 | KX250439 | KX250440 |
| P. irrigata | Classified as P. drechsleri I by Mannon Gallegly. | Hong C; Gallegly ME; Richardson PA; Kong P;and Moorman GW. 2008. Phytophthora irrigata; a new species isolated from irrigation reservoirs and rivers in eastern United States of America. FEMS Microbiol Lett 285: 203-211. | MN513268.1 | KC733453.1 | KX252312, KX252319, KX252326 | KX252314, KX252321, KX252328 | KX252310, KX252317, KX252324 | KX252313, KX252320, KX252327 | KX252311, KX252318, KX252325 | KX252308, KX252315, KX252322 | KX252309, KX252316, KX252323 |
| P. kelmanii | None | Crous, P. W., Cowan, D. A., Maggs-Kölling, G., Yilmaz, N., Thangavel, R., Wingfield, M. J., ... & Groenewald, J. Z. (2021). Fungal planet description sheets: 1182-1283. Persoonia: Molecular Phylogeny and Evolution of Fungi, 46, 313-528. | | MT210498.1 | MT210494.1 | | OK267380.1 | | | OK533442.1 | MT210491.1 |
| P. kernoviae | None | BrasierCM; BealesPA; KirkSA; DenmanS;and RoseJ. 2005. Phytophthora kernoviae sp. nov.; an invasive pathogen causing bleeding stem lesions on forest trees and foliar necrosis of ornamentals in the UK. Mycol. Res. 109: 853-859. | MG865521.1 | KF358235.1 | EU080045, EU079649, EU080031 | KX252631, EU079651, KX252632 | EU080043, EU079647, EU080029 | EU080046, EU079650, EU080032 | EU080044, EU079648, EU080030 | EU080041, EU079645, EU080027 | EU080042, EU079646, EU080028 |
| P. kwongonina | None | Burgess TI Simamora AV White D Wiliams B Schwager M Stukely MJC and Hardy GE StJ. 2018. New species from Phytophthora Clade 6a: evidence for recent radiation. Persoonia 41: 117. | JN547636.1 | MF326846.1 | MF326876.1, MF326875.1, HQ012932.1, MK020329.1 | | MK864044.1 | | | | MF326824.1, MF326823.1, MF326822.1 |
| P. lactucae | None | BertierL; BrouwerH; de CockAWAM; CookeDEL; OlssonCHB; and HfteM. 2013. The expansion of PhytophthoraClade 8b: three new species associated with winter grown vegetable crops. Persoonia 31: 63-76. | MG865573.1 | MH620085.1 | KX252046, KX252053, KX252060 | KX252048, KX252055, KX252062 | KX252044, KX252051, KX252058 | KX252047, KX252054, KX252061 | KX252045, KX252052, KX252059 | KX252042, KX252049, KX252056 | KX252043, KX252050, KX252057 |
| P. lacustris | None | NechwatalJ;BakonyibJ;CacciolaSO; CookeDEL; JungT; NagyZA; VanniniA; VettrainoAM;and BrasierCM. 2013. The morphology; behaviour; and molecular phylogeny of Phytophthora taxon Salixsoil and its redesignation as Phytophthora lacustris sp. nov. Plant Pathology 62: 355-369. | KT383049.2 | KJ631597.2 | KX251261, KX251268, KX251275, EU080534 | KX251263, KX251270, KX251277, EU080536 | KX251259, KX251266, KX251273, EU080532 | KX251262, KX251269, KX251276, EU080535 | KX251260, KX251267, KX251274, EU080533 | KX251257, KX251264, KX251271, EU080530 | KX251258, KX251265, KX251272, EU080531 |
| P. lateralis | None | TuckerCM andMilbrathJA. 1942. Root rot of Chamaecyparis caused by a species of Phytophthora. Mycologia 34: 94-103. | HQ875390.1 | JX121293.1 | KX252137, KX252144 | KX252139, KX252146 | KX252135, KX252142 | KX252138, KX252145 | KX252136, KX252143 | KX252133, KX252140 | KX252134, KX252141 |
| P. lilii | None | Rahman MZ Uematsu S Kimishima E Kanto T Kusunoki M et al. (2015) Two plant pathogenic species of Phytophthora associated with stem blight of Easter lily and crown rot of lettuce in Japan. Mycoscience 56: 419433. | MG865523.1 | MH136918.1 | AB856794 | AB856800 | AB856788 | AB856797 | AB856791 | AB856779 | AB856782 |
| P. litchii | Previously thought to be a downy mildew | Ye, W., Wang, Y., Shen, D., Li, D., Pu, T., Jiang, Z., ... & Wang, Y. (2016). Sequencing of the litchi downy blight pathogen reveals it is a Phytophthora species with downy mildew-like characteristics. Molecular Plant-Microbe Interactions, 29(7), 573-583. | MG865524.1 | MH136919.1 | MK020333.1 | | MH359020.1 | | | MH380118.1 | OL466902.1 |
| P. litoralis | None | Jung T; Stukely MJC; Hardy GE St; White MD; Paap T; Dunstan WA; and Burgess TI. 2011. Multiple new Phytophthora species from ITS Clade 6 associated with natural ecosystems in Australia: evolutionary and ecological implications. Persoonia 13: 13-39. | MG865526.1 | HQ012866.1 | KX251282 | KX251284 | KX251280 | KX251283 | KX251281 | KX251278 | KX251279 |
| P. macilentosa | None | Yang X; Copes WE; and Hong C. 2014. Two novel species representing a new clade and cluster of Phytophthora. Fungal Biology 118: 72-82. | MG865527.1 | KF192706.1 | KX252333, KX252340, KX252347, KX252354 | KX252335, KX252342, KX252349, KX252356 | KX252331, KX252338, KX252345, KX252352 | KX252334, KX252341, KX252348, KX252355 | KX252332, KX252339, KX252346, KX252353 | KX252329, KX252336, KX252343, KX252350 | KX252330, KX252337, KX252344, KX252351 |

| | | | | | | | | | | | |
|---|---|---|---|---|---|---|---|---|---|---|---|
| P. macrochlamydospora | None | Irwin JAG. 1991. Phytophthora macrochlamydospora a new species from Australia. Mycologia 83: 517519. | MN872792.1 | MN866111.1 | EU080008 | EU080010 | EU080006 | EU080009 | EU080007 | EU080004 | EU080005 |
| P. marrasii | None | Bregant, C; Rossetto, G; Deidda, A; Maddau, L; Franceschini, A; Ionta, G; Raiola, A; Montecchio, L; Linaldeddu, BT. 2021. Phylogeny and Pathogenicity of Phytophthora Species Associated with Artichoke Crown and Root Rot and Description of Phytophthora marrasii sp. nov. Agriculture. 11(no. 873):1-14 | MZ569854.1 | OK054535.1 | | | | | | | MZ603724.1 |
| P. meadii | None | McRaeW. 1918. Phylophthora meadii n. sp. on Hevea brasiliensis. Mem. Dept. Agric. India Vol IX: 219-273. | KC247922.1 | AY564192.1 | KX250590, KX250597 | KX250592, KX250599 | KX250588, KX250595 | KX250591, KX250598 | KX250589, KX250596 | KX250586, KX250593 | KX250587, KX250594 |
| P. medicaginis | Classified as P. megasperma by Mannon Gallegly, now P. medicaginis. | HansenEMand MaxwellDP. 1991. Species of the Phytophthora megasperma-complex. Mycologia 83: 376-381(pg. 377). | AY995390.1 | KF358236.1 | KX251906, KX251913 | KX251908, KX251915 | KX251904, KX251911 | KX251907, KX251914 | KX251905, KX251912 | KX251902, KX251909 | KX251903, KX251910 |
| P. mediterranea | None | Bregant, C; Mulas, AA; Rossetto, G; Deidda, A; Maddau, L; Piras, G; Linaldeddu, BT. 2021. Phytophthora mediterranea sp. nov., a New Species Closely Related to Phytophthora cinnamomi from Nursery Plants of Myrtus communis in Italy. Forests. 12(6, no. 682):1-16 | MW892398.1 | MW900448.1 | | | | | | | MW900444.1 |
| P. megakarya | None | BrasierCMand GriffinMJ. 1979. Taxonomy of 'Phytophthora palmivora' on cocoa. T. Brit. Mycol. Soc. 72: 111-143 (pg. 137). | MH401202.1 | AY564193.1 | KX251031, KX251038, KX251045 | KX251033, KX251040, KX251047 | KX251029, KX251036, KX251043 | KX251032, KX251039, KX251046 | KX251030, KX251037, KX251044 | KX251027, KX251034, KX251041 | KX251028, KX251035, KX251042 |
| P. megasperma | Phytophthora megasperma var. megasperma | DrechslerCA. 1931. A crown rot of hollyhock caused by Phytophthora megasperma n. sp. Journal of the Washington Academy of Science 21: 513-526. | KJ755116.1 | L04457.1 | KX251421, KX251428 | KX251423, KX251430 | KX251419, KX251426 | KX251422, KX251429 | KX251420, KX251427 | KX251417, KX251424 | KX251418, KX251425 |
| P. mekongensis | P. aff. meadii[1] | Cacciola SO, La Spada F, Hoa NV, Jung MH, and Scanu B. 2017. *Phytophthora mekongensis* Cacciola & N.V. Hoa, *sp. nov.* Fungal Planet description sheet: 615. Persoonia 38: 363. In Crous et al. 2017. Fungal Planet description sheets: 558–624. Persoonia 38: 240–384. | LC595792.1 | LC595927.1 | | | | LC595822.1 | | | LC595870.1 |
| P. melonis | None | KatsuraK. 1976. Two new species of Phytophthora causing damping-off of cucumber and trunk rot of chestnut. Trans. Mycol. Soc. Jpn. 17: 238-242. | MK386561.1 | EF372621.1 | KX251697, KX251704, KX251711 | KX251699, KX251706, KX251713 | KX251695, KX251702, KX251709 | KX251698, KX251705, KX251712 | KX251696, KX251703, KX251710 | KX251693, KX251700, KX251707 | KX251694, KX251701, KX251708 |
| P. mengei | None | Hong CX; Gallegly ME; Browne GT; Bhat RG;Richardson PA; and Kong P. 2009. The avocado subgroup of Phytophthora citricola constitutes a distinct species; Phytophthora mengei sp. nov. Mycologia 101: 833-840. | MG865539.1 | GU191299.1 | KX250660, KX250667 | KX250662, KX250669 | KX250658, KX250665 | KX250661, KX250668 | KX250659, KX250666 | KX250656, KX250663 | KX250657, KX250664 |
| P. mexicana | Treated separately in Yang et al. and by Mannon Gallegly but may be a synonym of P. capsici. | Hotson JW and Hartge L. 1923. A disease of tomato caused by Phytophthora mexicana sp. nov. Phytopathology 13: 520-531 (pg 520). | GU111665.1 | MH620024.1 | KX250674 | KX250676 | KX250672 | KX250675 | KX250673 | KX250670 | KX250671 |
| P. mirabilis | None | Galindo AJ and Hohl HR. 1985. Phytophthora mirabilis a new species of Phytophthora. Sydowia 38: 87-96 (p 95). | MH178350.1 | HM590421.1 | KX250485, KX250492 | KX250487, KX250494 | KX250483, KX250490 | KX250486, KX250493 | KX250484, KX250491 | KX250481, KX250488 | KX250482, KX250489 |
| P. mississippiae | None | Yang X; Copes WE; and Hong C. 2013. Phytophthora mississippiae sp. nov.; a new species recovered from irrigation reservoirs at a plant nursery in Mississippi. J Plant Pathol Microb 4: 1-7. | KP780453.1 | KF112858.1 | KX251295, KX251302, KX251309, KX251316 | KX251297, KX251304, KX251311, KX251318 | KX251293, KX251300, KX251307, KX251314 | KX251296, KX251303, KX251310, KX251317 | KX251294, KX251301, KX251308, KX251315 | KX251291, KX251298, KX251305, KX251312 | KX251292, KX251299, KX251306, KX251313 |
| P. morindae | None | Nelson S andAbad ZG. 2010. Phytophthora morindae; a new species causing black flag on noni (Morinda citrifolia L) in Hawaii. Mycologia 102: 122-134. | MG865543.1 | KT183050.1 | KX252637 | KX252639 | KX252635 | KX252638 | KX252636 | KX252633 | KX252634 |
| P. moyootj | None | BurgessTIand StukelyMJC. 2014. Phytophthora moyootj Persoonia 33: 278279. In CrousP et al. 2014. Fungal Planet description sheets: 281319. Persoonia 33: 212289. | KJ372256.1 | KM883134.1 | KJ396730.1, KJ396729.1, KJ396728.1, MK020349.1, KM883162.1, KM883156.1, KM883150.1 | | MK864046.1 | KP004501.1, KP004500.1, KP004499.1 | | | KJ372303.1, KJ372302.1, KJ372301.1, KM883106.1, KM883100.1, KM883094.1 |
| P. multibullata | None | Dang, QN; Pham, TQ; Arentz, F; Hardy, GEStJ; Burgess, TI. 2021. New Phytophthora species in clade 2a from the Asia-Pacific region including a re-examination of P. colocasiae and P. meadii. Mycological Progress. 20(2):111-129 | MT568655.1 | MT583658.1 | MT583684.1 | | OK267381.1 | | | OK533443.1 | MT583643.1 |
| P. multivesiculata | None | IlievaE; Man in 't VeldWA; Veenbaas-RijksW; and PietersR. 1998.Phytophthora multivesiculata; a new species causing rot in Cymbidium. Eur. J. Plant Pathol. 104: 677-684. | DQ835678.1 | AY564195.1 | EU080069, KX250926 | EU080071, KX250928 | EU080067, KX250924 | EU080070, KX250927 | EU080068, KX250925 | EU080065, KX250922 | EU080066, KX250923 |

| | | | | | | | | | | | |
|---|---|---|---|---|---|---|---|---|---|---|---|
| P. multivora | None | ScottPM;BurgessTI;BarberPA;ShearerBL;StukelyMJC;HardyGEStJ; andJungT.2009. Phytophthora multivora sp. nov.; a new species recovered from declining Eucalyptus; Banksia; Agonis; and other plant species in Western Australia. Persoonia 22: 1-13. | MT355394.1 | FJ237503.1 | KX250779 | KX250781 | KX250777 | KX250780 | KX250778 | KX250775 | KX250776 |
| P. nagaii | None | Rahman MZ; Uematsu S; Takeuchi T; Shirai K; Ishiguro Y; Suga H; and Kageyama K. 2014. Two new species; Phytophthora nagaii sp. nov. and P. fragariaefolia sp. nov.; causing serious diseases on rose and strawberry plants; respectively; in Japan. J. Gen. Plant Pathol. 80: 348-365. | MN270927.1 | MH620074.1 | KX251864 | KX251866 | KX251862 | KX251865 | KX251863 | KX251860 | KX251861 |
| P. nemorosa | None | Hansen EM; Reeser PW; Davidson JM; Garbelotto M; Ivors K; Douhan L; and Rizzo DM. 2003. Phytophthora nemorosa; a new species causing cankers and leaf blight of forest trees in California and Oregon; U.S.A. Mycotaxon 88: 129-138 (pg 131). | MG696505.1 | KF317104.1 | KX250961,<br>KX250968 | KX250963,<br>KX250970 | KX250959,<br>KX250966 | KX250962,<br>KX250969 | KX250960,<br>KX250967 | KX250957,<br>KX250964 | KX250958,<br>KX250965 |
| P. nicotianae | P. parasitica, P. parasitica var nicotianae, P. melongenae, P. tabaci, P. terrestris, P. manoana | Breda de Haan van. 1896. De bibitziekte in de Deli-tabak veroorzaakt door Phytophthora nicotianae. Meded. uit 's Lands Plantentuin Coll. XV (met plaat) Batavia 15: 1-107 (pg 57). | JX978447.1 | KY851301.1 | KX250513,<br>KX250520,<br>EU079966,<br>EU080507 | KX250515,<br>KX250522,<br>EU079968,<br>EU080509 | KX250511,<br>KX250518,<br>EU079964,<br>EU080505 | KX250514,<br>KX250521,<br>EU079967,<br>EU080508 | KX250512,<br>KX250519,<br>EU079965,<br>EU080506 | KX250509,<br>KX250516,<br>EU079962,<br>EU080503 | KX250510,<br>KX250517,<br>EU079963,<br>EU080504 |
| P. niederhauseri | P. niederhauserii, P. niederhauseria | Abad ZG; Abad JA; Cacciola SO; Pane A; Faedda R; Moralejo E; Prez-Sierra A; Abad-Campos P; Alvarez-Bernaola LA; Bakonyi J; Jzsa A; Herrero MA; Burgess TI; Cunnington JH; Smith IW; Balci Y; Blomquist C; Henricot B; Denton G; Spies C; Mcleod A; Belbahri L; Cooke D; Kageyama K; Uematsu S; Kurbetli I;and Degrmenci K. 2014. Phytophthora niederhauserii sp. nov.; a polyphagous species associated with ornamentals; fruit trees; and native plants in 13 countries. Mycologia 106 (3): 431-447. | MK330701.1 | GU477617.1 | KX251718,<br>KX251725,<br>KX251732 | KX251720,<br>KX251727,<br>KX251734 | KX251716,<br>KX251723,<br>KX251730 | KX251719,<br>KX251726,<br>KX251733 | KX251717,<br>KX251724,<br>KX251731 | KX251714,<br>KX251721,<br>KX251728 | KX251715,<br>KX251722,<br>KX251729 |
| P. obscura | P. drechsleri III | GrnwaldNJ; WerresS; GossEM; TaylorCR; and FielandVJ. 2012. Phytophthora obscura sp. nov.; a new species of the novel Phytophthora subclade 8d. Plant Pathol. 61: 610-622. | MN228692.1 | HQ917879.1 | KX252179,<br>KX252186,<br>KX252193 | KX252181,<br>KX252188,<br>KX252195 | KX252177,<br>KX252184,<br>KX252191 | KX252180,<br>KX252187,<br>KX252194 | KX252178,<br>KX252185,<br>KX252192 | KX252175,<br>KX252182,<br>KX252189 | KX252176,<br>KX252183,<br>KX252190 |
| P. occultans | None | Man In 't Veld WA; Rosendahl KC; van Rijswick PC; Meffert JP;Westenberg M; van de Vossenberg BT; Denton G; and van Kuik FA. 2015. Phytophthora terminalis sp. nov. and Phytophthora occultans sp. nov.; two invasive pathogens of ornamental plants in Europe. Mycologia 107: 54-65. | MH141415.1 | KU891804.1 | KX250604 | KX250606 | KX250602 | KX250605 | KX250603 | KX250600 | KX250601 |
| P. oleae | None | Ruano-Rosa D Schena L Agosteo GE Magnano di San Lio G and Cacciola SO. 2018.Phytophthora oleae sp. nov. causing fruit rot of olive in southern Italy. Plant Pathology 67: 13621373. | KY982934.1 | MF083576.1 | | | | | | | |
| P. oreophila | None | Khaliq I. Hardy G. E. McDougall K. L. & Burgess T. I. (2018 November 01). Phytophthora species isolated from alpine and sub-alpine regions of Australia including the description of two new species; Phytophthora cacuminis sp. nov and Phytophthora oreophila sp. nov. Retrieved September 10 2020 from https://www.sciencedirect.com/science/article/pii/S1878614618302332 | MG542976.1 | MG543002.1 | MG543025.1 | | | | | | MG543037.1 |
| P. ornamentata | None | Scanu B; Linaldeddu BT; Deidda A; and Jung T. 2015. Diversity of Phytophthoraspecies from declining Mediterranean maquis vegetation; including two new species; Phytophthora crassamura and P. ornamentata sp. nov. PLoS ONE 10 (12): e0143234. Doi:10.1371/journal.pone.0143234 | MN959765.1 | KP863486.1 | KX251323,<br>KX251330 | KX251325,<br>KX251332 | KX251321,<br>KX251328 | KX251324,<br>KX251331 | KX251322,<br>KX251329 | KX251319,<br>KX251326 | KX251320,<br>KX251327 |
| P. pachypleura | None | Henricot B Perez Sierra A Jung T (2014) Phytophthora pachypleura sp. nov. a new species causing root rot of Aucuba japonica and other ornamentals in the United Kingdom. Plant Pathology 63: 10951109. | MG865558.1 | MH620028.1 | KX250786,<br>KX250793,<br>KX250800 | KX250788,<br>KX250795,<br>KX250802 | KX250784,<br>KX250791,<br>KX250798 | KX250787,<br>KX250794,<br>KX250801 | KX250785,<br>KX250792,<br>KX250799 | KX250782,<br>KX250789,<br>KX250796 | KX250783,<br>KX250790,<br>KX250797 |
| P. palmivora | P. faberi, P. omnivora, P. arecae | Butler EJ. 1918-1919. Report of the imperial mycologist; Science Report Institute Pusa. 82 pp. | MF370572.1 | AY564197.1 | KX251052,<br>KX251059 | KX251054,<br>KX251061 | KX251050,<br>KX251057 | KX251053,<br>KX251060 | KX251051,<br>KX251058 | KX251048,<br>KX251055 | KX251049,<br>KX251056 |
| P. parsiana | None | Mostowfizadeh-GhalamfarsaR;CookeDEL; and BanihashemiZ. 2008. Phytophthora parsiana sp. nov.; a new high-temperature tolerant species. Mycological Research 112: 783-794. | MT232850.1 | KC733455.1 | KX252361 | KX252363 | KX252359 | KX252362 | KX252360 | KX252357 | KX252358 |
| P. parvispora | P. cinnamomi var. parvispora | ScanuB; HunterGC; LinaldedduBT; FranceschiniA; MaddauL; JungT; and DenmanS. 2014. A taxonomic re-evaluation reveals that Phytophthora cinnamomi and Phytophthora cinnamomi var. parvispora are separate species. Forest Pathol. 44: 1-20. | | MN866092.1 | KX251822,<br>KX251829,<br>KX251836,<br>KX251843 | KX251824,<br>KX251831,<br>KX251838,<br>KX251845 | KX251820,<br>KX251827,<br>KX251834,<br>KX251841 | KX251823,<br>KX251830,<br>KX251837,<br>KX251844 | KX251821,<br>KX251828,<br>KX251835,<br>KX251842 | KX251818,<br>KX251825,<br>KX251832,<br>KX251839 | KX251819,<br>KX251826,<br>KX251833,<br>KX251840 |
| P. personensis | P. sp. personii[3] | Crous PW, Wingfield MJ, Chooi YH, et al. Fungal Planet description sheets: 1042-1111. *Persoonia* . 2020;44:301-459. doi:10.3767/persoonia.2020.44.11 | EU301169.2 | MF441671.1 | EU080316 | EU080318 | EU080314 | EU080317 | EU080315 | EU080312 | EU080313 |

| | | | | | | | | | | | |
|---|---|---|---|---|---|---|---|---|---|---|---|
| P. phaseoli | None | Thaxter R. 1889. A new American Phytophthora. Bot. Gaz. 14: 273-274. Phytophthora phaseoli. | MG865564.1 | HM590418.1 | KX250499, KX250506, EU080752, EU080765 | KX250501, KX250508, EU080754, EU080767 | KX250497, KX250504, EU080750, EU080763 | KX250500, KX250507, EU080753, EU080766 | KX250498, KX250505, EU080751, EU080764 | KX250495, KX250502, EU080748, EU080761 | KX250496, KX250503, EU080749, EU080762 |
| P. pini | P. citricola I | Leonian LH. 1925. Physiological studies on the genus Phytophthora. American Journal of Botany 12: 444-498. | MT597899.1 | KJ631589.2 | KX250807, KX250814, KX251337, KX251344 | KX250809, KX250816, KX251339, KX251346 | KX250805, KX250812, KX251335, KX251342 | KX250808, KX250815, KX251338, KX251345 | KX250806, KX250813, KX251336, KX251343 | KX250803, KX250810, KX251333, KX251340 | KX250804, KX250811, KX251334, KX251341 |
| P. pinifolia | None | DurnA; GryzenhoutM; SlippersB; AhumadaR; RotellaA; FloresF; WingfieldBD; and WingfieldMJ. 2008. Phytophthora pinifolia sp. nov. associated with a serious needle disease of Pinus radiata in Chile. Plant Pathol. 57: 715-727. | KJ755114.1 | GU799673.1 | KX251337, KX251344 | KX251339, KX251346 | KX251335, KX251342 | KX251338, KX251345 | KX251336, KX251343 | KX251333, KX251340 | KX251334, KX251341 |
| P. pisi | None | Heyman F; Blair JE; Persson L; and Wikstrm M. 2013. Root rot of pea and faba bean in southern Sweden caused by Phytophthora pisi sp. nov. Plant Dis. 97: 461-471. | MG865567.1 | MH620066.1 | KX251739 | KX251741 | KX251737 | KX251740 | KX251738 | KX251735 | KX251736 |
| P. pistaciae | None | MirabolfathyM; CookeDEL; DuncanJM; WilliamsNA; ErshadD; and AlizadehA. 2001. Phytophthora pistaciae sp. nov. and P. melonis: the principal causes of pistachio gummosis in Iran. Mycol. Res. 105: 1166-1175. | MT328703.1 | MH620067.1 | KX251752, KX251759 | KX251754, KX251761 | KX251750, KX251757 | KX251753, KX251760 | KX251751, KX251758 | KX251748, KX251755 | KX251749, KX251756 |
| P. plurivora | P. citricola II | JungT andBurgessTI. 2009. Re-evaluation of Phytophthora citricola isolates from multiple woody hosts in Europe and North America reveals a new species; Phytophthora plurivora sp. nov. Persoonia 22: 95-110. | KF963047.1 | KJ631582.2 | KX250821, KX250828, KX250835 | KX250823, KX250830, KX250837 | KX250819, KX250826, KX250833 | KX250822, KX250829, KX250836 | KX250820, KX250827, KX250834 | KX250817, KX250824, KX250831 | KX250818, KX250825, KX250832 |
| P. pluvialis | None | Reeser P; Sutton W; and Hansen E. 2013. Phytophthora pluvialis; a new species from mixed tanoak-Douglas fir forests of western Oregon; U.S.A. North American Fungi 8: 1-8. | MG696455.1 | MH620033.1 | KX250975 | KX250977 | KX250973 | KX250976 | KX250974 | KX250971 | KX250972 |
| P. podocarpi | None | Dobbie, K; Scott, P; Taylor, P; Panda, P; Sen, D; Dick, M; McDougal, R. 2022. Phytophthora podocarpi sp. nov. from Diseased Needles and Shoots of Podocarpus in New Zealand. Forests. 13(2, no. 214):1-13 | LGSN01002589.1 | LGSN01000280.1 | LGSN01000824.1 | LGSN01000163.1 | LGSN01003884.1 | LGSN01002589.1 | LGSN01000006.1 | LGSN01000038.1 | LGSN01000928.1 |
| P. polonica | None | BelbahriL;MoralejoE;CalminG;OszakoT;Garciia; JA;DescalsE;LefortF. 2006. Phytophthora polonica; a new species isolated from declining Alnus glutinosa in Poland. FEMS Microbiology Letters 261: 165-174. | MG753543.1 | KT946598.1 | KX252536, KX252543, EU080260 | KX252538, KX252545, EU080262 | KX252534, KX252541, EU080258 | KX252537, KX252544, EU080261 | KX252535, KX252542, EU080259 | KX252532, KX252539, EU080256 | KX252533, KX252540, KX252546 |
| P. porri | Referred to as P. aff. brassicae-2[1] by Yang et al. | FoisterCE. 1931. The white tip disease of leeks and its causal fungus; Phytophthora porri n. sp. Transactions of the Botanical Society of Edinburgh. 4: 257-281. | AB688405.1 | MH136961.1 | EU079884 | EU079886 | EU079882 | EU079885 | EU079883 | EU079880 | EU079881 |
| P. primulae | None | TomlinsonJA. 1952. Brown core root rot of Primula caused by Phytophthora primulae n. sp. T. Brit. Mycol. Soc. 35: 221-235. | MG865571.1 | KF358238.1 | KX252067, KX252074 | KX252069, KX252076 | KX252065, KX252072 | KX252068, KX252075 | KX252066, KX252073 | KX252063, KX252070 | KX252064, KX252071 |
| P. prodigiosa | P. insolita PF6e | Cacciola SO, Aloi F, Tri VM, Jung T, and Schena L. 2017. *Phytophthora prodigiosa* Cacciola & Tri. Fungal Planet Description Sheet: 616. Persoonia 38: 365. In: Crous et. al. 2017. Fungal Planet Description Sheets: 558–624. Persoonia 38: 240–384. | LC595799.1 | LC595937.1 | | | | LC595830.1 | | | LC595880.1 |
| P. pseudocryptogea | Confused with P. cryptogea | Safaiefarahani B Mostowfizadeh-Ghalamfarsa R Hardy GEStJ and Burgess TI. 2015. Re-evaluation of the Phytophthora cryptogea species complex and the description of a new species Phytophthora pseudocryptogea sp. nov. Mycol Progress 14: 112. | MG587716.1 | MN866101.1 | EU080630 | | EU080628 | EU080631 | EU080629 | EU080626 | EU080627 |
| P. pseudolactucae | None | Rahman MZ Uematsu S Kimishima E Kanto T Kusunoki M Motohashi K Ishiguro Y Suga H and Kageyama K. 2015. Two plant pathogenic species of Phytophthora associated with stem blight of Easter lily and crown rot of lettuce in Japan. Mycoscience 56: 419433. | MG865573.1 | MH136965.1 | MK020381.1 | | | | | MH380160.1 | |

| | | | | | | | | | | | |
|---|---|---|---|---|---|---|---|---|---|---|---|
| P. pseudopolonica | Not sure if this is truly distinct from P. polonica. | Li W Zhao W and Hua W. 2017. Phytophthora pseudopolonica sp. nov. a new species recovered from stream water in subtropical forests of China. Int. J. Syst. Evol. Microbiol. 67: 36663675. | KY707109.1 | | | | KY787211.1,<br>KY787210.1,<br>KY787209.1,<br>KY787208.1,<br>KY787207.1,<br>KY787206.1,<br>KY787205.1,<br>KY787204.1,<br>KY787203.1,<br>KY787202.1,<br>KY787201.1,<br>KY787200.1,<br>KY787199.1,<br>KY787198.1,<br>KY787197.1,<br>KY787196.1,<br>KY787195.1,<br>KY787194.1 | | | | KY707104.1,<br>KY707103.1,<br>KY707102.1,<br>KY707101.1,<br>KY707100.1,<br>KY707099.1,<br>KY707098.1,<br>KY707097.1,<br>KY707096.1,<br>KY707095.1,<br>KY707094.1,<br>KY707093.1,<br>KY707092.1,<br>KY707091.1,<br>KY707090.1,<br>KY707089.1,<br>KY707088.1,<br>KY707087.1 |
| P. pseudorosacearum | None | Burgess TI Simamora AV White D Wiliams B Schwager M Stukely MJC and Hardy GE StJ. 2018. New species fromPhytophthoraClade 6a: evidence for recent radiation. Persoonia 41: 117. | | MF326857.1 | MF326878.1,<br>MF326877.1,<br>HQ012931.1,<br>MK020378.1 | | MK864055.1 | | | | MN207277.1,<br>MF326827.1,<br>MF326826.1,<br>MF326825.1 |
| P. pseudosyringae | None | JungT;NechwatalJ;CookeDE;HartmannG;BlaschkeM;OsswaldWF;DuncanJM; andDelatourC. 2003. Phytophthora pseudosyringae sp. nov.; a new species causing root and collar rot of deciduous tree species in Europe. Mycological Research 107: 772-789. | MG696525.1 | KJ458943.1 | KX250982,<br>KX250989 | KX250984,<br>KX250991 | KX250980,<br>KX250987 | KX250983,<br>KX250990 | KX250981,<br>KX250988 | KX250978,<br>KX250985 | KX250979,<br>KX250986 |
| P. pseudotsugae | None | Hamm PB andHansen EM. 1983. Phytophthora pseudotsugae; a new species causing root rot of Douglas fir. Canadian Journal of Botany 61: 2630-2626. | MG685785.1 | AY129167.1 | EU080430 | EU080432 | EU080428 | EU080431 | EU080429 | EU080426 | EU080427 |
| P. psychrophila | None | JungT;HansenEM;WintonL;OsswaldW;and DelatourC.2002. Three new species of Phytophthora from European oak forests. Mycological Research 106: 397-411. | MG865576.1 | | KX250996,<br>KX251003 | KX250998,<br>KX251005 | KX250994,<br>KX251001 | KX250997,<br>KX251004 | KX250995,<br>KX251002 | KX250992,<br>KX250999 | KX250993,<br>KX251000 |
| P. quercetorum | None | BalciY;BalciS;BlairJE;ParkSY;KangS; andMacDonaldWL.2008.Phytophthora quercetorum sp. nov.; a novel species isolated from eastern and north-central U.S. oak forests. Mycological Research 112: 906-916 (pp. 911). | EF539174.1 | KX759520.1 | KX251066,<br>KX251073 | KX251068,<br>KX251075 | KX251064,<br>KX251071 | KX251067,<br>KX251074 | KX251065,<br>KX251072 | KX251062,<br>KX251069 | KX251063,<br>KX251070 |
| P. quercina | None | JungT; CookeDEL; BlaschkeH; DuncanJM; and OswaldW. 1999. Phytophthora quercina sp. nov.; causing root rot of European oaks. Mycol. Res. 103:785-798. | KJ755112.1 | MG701998.1 | KX252651,<br>KX252658,<br>KX252665 | KX252653,<br>KX252660,<br>KX252667 | KX252649,<br>KX252656,<br>KX252663 | KX252652,<br>KX252659,<br>KX252666 | KX252650,<br>KX252657,<br>KX252664 | KX252647,<br>KX252654,<br>KX252661 | KX252648,<br>KX252655,<br>KX252662 |
| P. quininea | None | CrandallBS. 1947. A new Phytophthora causing root and collar rot of Cinchona in Peru. Mycologia 39: 218-223. | MG865580.1 | AY564200.1 | EU079805 | EU079807 | EU079804 | EU079806 | KX252524 | EU079802 | EU079803 |
| P. ramorum | None | WerresS; MarwitzR; Man in 't VeldWA; de CockAWAM; BonantsPJM; de WeerdtM; ThemannK; IlievaE; and BaayenRP. 2001. Phytophthora ramorum sp. nov.; a new pathogen on Rhododendron and Viburnum. Mycol. Res. 105: 1155-1165. | MT031975.1 | MN861126.1 | KX252151,<br>KX252158 | KX252153,<br>KX252160 | KX252149,<br>KX252156 | KX252152,<br>KX252159 | KX252150,<br>KX252157 | KX252147,<br>KX252154 | KX252148,<br>KX252155 |
| P. rhizophorae | P. sp. rhizophorae[3] | Marano Jesus AL & Pires-Zottar. 2016.Phytophthora rhizophorae.Fungal Diversity 78: 196. in LiGJ HydeKD and ZhaoRL. et al. Fungal Diversity (2016) 78: 1. https://doi.org/10.1007/s13225-016-0366-9 | KT886032.1 | | | | | KT886028.1 | | | |
| P. richardiae | Phytophthora cryptogea var. richardiae | BuismanCJ. 1927. Root rots caused by Phycomycetes. Mededeelingen van bet Phytopathologisch Laboratorium "Willie Commelin Scholten"; Baarn; 11: 1-51. | MK496521.1 | AY564201.1 | KX251920,<br>KX251927,<br>EU080500 | KX251922,<br>KX251929,<br>EU080502 | KX251918,<br>KX251925,<br>EU080498 | KX251921,<br>KX251928,<br>EU080501 | KX251919,<br>KX251926,<br>EU080499 | KX251916,<br>KX251923,<br>EU080496 | KX251917,<br>KX251924,<br>EU080497 |
| P. riparia | None | Hansen EM; Reeser PW; and Sutton W. 2012. Phytophthora borealis and Phytophthora riparia; new species in Phytophthora ITS Clade 6. Mycologia 104: 1133-1142. | MG696530.1 | JQ626629.1 | KX251351 | KX251353 | KX251349 | KX251352 | KX251350 | KX251347 | KX251348 |
| P. rosacearum | P. megasperma var rosacearum, P. megasperma III | Hansen EM; Wilcox WF; Reeser PW; and Sutton W. 2009. Phytophthora rosacearum and P. sansomeana; new species segregated from the Phytophthora megasperma "complex". Mycologia 101: 129-135. | KU211080.1 | HQ012882.1 | KX251435,<br>KX251442,<br>KX251449 | KX251437,<br>KX251444,<br>KX251451 | KX251433,<br>KX251440,<br>KX251447 | KX251436,<br>KX251443,<br>KX251450 | KX251434,<br>KX251441,<br>KX251448 | KX251431,<br>KX251438,<br>KX251445 | KX251432,<br>KX251439,<br>KX251446 |
| P. rubi | P. fragariae var. rubi | Man in 't VeldWA. 2007. Gene flow analysis demonstrates that Phytophthora fragariae var. rubi constitutes a distinct species; Phytophthora rubi comb. nov. Mycologia 99: 222-226. | KJ755094.1 | KU899309.1 | KX251554,<br>KX251561,<br>KX251568 | KX251556,<br>KX251563,<br>KX251570 | KX251552,<br>KX251559,<br>KX251566 | KX251555,<br>KX251562,<br>KX251569 | KX251553,<br>KX251560,<br>KX251567 | KX251550,<br>KX251557,<br>KX251564 | KX251551,<br>KX251558,<br>KX251565 |
| P. sansomeana | None | Hansen EM; Wilcox WF; Reeser PW; and Sutton W. 2009. Phytophthora rosacearum and P. sansomeana; new species segregated from the Phytophthora megasperma "complex". Mycologia 101: 129-135. | MK794708.1 | NC045089.1 | KX251934,<br>KX251941,<br>KX251948 | KX251936,<br>KX251943,<br>KX251950 | KX251932,<br>KX251939,<br>KX251946 | KX251935,<br>KX251942,<br>KX251949 | KX251933,<br>KX251940,<br>KX251947 | KX251930,<br>KX251937,<br>KX251944 | KX251931,<br>KX251938,<br>KX251945 |

| | | | | | | | | | | | |
|---|---|---|---|---|---|---|---|---|---|---|---|
| P. siskiyouensis | None | ReeserPW;HansenEM;and SuttonW. 2008. Phytophthora siskiyouensis; a new species from soil; water; myrtlewood (Umbellularia californica); and tanoak (Lithocarpus densiflorus) in southwestern Oregon. Mycologia 99: 639-643. | MG696500.1 | KF317102.1 | KX250681, KX250688 | KX250683, KX250690 | KX250679, KX250686 | KX250682, KX250689 | KX250680, KX250687 | KX250677, KX250684 | KX250678, KX250685 |
| P. sojae | P. megasperma var. sojae, P. megasperma f.sp. Glycinea | KaufmannMJ andGerdemannJW. 1958. Root and stem rot of soybean caused by Phytophthora sojae n. sp. Phytopathology 48: 201-208. | AB217685.1 | DQ832717.1 | KX251766, KX251773 | KX251768, KX251775 | KX251764, KX251771 | KX251767, KX251774 | KX251765, KX251772 | KX251762, KX251769 | KX251763, KX251770 |
| P. sp. awatangi[3] | No formal description. Informally described in Dang et al. 2021. | Dang, QN; Pham, TQ; Arentz, F; Hardy, GEStJ; Burgess, TI. 2021. New Phytophthora species in clade 2a from the Asia-Pacific region including a re-examination of P. colocasiae and P. meadii. Mycological Progress. 20(2):111-129 | MT568650.1 | MT583653.1 | MT583679.1 | | | | | | MT583642.1 |
| P. sp. brasiliensis capsici[3] | No formal description. World Phytophthora Collection P0630. | Oudemans P Coffey MD (1991) A revised systematics of twelve papillate Phytophthora species based on isozyme analysis. Mycological Research 95: 10251046. | | | EU080423 | EU080425 | EU080421 | EU080424 | EU080422 | EU080419 | EU080420 |
| P. sp. canalensis[3] | Provisional name. World Phytophthora Collection P10456. | Yang, X., Tyler, B. M., & Hong, C. (2017). An expanded phylogeny for the genus Phytophthora. IMA fungus, 8(2), 355-384. | | | EU079573 | n/a | EU079571 | KX251380 | KX251378 | KX251375 | KX251376 |
| P. sp. cuyabensis[3] | Collected by Michael Coffey in 1993, not formally described. World Phytophthora Collection P8213. | Public Data Portal - Specimen Record. (n.d.). Retrieved September 17 2020 fromhttp://v3.boldsystems.org/index.php/Public. RecordView?processid=PHYTO230-10 | FJ801990.1 | GU594826.1 | EU080668 | EU080331 | EU080666 | EU080669 | EU080667 | EU080664 | EU080665 |
| P. sp. germisporangia[3] | No formal description. Informally described in Dang et al. 2021. | Dang, QN; Pham, TQ; Arentz, F; Hardy, GEStJ; Burgess, TI. 2021. New Phytophthora species in clade 2a from the Asia-Pacific region including a re-examination of P. colocasiae and P. meadii. Mycological Progress. 20(2):111-129 | MT568649.1 | MT583657.1 | MT583678.1 | | | | | | MT583641.1 |
| P. sp. kelmania[3] | Classified as P. kelmania and P. cryptogea by Mannon Gallegly. A different species has been named P. kelmanii, this is not the same isolate. | McKeever K. M. & Chastagner G. (2016 March 16). A Survey of Phytophthora spp. Associated with Abies in U.S. Christmas Tree Farms. Retrieved September 15 2020 from https://apsjournals.apsnet.org/doi/full/10.1094/PDIS-08-15-0939-RE | | MN866075.1 | KX251990, EU079609 | KX251992, EU079611 | KX251988, EU079607 | KX251991, EU079610 | KX251989, EU079608 | KX251986, EU079605 | KX251987, EU079606 |
| P. sp. lagoariana[3] | Collected by Michael Coffey in 1993, not formally described. World Phytophthora Collection P8220. | Public Data Portal - Specimen Record. (n.d.). Retrieved September 17 2020 from http://v3.boldsystems.org/index.php/Public. RecordView?processid=PHYTO232-10 | MT232839.1 | GU594812.1 | EU080361, KX252507, EU080368 | EU080363, KX252509, EU080370 | EU080359, KX252505, EU080366 | EU080362, KX252508, EU080369 | EU080360, KX252506, EU080367 | EU080358, KX252503, EU080364 | KX252502, KX252504, EU080365 |
| P. sp. novaeguineae[3] | Not formally described, World Phytophthora Collection accession P1256. Previously classified as P. katsurae and castaneae. | Unknown | | | | | | JF273228.1, JF273224.1 | | | |
| P. sp. sulawesiensis[3] | World Phytophthora Collection accession P6306. | Unknown | EF590257.1 | HQ261458.1 | EU080348 | EU080350 | EU080346 | EU080349 | EU080347 | EU080345 | n/a |
| P. sp. taxon walnut[3] | No formal description. | Brasier CM Cooke DEL Duncan JM Hansen EM. (2003a) Multiple new phenotypic taxa from trees and riparian ecosystems in Phytophthora gonapodyidesP. megasperma ITS Clade 6 which tend to be high-temperature tolerant and either inbreeding or sterile. Mycological Research 107: 277290. | MT065854.1 | MH620040.1 | KX251456, KX251463 | KX251458, KX251465 | KX251454, KX251461 | KX251457, KX251464 | KX251455, KX251462 | KX251452, KX251459 | KX251453, KX251460 |
| P. sp.22J5 gregata like[3] | Classified by Mannon Gallegly as not Phtophthora. Deposited in American Type Culture Collection as P. erythroseptica. | Yang, X., Tyler, B. M., & Hong, C. (2017). An expanded phylogeny for the genus Phytophthora. IMA fungus, 8(2), 355-384. | | | KX251414 | KX251416 | KX251412 | KX251415 | KX251413 | KX251410 | KX251411 |
| P. sp.27D9 citricolaVIII[3] | No formal description. | Yang, X., Tyler, B. M., & Hong, C. (2017). An expanded phylogeny for the genus Phytophthora. IMA fungus, 8(2), 355-384. | | | KX250842 | KX250844 | KX250840 | KX250843 | KX250841 | KX250838 | KX250839 |
| P. sp.35G4[3] | No formal description. | Yang, X., Tyler, B. M., & Hong, C. (2017). An expanded phylogeny for the genus Phytophthora. IMA fungus, 8(2), 355-384. | | | KX252485 | KX252487 | KX252483 | KX252486 | KX252484 | KX252481 | KX252482 |
| P. sp.38D9[3] | No formal description. | Yang, X., Tyler, B. M., & Hong, C. (2017). An expanded phylogeny for the genus Phytophthora. IMA fungus, 8(2), 355-384. | | | KX252492 | KX252494 | KX252490 | KX252493 | KX252491 | KX252488 | KX252489 |
| P. sp.38J5 taxon aquatilis[3] | Not formally described, American Type Culture Collection MYA-4577 as P. aquatilis. | Hong CX Richardson PA Hao W Ghimire SR Kong P et al. (2012) Phytophthora aquimorbida sp. nov. and Phytophthora taxon aquatilis recovered from irrigation reservoirs and a stream in Virginia USA. Mycologia 104: 10971108. | | | KX250933 | KX250935 | KX250931 | KX250934 | KX250932 | KX250929 | KX250930 |

| | | | | | | | | | | | |
|---|---|---|---|---|---|---|---|---|---|---|---|
| P. sp.40J5[3] | No formal description. | Yang, X., Tyler, B. M., & Hong, C. (2017). An expanded phylogeny for the genus Phytophthora. IMA fungus, 8(2), 355-384. | | | KX252499 | KX252501 | KX252497 | KX252500 | KX252498 | KX252495 | KX252496 |
| P. sp.46H5 ax[3] | No formal description. | Yang, X., Tyler, B. M., & Hong, C. (2017). An expanded phylogeny for the genus Phytophthora. IMA fungus, 8(2), 355-384. | | MH620072.1 | KX251850 | KX251852 | KX251848 | KX251851 | KX251849 | KX251846 | KX251847 |
| P. sp.48H2[3] | No formal description. | Yang, X., Tyler, B. M., & Hong, C. (2017). An expanded phylogeny for the genus Phytophthora. IMA fungus, 8(2), 355-384. | | | KX251484 | KX251486 | KX251482 | KX251485 | KX251483 | KX251480 | KX251481 |
| P. sp.56G1 pini like[3] | No formal description. | Yang, X., Tyler, B. M., & Hong, C. (2017). An expanded phylogeny for the genus Phytophthora. IMA fungus, 8(2), 355-384. | | | KX250856 | KX250858 | KX250854 | KX250857 | KX250855 | KX250852 | KX250853 |
| P. sp.61E7 taxon castitis[3] | No formal description. Unofficially described in Bertier et al. 2013. | Bertier L Brouwer H De Cock A Cooke DEL Olsson CHB et al. (2013) The expansion of Phytophthora clade 8b: three new species associated with winter grown vegetable crops. Persoonia 31: 6376 | DQ821186.1 | MH620086.1 | KX252102 | KX252104 | KX252100 | KX252103 | KX252101 | KX252098 | KX252099 |
| P. sp.61G1 taxon parsley[3] | No formal description. Unofficially described in Bertier et al. 2013. | Bertier L Brouwer H De Cock A Cooke DEL Olsson CHB et al. (2013) The expansion of Phytophthora clade 8b: three new species associated with winter grown vegetable crops. Persoonia 31: 6377 | MG865571.1 | MH620087.1 | KX252109 | KX252111 | KX252107 | KX252110 | KX252108 | KX252105 | KX252106 |
| P. sp.62C9[3] | No formal description. | Yang, X., Tyler, B. M., & Hong, C. (2017). An expanded phylogeny for the genus Phytophthora. IMA fungus, 8(2), 355-384. | | | KX251491 | KX251493 | KX251489 | KX251492 | KX251490 | KX251487 | KX251488 |
| P. sp.63H4 delaware[3] | No formal description. | Yang, X., Tyler, B. M., & Hong, C. (2017). An expanded phylogeny for the genus Phytophthora. IMA fungus, 8(2), 355-384. | | | KX251400 | KX251402 | KX251398 | KX251401 | KX251399 | KX251396 | KX251397 |
| P. sp.63H7 delaware[3] | No formal description. | Yang, X., Tyler, B. M., & Hong, C. (2017). An expanded phylogeny for the genus Phytophthora. IMA fungus, 8(2), 355-384. | | | KX251407 | KX251409 | KX251405 | KX251408 | KX251406 | KX251403 | KX251404 |
| P. stricta | None | Yang X; Copes WE; and Hong C. 2014. Two novel species representing a new clade and cluster of Phytophthora. Fungal Biology 118: 72-82. | MF138883.1 | MF138885.1 | KX252214, KX252221, KX252228, KX252235 | KX252216, KX252223, KX252230, KX252237 | KX252212, KX252219, KX252226, KX252233 | KX252215, KX252222, KX252229, KX252236 | KX252213, KX252220, KX252227, KX252234 | KX252210, KX252217, KX252224, KX252231 | KX252211, KX252218, KX252225, KX252232 |
| P. syringae | None | KlebahnH. 1909. Krankheiten des Flieders; pp. 75. Figs. 45. Berlin Gebruder Borntraeger. (Phytophthora syringae) | L41386.1 | HQ917884.1 | KX252200, KX252207 | KX252202, KX252209 | KX252198, KX252205 | KX252201, KX252208 | KX252199, KX252206 | KX252196, KX252203 | KX252197, KX252204 |
| P. tentaculata | None | KroberH, MarwitzR. 1993. Phytophthora tentaculata sp. nov. und Phytophthora cinnamomi var. parvispora var. nov. zwei neue Pilze von Deutschland. Z. Pflanzenk. Pflanzen. 100: 250-258. | MG761692.1 | AY564204.1 | EU079959, KX250450, KX250457 | EU079961, KX250452, KX250459 | EU079957, KX250448, KX250455 | EU079960, KX250451, KX250458 | EU079958, KX250449, KX250456 | EU079955, KX250446, KX250453 | EU079956, KX250447, KX250454 |
| P. terminalis | None | Man In 't Veld WA; Rosendahl KC; van Rijswick PC; Meffert JP; Westenberg M; van de Vossenberg BT; Denton G; and van Kuik FA. 2015. Phytophthora terminalis sp. nov. and Phytophthora occultans sp. nov.; two invasive pathogens of ornamental plants in Europe. Mycologia 107: 54-65. | MG865592.1 | JX978165.1 | KX250611 | KX250613 | KX250609 | KX250612 | KX250610 | KX250607 | KX250608 |
| P. theobromicola | None | Decloquement, J; Ramos-Sobrinho, R; Elias, SG; Britto, DS; Puig, AS; Reis, A; da Silva, RAF; Honorato-Júnior, J; Luz, EDMN; Pinho, DB; Marelli, J-P. 2021. Phytophthora theobromicola sp. nov.: a new species causing black pod disease on Cacao in Brazil. Frontiers in Microbiology. 12(no. 537399):1-15 | MT074263.1 | | MT074287.1 | | MT074279.1 | | | | MT074223.1 |
| P. thermophila | None | Jung T; Stukely MJC; Hardy GE St; White MD; Paap T; Dunstan WA; and Burgess TI. 2011. Multiple new Phytophthora species from ITS Clade 6 associated with natural ecosystems in Australia: evolutionary and ecological implications. Persoonia 13: 13-39. | MG696460.1 | MG931484.1 | KX251358 | KX251360 | KX251356 | KX251359 | KX251357 | KX251354 | KX251355 |
| P. trifolii | None | HansenEM andMaxwellDP. 1991. Species of the Phytophthora megasperma-complex. Mycologia 83: 376-381. | MG865595.1 | MH620080.1 | KX251955, KX251962 | KX251957, KX251964 | KX251953, KX251960 | KX251956, KX251963 | KX251954, KX251961 | KX251951, KX251958 | KX251952, KX251959 |
| P. tropicalis | Classified as P. palmivora by Mannon Gallegly. | Aragaki M andUchida JY. 2001. Morphological distinctions between Phytophthora capsici and P. tropicalis sp. nov. Mycologia 93: 137-145. | MG865596.1 | AY564161.1 | KX250695, KX250702 | KX250697, KX250704 | KX250693, KX250700 | KX250696, KX250703 | KX250694, KX250701 | KX250691, KX250698 | KX250692, KX250699 |
| P. tubulina | None | Jung T Horta Jung M Cacciola SO Cech T Bakonyi J Seress D Mosca S Schena L Seddaiu S Pane A Magnano di San Lio G Maia C Cravador A Franceschini A and Scanu B. 2017. Multiple new cryptic pathogenic Phytophthora species from Fagaceae forests in Austria Italy and Portugal. IMA Fungus 8 (2): 219244. | MH750723.1 | MF036279.1 | MF036255.1, MF036254.1, MF036253.1, MF036252.1, MF036251.1 | | | | | | MF036229.1, MF036228.1, MF036227.1, MF036226.1, MF036225.1 |
| P. tyrrhenica | None | Jung T Horta Jung M Cacciola SO Cech T Bakonyi J Seress D Mosca S Schena L Seddaiu S Pane A Magnano di San Lio G Maia C Cravador A Franceschini A andScanu B. 2017. Multiple new cryptic pathogenic Phytophthora species from Fagaceae forests in Austria Italy and Portugal. IMA Fungus 8 (2): 219244. | MT328712.1 | MF036286.1 | KU899423.1, KU899422.1, MF036260.1, MF036259.1, MF036258.1, MF036257.1, MF036256.1 | | | | | | KU899266.1, KU899265.1, MF036234.1, MF036233.1, MF036232.1, MF036231.1, MF036230.1 |
| P. uliginosa | None | JungT;HansenEM;WintonL;OsswaldW; andDelatourC.2002. Three new species of Phytophthora from European oak forests. Mycological Research 106: 397-411. | KJ755098.1 | KU681023.1 | KX251572, EU079696 | KX251573, EU079698 | EU080013, EU079694 | EU080015, EU079697 | KX251571, EU079695 | EU080011, EU079692 | EU080012, EU079693 |

| | | | | | | | | | | | |
|---|---|---|---|---|---|---|---|---|---|---|---|
| P. urerae | None | Grnwald N. J. Forbes G. A. PerezBarrera W. Stewart J. E. Fieland V. J. & Larsen M. M. (2018 December 26). Phytophthora urerae sp. nov. a new clade 1c relative of the Irish famine pathogen Phytophthora infestans from South America. Retrieved September 11 2020 from https://bsppjournals.onlinelibrary.wiley.com/doi/full/10.1111/ppa.12968 | KR632861.1 | KR632857.1 | | | | | | | KR632887.1, KR632888.1, KR632889.1 |
| P. versiformis | None | Paap, T., Croeser, L., White, D. et al. Phytophthora versiformis sp. nov., a new species from Australia related to P. quercina . Australasian Plant Pathol. 46, 369–378 (2017). https://doi.org/10.1007/s13313-017-0499-7 | | MH477760.1 | MK020401.1 | | MK864060.1 | | | OK533433.1 | MN207280.1 |
| P. vignae | None | PurssGS. 1957. Stem rot; a disease of cowpeas caused by an undescribed species of Phytophthora. Queensland J. Agric. Anim. Sci. 14: 125-154. | MK573644.1 | AY564205.1 | KX251780, KX251787, KX251794 | KX251782, KX251789, KX251796 | KX251778, KX251785, KX251792 | KX251781, KX251788, KX251795 | KX251779, KX251786, KX251793 | KX251776, KX251783, KX251790 | KX251777, KX251784, KX251791 |
| P. virginiana | None | Yang Xand Hong C. 2013. Phytophthora virginiana sp. nov.; a high-temperature tolerant species from irrigation water in Virginia. Mycotaxon 126: 167-176. | MT232849.1 | KC955181.1 | KX252368, KX252375, KX252382 | KX252370, KX252377, KX252384 | KX252366, KX252373, KX252380 | KX252369, KX252376, KX252383 | KX252367, KX252374, KX252381 | KX252364, KX252371, KX252378 | KX252365, KX252372, KX252379 |
| P. vulcanica | None | Jung T Horta Jung M Cacciola SO Cech T Bakonyi J Seress D Mosca S Schena L Seddaiu S Pane A Magnano di San Lio G Maia C Cravador A Franceschini A Scanu B. 2017. Multiple new cryptic pathogenic Phytophthora species from Fagaceae forests in Austria Italy and Portugal. IMA Fungus 8 (2): 219244. | MG603591.1 | MF036291.1 | MF036265.1, MF036264.1, MF036263.1, MF036262.1, MF036261.1 | | | | | | MF036239.1, MF036238.1, MF036237.1, MF036236.1, MF036235.1 |
| P. x cambivora[4] | Classified as P. cambivora by Mannon Gallegly. Renamed after it was determined that it is of hybrid origin. | Jung T Jung MH Scanu B Seress D Kovcs GM et al. (2017) Six new Phytophthora species from ITS Clade 7a including two sexually functional heterothallic hybrid species detected in natural ecosystems in Taiwan. Persoonia 38: 100135. | MH178329.1 | | KX251498, KX251505 | KX251500, KX251507 | KX251496, KX251503 | KX251499, KX251506 | KX251497, KX251504 | KX251494, KX251501 | KX251495, KX251502 |
| P. x heterohybrida[4] | None | Jung T Jung MH Scanu B Seress D Kovcs GM et al. (2017) Six new Phytophthora species from ITS Clade 7a including two sexually functional heterothallic hybrid species detected in natural ecosystems in Taiwan. Persoonia 38: 100135. | MN872721.1 | MN866071.1 | KX251641 | KX251643 | KX251639 | KX251642 | KX251640 | KX251637 | KX251638 |
| P. x incrassata[4] | None | Jung T Jung MH Scanu B Seress D Kovcs GM et al. (2017) Six new Phytophthora species from ITS Clade 7a including two sexually functional heterothallic hybrid species detected in natural ecosystems in Taiwan. Persoonia 38: 100135. | KU899206.1 | KU517150.1 | KX251648 | KX251650 | KX251646 | KX251649 | KX251647 | KX251644 | KX251645 |
| P. x pelgrandis[4] | None | Nirenberg, H.I.; Gerlach, W.F.; Gräfenhan, T. 2009. Phytophthora x pelgrandis, a new natural hybrid pathogenic to Pelargonium grandiflorum hort. Mycologia. 101(2):220-231 | MK496523.1 | MH136990.1 | | | MH359086.1 | | | MH380180.1 | MH494026.1 |
| P. x serendipita[4] | None | Man in 't Veld W.A., Rosendahl K.C.H.M., Hong C. 2012. Phytophthora X serendipita sp. nov. and P. X pelgrandis, two destructive pathogens generated by natural hybridization. Mycologia 104:1390-1396. | MG865599.1 | MH477762.1 | | | MH359087.1 | | | MH380181.1 | MH494027.1 |
| P. x stagnum[4] | None | Yang X. Richardson P.A Hong C.X. 2014. Phytophthora stagnum nothosp. nov. a new hybrid from irrigation reservoirs at ornamental plant nurseries in Virginia. PloS ONE. 9 (7):1-8 | MN306100.1 | KC631617.1 | KX251365, KX251372, KX251379, KX251386 | KX251367, KX251374, KX251381, KX251388 | KX251363, KX251370, KX251377, KX251384 | KX251366, KX251373, KX251380, KX251387 | KX251364, KX251371, KX251378, KX251385 | KX251361, KX251368, KX251375, KX251382 | KX251362, KX251369, KX251376, KX251383 |
| P. x vanyenensis[4] | None | Dang, QN; Pham, TQ; Arentz, F; Hardy, GEStJ; Burgess, TI. 2021. New Phytophthora species in clade 2a from the Asia-Pacific region including a re-examination of P. colocasiae and P. meadii. Mycological Progress. 20(2):111-129 | MT568651.1 | MT583648.1 | MT583680.1 | | OK267384.1 | | | OL342767.1 | MT583634.1 |
| Pp. vexans | [Outgroup] | Yang, X., Tyler, B. M., & Hong, C. (2017). An expanded phylogeny for the genus Phytophthora. IMA fungus, 8(2), 355-384. | MT533275.1 | KT692907.1 | EU080486 | EU080488 | EU080485 | EU080487 | | EU080483 | EU080484 |

Footnotes
1. Affinis. Isolate is considered to be distinct from this species, but to have affiinity to it.
2. G# indicates informal subgroup taxon belongs to

3. Not a formally described species. Referred to here with informal and/or isolate name.
4. Hybrid

Supplemental Table 2. Primers used to amplify the nine loci used in this study.

| Locus | Forward Primer Name | Forward Primer Sequence | Reverse Primer Name | Reverse Primer Sequence | Annealing Temperature | Source |
|---|---|---|---|---|---|---|
| 28S | LROR-O | 5’- ACCCGCTGAACTYAAGC-3’ | LR6-O | 5’-CGCCAGACGAGCTTACC-3’ | 53ºC | [1, 2] |
| 60SL10 | 60SL10_for | 5'-GCTAAGTGTTACCGTTTCCAG-3’ | 60SL10_rev | 5’- ACTTCTTGGAGCCCAGCAC-3’ | 53ºC | [3] |
| Btub | Btub_F1 | 5’-GCCAAGTTCTGGGAGGTCATC-3’ | Btub_R1 | 5’- CCTGGTACTGCTGGTACTCAG-3’ | 60ºC | [3, 4] |
| CoxI | COXF4N | 5’-GTATTTCTTCTTTATTAGGTGC-3’ | COXR4N | 5’-CGTGAACTAATGTTACATATAC-3’ | 52ºC | [4] |
| EF1a | EF1A_FL | 5’- GGTCACCTGATCTACAAGTGC-3’ | EF1A_RL | 5’- CCTTCTTGTTCACCGACTTG-3’ | 60ºC | [3] |
| ENL | Enl_for | 5’-CTTTGACTCGCGTGGCAAC-3’ | Enl_rev | 5’-CCTCCTCAATACGMAGAAGC-3’ | 60ºC | [3] |
| HS90 | HSP90_F1<br>HSP90_F2 | 5’-GCTGGACACGGACAAGAACC-3’<br>5’-ATGGACAACTGCGAGGAGC-3’ | HSP90_R1<br>HSP90_R2 | 5’-ACACCCTTGACRAACGACAG-3’<br>5’-CGTGTCGTACAGCAGCCAGA-3’ | 62ºC | [3] |
| ITS | ITS4 | 5’-TCCTCCGCTTATTGATATGC-3’ | ITS6 | 5’-GAAGGTGAAGTCGTAACAAGG-3’ | 55ºC | [5] |
| TigA | Tig_for<br>G3PDH_for | 5’-TTCGTGGGCGGYAACTGG-3’<br>5’-TCGCYATCAACGGMTTCGG-3’ | Tig_rev<br>G3PDH_rev | 5’-CCGAAKCCGTTGATRGCGA-3’<br>5’-GCCCCACTCRTTGTCRTACCAC-3’ | 64ºC | [3] |

Supplemental Table 3. Summary statistics for *Phytophthora infestans* SSR classifier by genotype class.

| Genotype | Number of Isolates | Sensitivity[a] | Specificity[b] | Prevalence[c] | Detection Rate[d] | Detection Prevalence[e] | Balanced Accuracy[f] |
|---|---|---|---|---|---|---|---|
| **North America** | | | | | | | |
| US_1 | 62 | 1.0000 | 1.0000 | 0.0407 | 0.0407 | 0.0407 | 1.0000 |
| US_6 | 2 | 1.0000 | 1.0000 | 0.0013 | 0.0013 | 0.0013 | 1.0000 |
| US_7 | 4 | 1.0000 | 0.9987 | 0.0026 | 0.0026 | 0.0039 | 0.9993 |
| US_8 | 7 | 0.8571 | 0.9980 | 0.0046 | 0.0039 | 0.0059 | 0.9276 |
| US_11 | 2 | 0.5000 | 0.9993 | 0.0013 | 0.0007 | 0.0013 | 0.7497 |
| US_13 | 1 | 0.0000 | 0.9993 | 0.0007 | 0.0000 | 0.0007 | 0.4997 |
| US_14 | 1 | 0.0000 | 1.0000 | 0.0007 | 0.0000 | 0.0000 | 0.5000 |
| US_15 | 1 | 0.0000 | 0.9993 | 0.0007 | 0.0000 | 0.0007 | 0.4997 |
| US_16 | 1 | 0.0000 | 0.9993 | 0.0007 | 0.0000 | 0.0007 | 0.4997 |
| US_17 | 1 | 0.0000 | 1.0000 | 0.0007 | 0.0000 | 0.0000 | 0.5000 |
| US_18 | 3 | 1.0000 | 1.0000 | 0.0020 | 0.0020 | 0.0020 | 1.0000 |
| US_20 | 5 | 1.0000 | 1.0000 | 0.0033 | 0.0033 | 0.0033 | 1.0000 |
| US_21 | 2 | 1.0000 | 1.0000 | 0.0013 | 0.0013 | 0.0013 | 1.0000 |
| US_23 | 5 | 0.6000 | 1.0000 | 0.0033 | 0.0020 | 0.0020 | 0.8000 |
| US_25 | 1 | 0.0000 | 1.0000 | 0.0007 | 0.0000 | 0.0000 | 0.5000 |
| **Central and South America** | | | | | | | |
| BR_1 | 5 | 1.0000 | 1.0000 | 0.0033 | 0.0033 | 0.0033 | 1.0000 |
| CR_1 | 18 | 0.9444 | 1.0000 | 0.0118 | 0.0111 | 0.0111 | 0.9722 |
| CR_2 | 3 | 0.3333 | 0.9993 | 0.0020 | 0.0007 | 0.0013 | 0.6663 |
| EC_1 | 786 | 1.0000 | 1.0000 | 0.5154 | 0.5154 | 0.5154 | 1.0000 |
| HN_1 | 12 | 0.7500 | 0.9974 | 0.0079 | 0.0059 | 0.0085 | 0.8737 |
| NI_1 | 22 | 1.0000 | 0.9993 | 0.0144 | 0.0144 | 0.0151 | 0.9997 |
| PE_3 | 81 | 1.0000 | 1.0000 | 0.0531 | 0.0531 | 0.0531 | 1.0000 |
| PE_7 | 40 | 0.9750 | 0.9993 | 0.0262 | 0.0256 | 0.0262 | 0.9872 |
| **Europe** | | | | | | | |
| EU_1_A1 | 2 | 1.0000 | 1.0000 | 0.0013 | 0.0013 | 0.0013 | 1.0000 |
| EU_2_A1 | 21 | 0.9524 | 0.9993 | 0.0138 | 0.0131 | 0.0138 | 0.9759 |
| EU_12_A1 | 1 | 0.0000 | 1.0000 | 0.0007 | 0.0000 | 0.0000 | 0.5000 |
| EU_13_A2 | 295 | 0.9864 | 0.9976 | 0.1934 | 0.1908 | 0.1928 | 0.9920 |
| EU_23_A1 | 95 | 1.0000 | 0.9986 | 0.0623 | 0.0623 | 0.0636 | 0.9993 |

| | | | | | | | |
|---|---|---|---|---|---|---|---|
| EU_36_A2 | 2 | 1.0000 | 0.9993 | 0.0013 | 0.0013 | 0.0020 | 0.9997 |
| EU_8_A1 | 6 | 1.0000 | 0.9993 | 0.0039 | 0.0039 | 0.0046 | 0.9997 |
| FAM_1 | 27 | 1.0000 | 1.0000 | 0.0177 | 0.0177 | 0.0177 | 1.0000 |
| **Asia** | | | | | | | |
| SIB_1 | 8 | 1.0000 | 1.0000 | 0.0052 | 0.0052 | 0.0052 | 1.0000 |
| CN_9 | 2 | 1.0000 | 1.0000 | 0.0013 | 0.0013 | 0.0013 | 1.0000 |
| CN_11 | 1 | 0.0000 | 1.0000 | 0.0007 | 0.0000 | 0.0000 | 0.5000 |

[a] true positive rate

[b] true negative rate

[c] the proportion of the dataset that is of this lineage

[d] the rate at which the lineage was detected

[e] the number of predicted positive lineages divided by the total number of lineages

[f] arithmetic mean of sensitivity and specificity

Supplemental Table 4. All genotypes of *Phytophthora infestans* included in the SSR classifier along with their SSR profiles and metadata.

| Genotype | Region | Country | Date | Host | D13 | PinfSSR8 | PinfSSR4 | Pi04 | Pi70 | PinfSSR6 | Pi63 | PiG11 | Pi02 | PinfSSR11 | PinfSSR2 | Pi4B |
|---|---|---|---|---|---|---|---|---|---|---|---|---|---|---|---|---|
| BR_1 | S_America | Bolivia | 1996 | S_tuberosum | 136/136/0 | 260/266/0 | 284/294/0 | 170/170/0 | 192/192/0 | 244/244/0 | 270/279/0 | 162/162/0 | 268/268/0 | 331/341/0 | 173/173/0 | 205/213/0 |
| CN_9 | Asia | China | 2001 | S_tuberosum | 136/140/0 | 264/266/0 | 292/294/0 | 166/170/0 | 192/192/0 | 242/244/0 | 279/279/0 | 160/162/0 | 268/268/0 | 331/355/0 | 173/173/0 | 213/217/0 |
| CN_10 | Asia | China | 2004 | S_tuberosum | 148/148/0 | 262/266/0 | 286/288/292 | 166/170/0 | 192/192/0 | 244/244/0 | 276/279/0 | 136/156/0 | 266/266/0 | 341/355/0 | 173/177/0 | 213/215/0 |
| CN_11 | Asia | China | 2002 | S_tuberosum | 110/110/0 | 264/264/0 | 284/296/302 | 166/170/0 | 192/192/0 | 244/244/0 | 279/279/0 | 134/154/208 | 258/266/268 | 331/341/0 | 173/173/0 | 213/213/0 |
| CR_1 | C_America | Costa_Rica | NA | NA | 162/164/0 | 260/266/0 | 290/292/0 | 166/170/0 | 192/192/0 | 244/244/0 | 273/279/0 | 156/156/0 | 266/268/0 | 341/355/0 | 173/173/0 | 213/213/0 |
| CR_2 | C_America | Costa_Rica | NA | NA | 108/112/0 | 260/260/0 | 284/288/294 | 166/170/0 | 192/192/0 | 244/244/0 | 273/279/0 | 156/160/0 | 266/266/0 | 341/355/0 | 173/173/0 | 213/225/0 |
| EC_1 | S_America | Ecuador | 1997 | S_phureja | 136/136/0 | 264/266/0 | 284/292/294 | 166/170/0 | 192/192/0 | 242/244/0 | 279/279/0 | 156/156/0 | 268/268/0 | 331/355/0 | 173/175/0 | 205/213/217 |
| EU_1_A1 | NA | NA | NA | NA | 136/136/0 | 260/266/0 | 284/294/0 | 166/170/0 | 192/192/0 | 244/244/0 | 270/279/0 | 142/162/0 | 266/268/0 | 341/355/0 | 173/175/0 | 213/217/0 |
| EU_2_A1 | NA | NA | NA | NA | 136/136/0 | 260/264/266 | 288/290/292 | 166/170/0 | 192/195/0 | 242/244/0 | 270/273/279 | 154/156/206 | 258/268/0 | 341/355/0 | 173/173/0 | 217/217/0 |
| EU_3_A2 | NA | NA | 2005 | NA | 118/136/0 | 260/266/0 | 284/288/294 | 166/170/0 | 192/195/0 | 242/244/0 | 270/279/0 | 154/160/0 | 268/268/0 | 331/341/0 | 173/173/0 | 213/217/0 |
| EU_5_A1 | N_America | US | 1950 | S_tuberosum | 136/136/0 | 266/266/0 | 292/294/0 | 166/170/0 | 192/192/0 | 244/244/0 | 279/279/0 | 156/162/0 | 268/268/0 | 341/355/0 | 175/175/0 | 205/217/0 |
| EU_6_A1 | NA | NA | NA | NA | NA | 260/266/0 | 284/292/296 | 166/170/0 | 192/195/0 | 244/244/0 | 273/279/0 | 160/160/0 | 266/268/0 | 341/355/0 | 173/175/0 | 213/217/0 |
| EU_8_A1 | NA | NA | NA | NA | 118/136/0 | 260/266/0 | 288/294/0 | 166/170/0 | 192/192/0 | 242/244/0 | 273/279/0 | 160/160/0 | 268/268/0 | 331/341/0 | 173/175/0 | 205/217/0 |
| EU_10_A2 | NA | NA | 2005 | NA | 136/136/0 | 260/266/0 | 284/288/0 | 166/170/0 | 192/192/0 | 240/242/244 | 273/279/0 | 162/208/0 | 268/268/0 | 331/341/0 | 173/175/0 | 213/217/0 |
| EU_13_A2 | Europe | UK | NA | NA | 136/140/154 | 260/266/0 | 284/294/0 | 166/170/0 | 192/192/0 | 240/244/0 | 273/279/0 | 154/160/0 | 266/268/0 | 341/341/0 | 173/173/0 | 205/213/0 |
| EU_17_A2 | NA | NA | NA | NA | 118/118/0 | 266/266/0 | 294/294/0 | 160/168/0 | 192/192/0 | 244/244/0 | 270/270/0 | 160/162/0 | 266/268/0 | 341/341/0 | 173/175/0 | 213/217/0 |
| EU_23_A1 | NA | NA | NA | NA | 136/210/0 | 260/266/0 | 288/294/0 | 170/170/0 | 192/192/0 | 244/244/0 | 270/279/0 | 142/156/0 | 266/268/270 | 331/341/0 | 173/175/0 | 213/217/0 |
| EU_33_A2 | NA | NA | NA | NA | 118/118/0 | 260/266/0 | 284/284/0 | 170/170/0 | 192/192/0 | 242/244/0 | 279/279/0 | 158/206/0 | 258/266/0 | 341/341/0 | 173/175/0 | 205/213/0 |
| EU_34_A1 | NA | NA | NA | NA | NA | 260/266/0 | 288/292/294 | 166/170/0 | 192/192/0 | 240/242/244 | 270/273/279 | 156/162/0 | 268/268/0 | 341/341/0 | 173/173/0 | 205/217/0 |
| EU_35_A2 | NA | NA | 2018 | S_tuberosum | 136/152/154 | 260/266/0 | 284/288/0 | 168/168/0 | 192/195/0 | 244/244/0 | 270/279/0 | 142/156/160 | 266/268/0 | 331/341/0 | 173/175/0 | 205/213/0 |
| EU_36_A2 | NA | NA | 2014 | NA | 136/152/156 | 260/266/0 | 288/294/0 | 166/170/0 | 192/195/0 | 240/242/244 | 270/279/0 | 150/154/160 | 268/268/0 | 341/341/0 | 173/175/0 | 205/213/217 |
| EU_37_A2 | NA | NA | NA | NA | NA | 260/266/0 | 284/298/0 | 168/168/0 | 192/192/0 | 242/244/0 | 279/279/0 | 148/154/0 | 268/268/0 | 341/355/0 | 173/173/0 | 213/217/0 |
| EU_38_A2 | Europe | France | NA | NA | 138/138/0 | 266/266/0 | 288/306/0 | 166/170/0 | 192/192/0 | 242/244/0 | 279/279/0 | 142/154/0 | 266/268/0 | 331/341/0 | 173/175/0 | 213/217/0 |
| EU_39_A1 | NA | NA | NA | NA | 136/154/0 | 260/260/0 | 288/294/0 | 168/168/0 | 192/192/0 | 240/244/0 | 279/279/0 | 142/206/0 | 268/268/0 | 341/341/0 | 173/175/0 | 205/213/0 |
| EU_41_A2 | Europe | Denmark | NA | NA | 136/136/0 | 260/266/0 | 284/288/294 | 168/168/0 | 192/192/0 | 242/244/0 | 279/279/0 | 156/206/0 | 266/268/0 | 341/341/0 | 173/175/0 | 213/213/0 |
| FAM_1 | Africa | Tanzania | 1942 | S_tuberosum | 108/136/138 | 266/266/0 | 288/298/0 | 166/170/0 | 192/192/0 | 244/244/0 | 270/273/0 | 160/200/0 | 266/266/0 | 341/355/0 | 173/173/0 | 209/213/0 |
| HN_1 | C_America | Honduras | 2014 | NA | 136/136/0 | 260/260/0 | 284/294/0 | 166/170/0 | 192/192/0 | 240/244/0 | 273/279/0 | 154/160/0 | 266/266/0 | 341/341/0 | 173/173/0 | 205/213/0 |
| MO_6 | Asia | China | 2004 | S_tuberosum | 104/142/0 | 266/266/0 | 288/292/0 | 166/170/0 | 192/192/0 | 244/244/0 | 273/279/0 | 160/208/0 | 268/268/0 | 331/341/355 | 173/173/0 | 205/213/217 |
| NI_1 | C_America | Nicaragua | NA | NA | 110/110/0 | 264/264/0 | 284/298/0 | 166/170/0 | 192/192/0 | 242/244/0 | 279/279/0 | 156/156/0 | 266/268/0 | 331/341/355 | 173/173/0 | 213/213/0 |
| PE_3 | S_America | Peru | 1997 | S_tuberosum | 134/134/0 | 266/266/0 | 288/292/0 | 166/170/0 | 192/195/0 | 244/244/0 | 279/279/0 | 160/160/0 | 268/268/0 | 341/355/0 | 175/175/0 | 205/217/0 |
| PE_7 | S_America | Peru | 1999 | S_peruvianur | 118/136/154 | 260/266/0 | 288/294/0 | 168/168/0 | 192/192/0 | 244/244/0 | 279/279/0 | 142/148/0 | 266/268/0 | 331/341/355 | 173/175/0 | 213/217/0 |
| SIB_1 | Asia | China | 2001 | S_tuberosum | 108/108/0 | 266/266/0 | 284/288/0 | 166/170/0 | 192/192/0 | 244/244/0 | 273/279/0 | 156/156/0 | 268/268/0 | 341/355/0 | 173/173/0 | 205/217/0 |
| US_1 | NA | NA | NA | NA | 138/140/0 | 266/266/0 | 288/290/300 | 166/170/0 | 189/192/0 | 244/262/0 | 270/273/279 | 152/156/200 | 266/266/0 | 341/355/0 | 173/177/0 | 213/217/0 |
| US_6 | N_America | US | 1992 | S_tuberosum | 144/144/0 | 264/264/0 | 292/294/0 | 166/170/0 | 192/192/0 | 244/256/0 | 279/279/0 | 156/206/0 | 258/266/0 | 341/355/0 | 173/173/0 | 213/217/0 |
| US_7 | N_America | Canada | 1993 | S_tuberosum | 110/110/0 | 266/266/0 | 296/302/0 | 166/170/0 | 192/192/0 | 244/244/0 | 279/279/0 | 148/156/0 | 266/270/0 | 341/341/0 | 173/173/0 | 213/213/0 |
| US_8 | N_America | US | 1994 | S_tuberosum | 108/112/0 | 260/266/0 | 284/288/294 | 166/170/0 | 192/192/0 | 244/244/0 | 279/279/0 | 156/156/0 | 266/268/0 | 341/355/0 | 173/173/0 | 213/225/0 |
| US_11 | N_America | Canada | 1995 | S_tuberosum | 110/110/0 | 264/266/0 | 284/294/302 | 166/170/0 | 192/192/0 | 244/244/0 | 279/279/0 | 134/156/0 | 258/266/268 | 331/341/0 | 173/173/0 | 213/213/0 |
| US_12 | N_America | US | 1994 | S_lycopersicu | 144/144/0 | 264/266/0 | 284/294/0 | 166/170/0 | 192/192/0 | 244/256/0 | 279/279/0 | 134/156/0 | 258/266/0 | 331/355/0 | 173/173/0 | 213/217/0 |
| US_13 | N_America | US | 1994 | Unknown | 110/110/0 | 266/266/0 | 294/304/0 | 166/170/0 | 192/192/0 | 244/244/0 | 279/279/0 | 134/156/0 | 266/266/0 | 341/355/0 | 173/173/0 | 213/213/0 |
| US_14 | N_America | US | 1994 | S_tuberosum | 108/112/0 | 260/266/0 | 284/288/294 | 166/170/0 | 192/192/0 | 244/244/0 | 279/279/0 | 152/156/0 | 266/268/0 | 341/355/0 | 173/173/0 | 213/225/0 |
| US_15 | N_America | US | 1994 | S_lycopersicu | 110/110/0 | 264/266/0 | 393/300/0 | 166/170/0 | 192/192/0 | 244/244/0 | 279/279/0 | 134/156/0 | 266/266/0 | 341/355/0 | 173/173/0 | 213/213/0 |
| US_16 | Asia | China | 2002 | S_tuberosum | 110/110/0 | 264/266/0 | 296/302/0 | 166/170/0 | 192/192/0 | 244/244/0 | 279/279/0 | 134/154/0 | 258/266/268 | 331/341/0 | 173/173/0 | 213/213/0 |
| US_17 | N_America | US | 1997 | S_lycopersicu | 110/110/0 | 264/266/0 | 284/294/0 | 166/170/0 | 192/192/0 | 244/244/0 | 279/279/0 | 134/156/0 | 258/268/0 | 331/341/0 | 173/173/0 | 213/213/0 |
| US_18 | N_America | US | 1996 | S_lycopersicu | 110/148/0 | 264/266/0 | 290/298/0 | 166/170/0 | 192/192/0 | 244/256/0 | 279/279/0 | 156/156/0 | 266/266/0 | 331/355/0 | 173/173/0 | 213/213/0 |
| US_19 | N_America | US | 1997 | S_lycopersicu | 110/110/0 | 260/264/0 | 288/298/0 | 166/170/0 | 192/192/0 | 244/244/0 | 279/279/0 | 134/154/0 | 266/266/0 | 355/355/0 | 173/173/0 | 213/213/0 |
| US_20 | N_America | US | 2004 | S_lycopersicu | 108/108/0 | 266/266/0 | 284/288/0 | 166/170/0 | 192/192/0 | 244/244/0 | 279/279/0 | 134/156/0 | 258/266/0 | 355/355/0 | 173/173/0 | 213/213/0 |

| | | | | | | | | | | | | | | | | |
|---|---|---|---|---|---|---|---|---|---|---|---|---|---|---|---|---|
| US_21 | N_America | US | 2006 | S_lycopersicu | 110/110/0 | 260/260/0 | 288/296/0 | 166/170/0 | 192/192/0 | 242/244/0 | 279/279/0 | 134/134/0 | 268/268/0 | 341/341/0 | 173/173/0 | 217/217/0 |
| US_23 | N_America | US | 2014 | S_lycopersicu | 134/210/0 | 260/266/0 | 288/294/296 | 170/170/0 | 192/192/0 | 244/244/0 | 270/279/0 | 142/156/206 | 266/268/270 | 331/341/0 | 173/175/0 | 213/217/0 |
| US_24 | N_America | US | 2009 | S_tuberosum | 108/108/0 | 260/266/0 | 284/288/298 | 166/170/0 | 192/195/0 | 244/244/0 | 279/279/0 | 156/156/0 | 268/268/0 | 341/355/0 | 173/173/0 | 217/225/0 |

T-BAS Public Tree Placement Instructions

1. Google “TBAS NCSU” and navigate to the [T-BAS webpage](T-BAS webpage).
2. To view the tree Select the “T-BAS Trees” option:

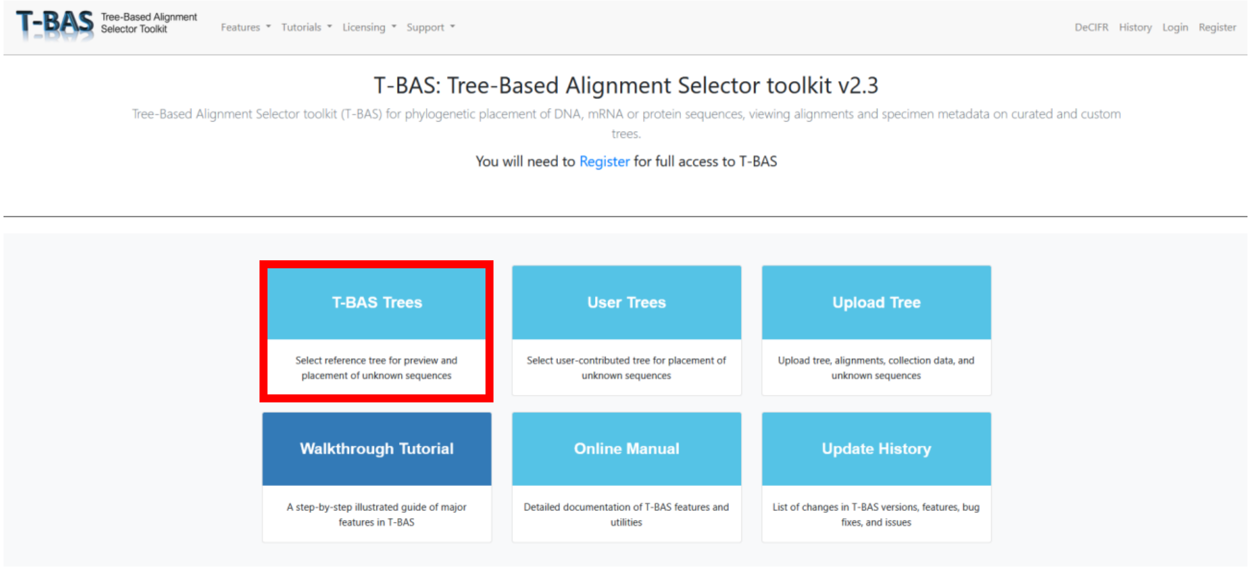

3. Select the **Phytophthora Nuc** Tree from the tree of life. This is the recommended tree for taxa placement. The Phytophthora CoxI tree contains only the mitochondrial CoxI locus. Look at this if you are interested, but it is not recommended for placement.

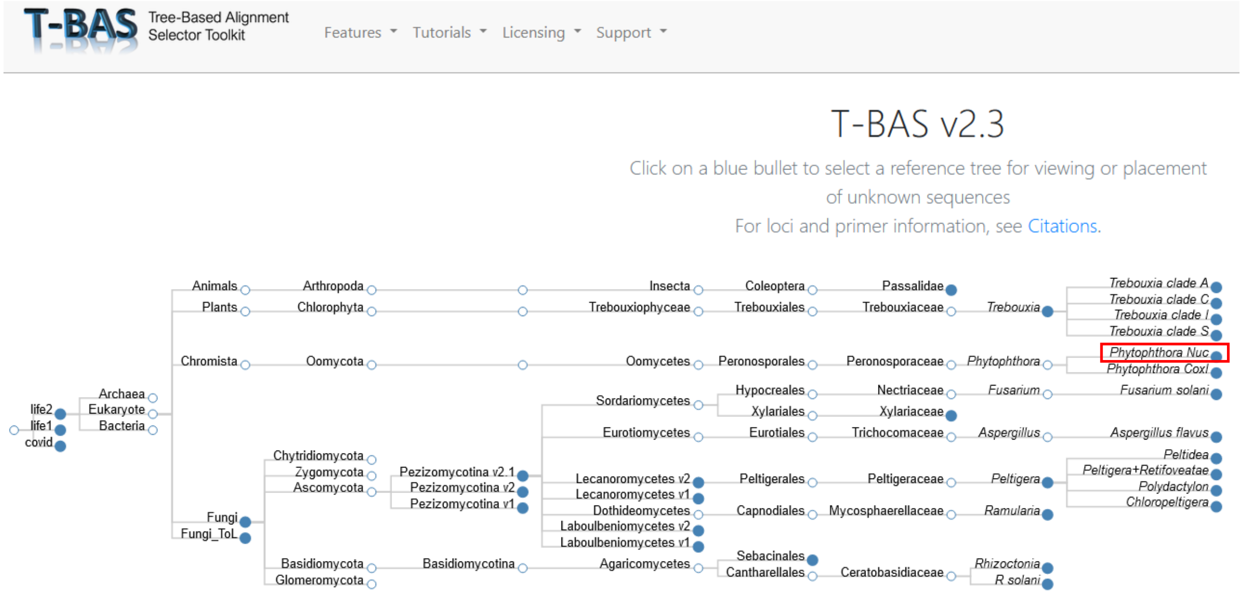

4. After selecting the tree, you should see two options. One for viewing the tree and one for placing taxa. Clicking view tree will open a new tab and load the tree, which takes a few moments. If you are interested in placing taxa, select “Place Unknowns”:

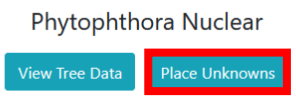

5. If you choose “Place Unknowns” a new screen will appear with several options. First, note the list of loci on the right (red box). These are the loci for which you can upload sequence data.

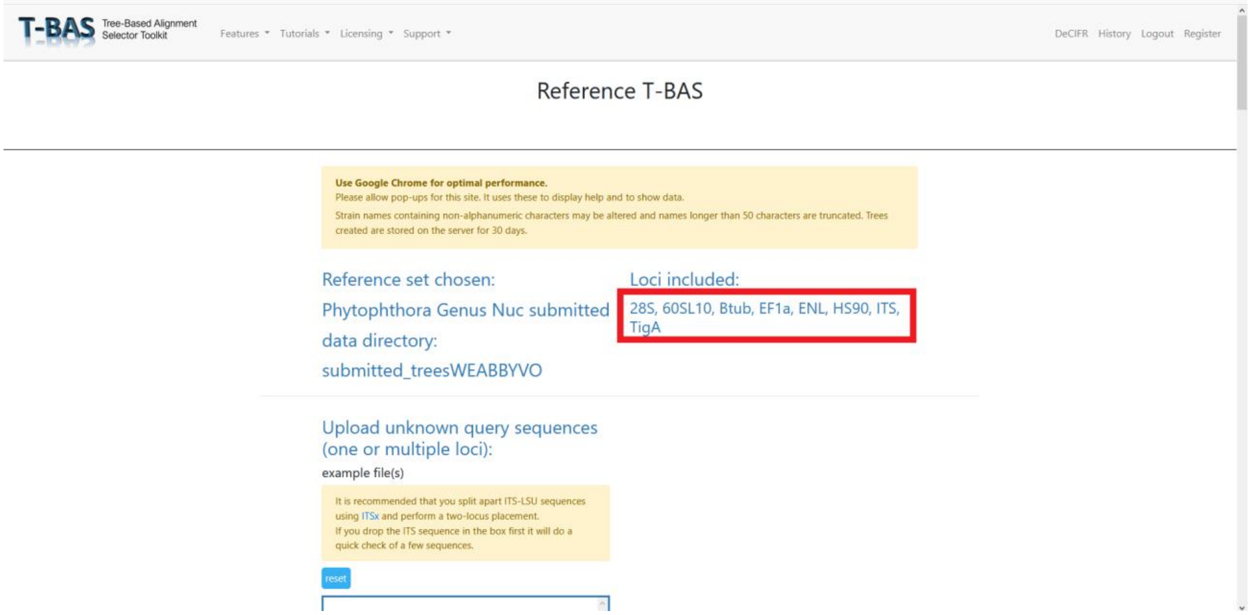

6. At the first input box, drag and drop sequence data in FASTA format for the taxa you would like to place. For this tutorial, example FASTA files have been provided in GitHub for each locus. They are named “locus_example.fasta” where ‘locus’ is the abbreviation for the relevant nuclear locus. Any or all of these files can be dragged and dropped into the Upload unknown query sequences box at this step.

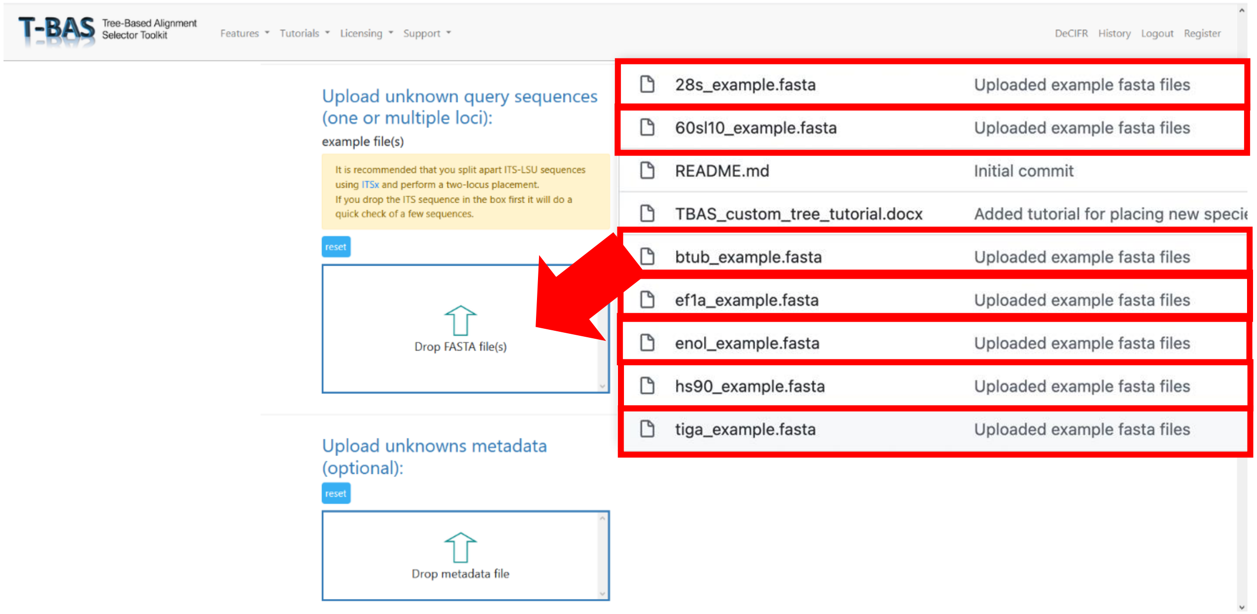

**Tips if you are uploading your own FASTA files:**

A separate FASTA file must be provided for each locus you are including.

If your FASTA files have multiple taxa, ensure the sequence headers for one taxon have the same name across files. For example, if you are uploading the HSP90 and TigA loci for two taxa named Phytophthora test1 and Phytophthora test2 your files should look like:

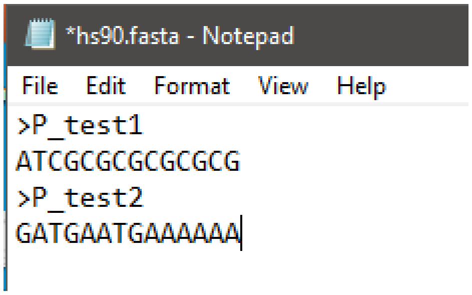

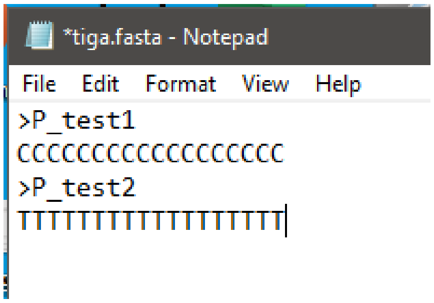

7. **Optional.** At the next dialog box, drag and drop a metadata file if you want to include one. If you do include a metadata file, change the drop-down menu below the dialog box from "Class" to "Species." Metadata must be a CSV file, where the first column is your taxa names. An example of how to format the metadata, including the required headers, is available at the associated GitHub page.

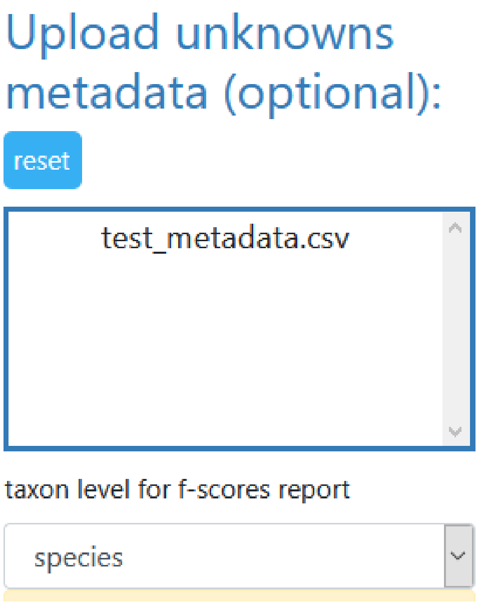

8. Scroll down and click "Submit." None of the other options need to be selected or changed. If you are interested in exploring phylogeny-based placement using a backbone constraint tree with bootstraps and other options, the TBAS Tutorials and Manual can help you navigate them.
9. After you click submit, a new webpage will appear where you can select which loci your FASTA files correspond to. Choose from the drop-down menu, then click submit again.

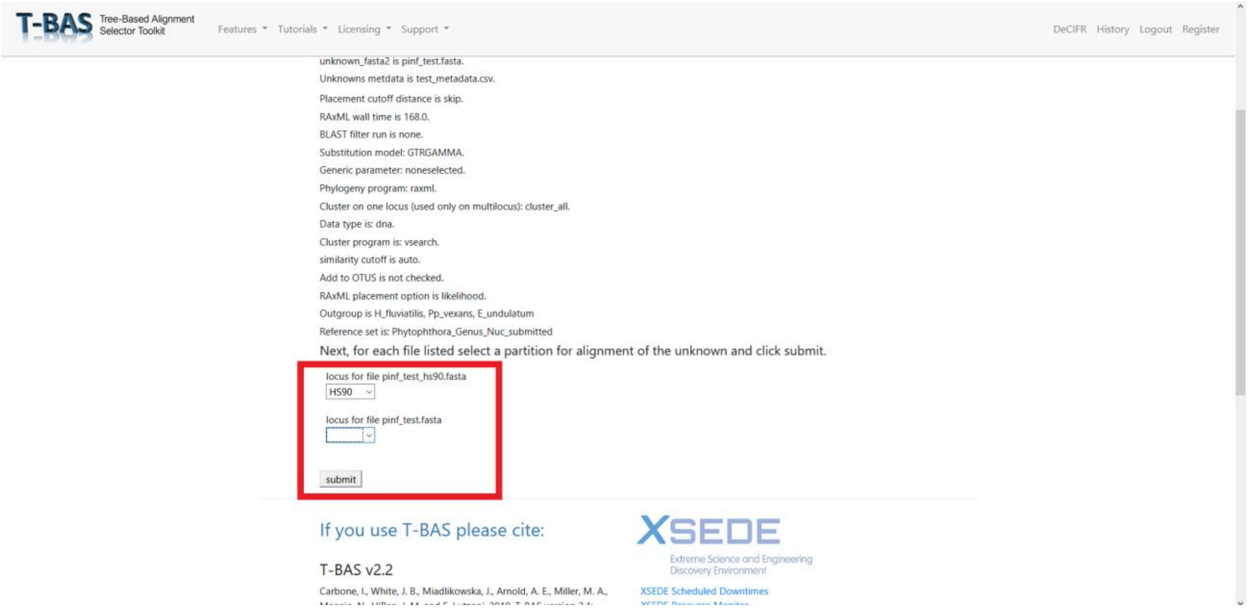

10. A loading bar will appear. Placement may take several minutes. When the tree placement is finished, several output files will be made available. Scroll down and click "View Tree." This will open the tree in a new tab.

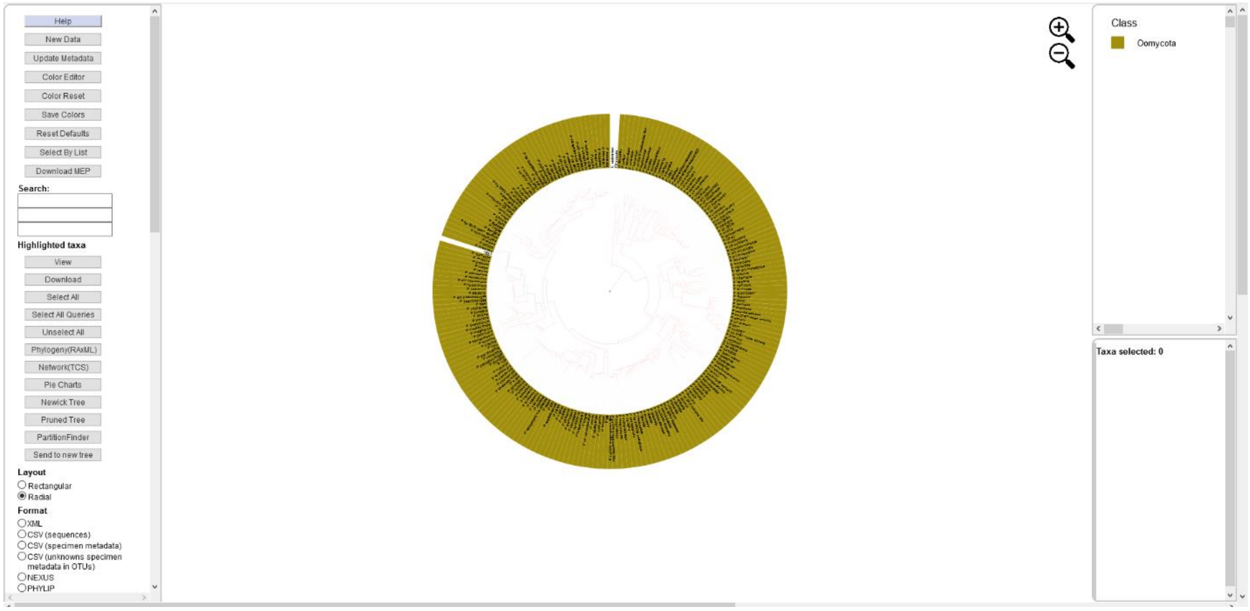

11. Select different options on the left as you choose to visualize the tree. We recommend colorizing leaves by different metadata, such as clade:

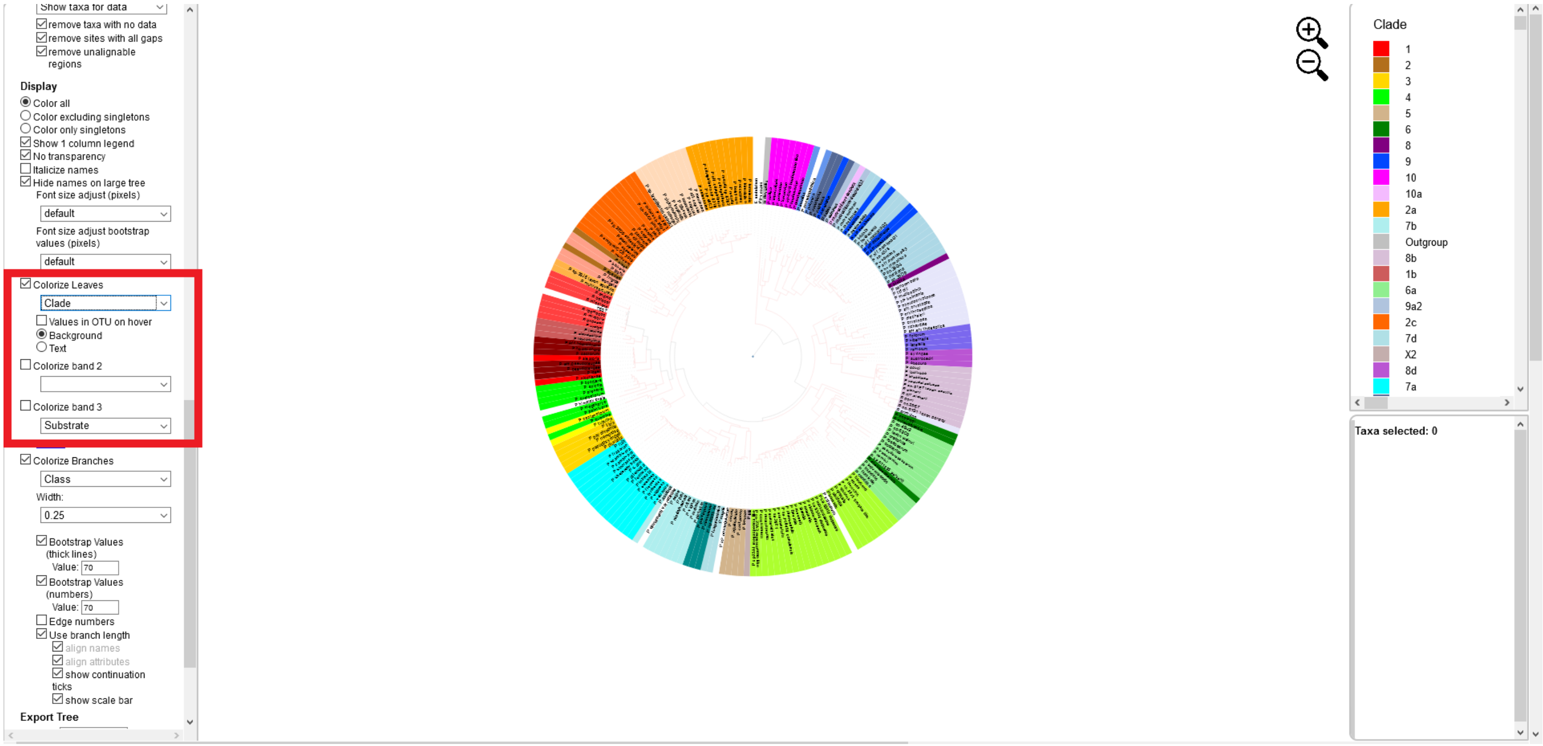
Show taxa for data
remove taxa with no data
remove sites with all gaps
remove unalignable regions
Display
Color all
Color excluding singletons
Color only singletons
Show 1 column legend
No transparency
Italicize names
Hide names on large tree
Font size adjust (pixels)
default
Font size adjust bootstrap values (pixels)
default
Colorize Leaves
Clade
Values in OTU on hover
Background
Text
Colorize band 2
Colorize band 3
Substrate
Colorize Branches
Class
Width:
0.25
Bootstrap Values (thick lines)
Value: 70
Bootstrap Values (numbers)
Value: 70
Edge numbers
Use branch length
align names
align attributes
show continuation ticks
show scale bar
Export Tree
Clade
1
2
3
4
5
6
8
9
10
10a
2a
7b
Outgroup
8b
1b
6a
9a2
2c
7d
X2
8d
7a
Taxa selected: 0

**Supplemental File 1. SSR Classifier Tutorial**

This program identifies unknown lineages of *Phytophthora infestans* based on 12 commonly used SSR markers (microsatellite data). The program is hosted on the USABlight.org website under "Identify SSR Genotype" and the database represents most of the currently named SSR lineages of *P. infestans*.

##### How it works

The SSR classifier compares the SSR data for your isolates of interest to over 2,500 described isolates from global populations in our *P infestans* database. The closest genetic match is determined using Bruvo's distance (described here).

##### How to run SSRs

This program uses results obtained from a 12-plex microsatellite amplification for *Phytophthora infestans*. These 12 microsatellite loci are well described for *Phytophthora infestans* and vary across lineages, allowing identification of different genotypes of the pathogen. Genotyping using this SSR method first involves determining the microsatellite alleles for each of the 12 loci. The methods for running SSRs have been previously described here. A simplified version of this protocol is attached in Appendix A.

##### How to format data

The raw microsatellite reads obtained from that methodology must then be processed and scored to create input data for the SSR classifier. This data will be available as a trace file (.fsa), which will contain colored peaks corresponding to lengths of different genetic fragments. The 12-plex protocol described at the link above uses five dyes and generates thirteen series of fragments in total. The 12 microsatellite loci are represented by four different dye colors, with fragment length between those with like colors dissimilar enough to allow for differentiation. There is also a ladder included, which will have its own color of dye. The raw reads visible in this trace file can be scored using several different bioinformatics programs, including Geneious, a popular paid program, and STRand, a free resource provided by UC Davis. Using one of these programs or something similar, the researcher identifies the peak (or peaks if heterozygous) of fluorescence for a particular color in a particular region. Using the tables below, this dye color and fragment length combination can be used to identify different alleles. Once the alleles are determined from the peak data, they can be formatted similar to the example CSV (see link or below) and uploaded to this tool for genotype identification. Three numbers are separated by forward slashes for each locus to represent all three alleles in triploid individuals. If only two alleles are present, the third number is a zero. There is one column for each locus, along with metadata columns for sample name, genotype, region, country, date, host, number of missing loci, and number of present loci. Each row beneath the header represents the data for one sample.

| Sample | Genotype | Region | Country | Date | Host | D13 | PinfSSR8 | PinfSSR4 | Pi04 | Pi70 | PinfSSR6 | Pi63 | PiG11 | Pi02 | PinfSSR11 | PinfSSR2 | Pi4B | Missing | Present |
|---|---|---|---|---|---|---|---|---|---|---|---|---|---|---|---|---|---|---|---|
| **EXAMPLE** | unknown | Europe | Poland | 2016 | unknown | 136/136/0 | 260/266/0 | 284/284/0 | 166/170/0 | 192/192/0 | 244/244/0 | 273/279/0 | 142/156/0 | 258/266/0 | 341/356/0 | 173/173/0 | 213/217/0 | 0 | 12 |

Figure 1. Example formatting of an input CSV for the SSR classifier.

**How to upload data**

Upload a CSV file containing metadata about your sample as well as SSR data for the 12 loci of interest. Follow the format in the example CSV. We recommend downloading this CSV, replacing the data with your own, and re-uploading it. The classifier will still work if your data is missing some loci, but at least 10 loci are recommended for the most accurate results. Include as many samples as you want.

**How to interpret results**

If your unknown sample has a close match, the program will return information about the match from our database of *Phytophthora infestans* genotypes (Figure 2). The tool first presents a table of input samples with their closest match and metadata associated with the closest match. For example, in Figure 2, input sample EG_256 most closely matches an isolate of lineage EU_23_A1 found in Egypt. After this table follows a more in-depth description of each match and where to learn more about the matching genotype. Finally, a list of all submitted samples for which no match was found are listed. Note just because it shows a match to an Egyptian genotype, this doesn't mean that was the source.  The region, country, sample date and host are just additional metadata you can use in downstream analysis.

Results

| Input Sample | Matching Genotype | Region | Country | Date | Host | Bruvo's Distance |
|---|---|---|---|---|---|---|
| EG_256 | EU_23_A1 | Africa | Egypt | 2012/2013 | S_lycopersicum | 0.00 |
| EG_152 | EU_23_A1 | Africa | Egypt | 2013/2014 | S_lycopersicum | 0.07 |
| BU-11.22 | Unknown | Europe | UK | 2013 | S_tuberosum | 0.03 |
| BU-13.P04 | Unknown | Europe | UK | 2013 | S_tuberosum | 0.00 |
| BU-13.P10 | Unknown | Europe | UK | 2013 | S_tuberosum | 0.00 |

| Input Sample | Description |
|---|---|
| EG_256 | Based on the markers you provided, the genotype of this isolate is most likely EU_23_A1 or US_23. These two lineages are synonymous. In the eastern United States, US_23 has been found on both potato and tomato and has mating type A1. For more information, see https://doi.org/10.1094/PDIS-03-11-0156-RE. |
| EG_152 | Based on the markers you provided, the genotype of this isolate is most likely EU_23_A1 or US_23. These two lineages are synonymous. In the eastern United States, US_23 has been found on both potato and tomato and has mating type A1. For more information, see https://doi.org/10.1094/PDIS-03-11-0156-RE. |
| BU-11.22 | The markers you provided indicate your isolate is closely related to a sample in our database of Unknown genotype. This means your sample could be a clone that has been found but not described, or it could be from a sexually recombining population. Unknown. |
| BU-13.P04 | The markers you provided indicate your isolate is closely related to a sample in our database of Unknown genotype. This means your sample could be a clone that has been found but not described, or it could be from a sexually recombining population. Unknown. |
| BU-13.P10 | The markers you provided indicate your isolate is closely related to a sample in our database of Unknown genotype. This means your sample could be a clone that has been found but not described, or it could be from a sexually recombining population. Unknown. |

```
No close matches found for sample TU0899 .
No close matches found for sample TU0761 .
No close matches found for sample TU0803 .
No close matches found for sample TU0750 .
```

**Figure 2**. Example output from SSR classifier.

**Having issues?**

**Other Resources**

USABlight: Genotype tracking website for North America
EuroBlight: Genotype tracking website for Europe
Tizón Latino: Genotype tracking website for Latin America

Help notes for scoring SSR alleles in *Phytophthora infestans*: Allele calling guide developed by David Cooke.

For information on the SSR binning scheme used by this tool, see "Population structure of *Phytophthora infestans* collected on potato and tomato in Italy" by Saville et al. 2021, Supplemental Table 3.

### Appendix A

#### 12-plex Microsatellite (SSR) Genotyping of *Phytophthora infestans*

Microsatellites can be used for genotyping lineages of *P. infestans*. Li and Cooke (2013) have developed a protocol that multiplexes 12 diagnostic SSR primer sets in a single tube for more rapid analysis and genotyping. The protocol uses fluorescently labeled primers, which can then be read by a capillary analyzer for analysis. This protocol is optimized for use with an ABI 3730xl DNA analyzer with a 5 dye set (6-FAM, VIC, NED, PET, and LIZ size standard). The following protocol is from Li and Cooke, with modifications implemented by the lab of Bill Fry at Cornell University and Anne Njoroge.

**Primers (5' – 3')**

| Locus | Dye | Product size range (bp) | Primer sequence |
|---|---|---|---|
| PiG11 | NED | 130-206 | FwdNED-TGCTATTTATCAAGCGTGGG<br>Rev-GTTTCAATCTGCAGCCGTAAGA |
| Pi02 | NED | 255-275 | FwdNED-ACTTGCAGAACTACCGCCC<br>Rev-GTTTGACCACTTTCCTCGGTTC |
| PinfSSR11 | NED | 325-360 | FwdNED-TTAAGCCACGACATGAGCTG<br>Rev-GTTTAGACAATTGTTTTGTGGTCGC |
| D13 | FAM | 100-210 | FwdFAM-TGCCCCCTGCTCACTC<br>Rev-GCTCGAATTCATTTTACAGACTTG |
| PinfSSR8 | FAM | 250-275 | FwdFAM-AATCTGATCGCAACTGAGGG<br>Rev-GTTTACAAGATACACACGTCGCTCC |
| PinfSSR4 | FAM | 280-305 | FwdFAM-TCTTGTTCGAGTATGCGACG<br>Rev-GTTTCACTTCGGGAGAAAGGCTTC |
| Pi04 | VIC | 160-175 | FwdVIC –AGCGGCTTACCGATGG<br>Rev-GTTTCAGCGGCTGTTTCGAC |
| Pi70 | VIC | 185-205 | FwdVIC – ATGAAAATACGTCAATGCTCG<br>Rev-CGTTGGATATTTCTATTTCTTCG |
| PinfSSR6 | VIC | 230-250 | FwdVIC-GTTTTGGTGGGGCTGAAGTTTT<br>Rev - TCGCCACAAGATTTATTCCG |
| Pi63 | VIC | 265-280 | FwdVIC – ATGACGAAGATGAAAGTGAGG<br>Rev-CGTATTTTCCTGTTTATCTAACACC |
| PinfSSR2 | PET | 165-180 | FwdPET-CGACTTCTACATCAACCGGC<br>Rev-GTTTGCTTGGACTGCGTCTTTAGC |
| Pi4B | PET | 200-295 | FwdPET – AAAATAAAGCCTTTGGTTCA<br>Rev-GCAAGCGAGGTTTGTAGATT |

**Reference:** Li, Y.; Cooke, D.E.L.; Jacobsen, E.; van der Lee, T. 2013. Efficient multiplex simple sequence repeat genotyping of the oomycete plant pathogen *Phytophthora infestans*. Journal of Microbiological Methods 92: 316-322.

#### Appendix A (cont.)

Instead of individually pipetting each primer into the master mix, a 10X multiplex primer mix is made that includes all primers. The primer mix is made as follows (makes 400µl):

| Primer | Volume of 100µM primer stock (µl) |
|---|---|
| PiG11F | 6 |
| PiG11R | 6 |
| Pi02F | 6 |
| Pi02R | 6 |
| PinfSSR11F | 6 |
| PinfSSR11R | 6 |
| PinfSSR4F | 6 |
| PinfSSR4R | 6 |
| Pi04F | 6 |
| Pi04R | 6 |
| Pi70F | 6 |
| Pi70R | 6 |
| PinfSSR6F | 6 |
| PinfSSR6R | 6 |
| Pi63F | 6 |
| Pi63R | 6 |
| PinfSSR2F | 6 |
| PinfSSR2R | 6 |
| D13F | 6.4 |
| D13R | 6.4 |
| PinfSSR8F | 12 |
| PinfSSR8R | 12 |
| Pi4BF | 12 |
| Pi4BR | 12 |

Combine with 231.2µL of 10 mM Tris buffer (pH=8.0) to make 400µl of primer mix.

**Reference:** Njoroge, A.W. et al. 2019. Genotyping of *Phytophthora infestans* in eastern Africa reveals a dominating invasive European lineage. Phytopathology 109: 670-680.

### Appendix A (cont.)

The master mix can be made using either the Qiagen multiplex PCR kit (Qiagen, cat. No 206145) or the Qiagen Type-it Microsatellite PCR kit (Qiagen, cat. No. 206243). For the purposes of this protocol we use the Type-it microsatellite PCR kit.

| **Reagent** | **Volume per reaction (µl)** |
|---|---|
| 2X Type-it master mix | 6.25 |
| 10X multiplex primer mix | 1.3 |
| PCR grade water | 1.95 |
| Total reaction mix volume per sample (µl) | 11.5 |

1µl of template DNA is added to bring the total volume per sample to 12.5 µl. If more DNA is desired, adjust the volume of reaction mix per sample.

Thermocycling program:

| | | |
|---|---|---|
| 1 cycle | 95C | 5 min. |
| | 95C | 30 sec. |
| 33 cycles | 58C | 90 sec. |
| | 72C | 20 sec. |
| 1 cycle | 60C | 30 min. |

Before loading on a DNA analyzer, samples must be prepared with the LIZ size standard (Applied Biosystems LIZ500, cat. No. 4322682) and suspended in an appropriate loading solution. For use on an ABI 3730xl DNA analyzer we use highly deionized formamide (hi di formamide, Applied Biosystems, cat. No. 4311320). Check with your local source for fragment analysis for preparation and submission protocols specific to their facilities.

Master mix for analysis preparation:

| Reagent | Volume per reaction (µl) |
|---|---|
| Hi-di formamide | 10 |
| LIZ500 size standard | 0.3 |
| Total volume per sample (µl) | 10.3 |

#### Appendix A (cont.)

Add 0.5 µl of template reaction product to bring the total volume per sample to 10.8 µl. More product may be used if desired, up to 3 µl.

An optional denaturation step can be employed after plate prep to increase peak resolution by heating the prepared plate at 95C for 3 min, then chilling on ice.
